# Design and analysis of the 4-anilino-quin(az)oline kinase inhibition profiles of GAK/SLK/STK10 using quantitative structure activity relationships

**DOI:** 10.1101/757047

**Authors:** Christopher R. M. Asquith, Tuomo Laitinen, James M. Bennett, Carrow I. Wells, Jonathan M. Elkins, William J. Zuercher, Graham Tizzard, Antti Poso

**Affiliations:** Department of Pharmacology, School of Medicine, University of North Carolina at Chapel Hill, Chapel Hill, NC 27599, USA; Structural Genomics Consortium, UNC Eshelman School of Pharmacy, University of North Carolina at Chapel Hill, Chapel Hill, NC 27599, USA; School of Pharmacy, Faculty of Health Sciences, University of Eastern Finland, 70211 Kuopio, Finland; Structural Genomics Consortium, Nuffield Department of Clinical Medicine, University of Oxford, Old Road Campus Research Building, Oxford, OX3 7DQ, United Kingdom; Structural Genomics Consortium, Departamento de Genetica e Evolução, Instituto de Biologia, UNICAMP, Campinas, SP 13083-886, Brazil; Lineberger Comprehensive Cancer Center, University of North Carolina at Chapel Hill, NC 27599, USA; UK National Crystallography Service, School of Chemistry, University of Southampton, Highfield Campus, Southampton, SO17 1BJ, United Kingdom

## Abstract

The 4-anilino-quinoline and 4-anilino-quinazoline ring systems have been the focus of significant efforts in prior kinase drug discovery programs, which have led to approved medicines. Broad kinome profiles of these compounds have now been assessed with the advent of advanced screening technologies. These ring systems while, originally designed for specific targets including epidermal growth factor receptor (EGFR), actually display a number of potent collateral kinase targets, some of which have been associated with negative clinical outcomes. We have designed and synthesized a series of 4-anilino-quin(az)olines in order to better understand the structure activity relationships of three main collateral kinase targets of quin(az)oline-based kinase inhibitors: cyclin G associated kinase (GAK), STE20-like serine/threonine-protein kinase (SLK) and Serine/threonine-protein kinase 10 (STK10) through a series of quantitative structure activity relationship (QSAR) analysis and water mapping of the kinase ATP binding sites.

## INTRODUCTION

Kinases have been successfully utilized as drug targets for the past 30 years, with 47 kinase inhibitors approved by the FDA to date. These drugs are predominantly multi-targeted tyrosine kinase inhibitors for the treatment of cancer.^1, 2^ A large number of additional inhibitors are currently being progressed through preclinical and clinical development,^2^ but with over 500 kinases in the human genome,^3^ there remains significant untapped potential within the kinome for the treatment a wide range of human disease.^4^

To advance the understanding of kinome structure activity relationships, large scale kinome-wide profiling of ATP-competitive kinase inhibitors have started to uncover the preferred chemotypes for the inhibition of both well studied and under studied kinases.^5–8^ The first-and second-generation GlaxoSmithKline Published Kinase Inhibitor Sets (PKIS and PKIS2) are collections of well characterized ATP-competitive kinase inhibitors with broad coverage of the human kinome.^9–10^ Together these screens have provided a more clear view of the kinome inhibitor landscape and the promiscuously profiles of these inhibitors. These kinome-wide profiling efforts have also started to uncover the preferred chemotypes for the inhibition of different families or sub-families of kinases.^5–10^

The 4-anilino-quin(az)oline is a common kinase inhibitor chemotype.^5–10^ Several quinazoline-based clinical kinase inhibitors show cross reactivity with cyclin G associated kinase (GAK), STE20-like serine/threonine-protein kinase (SLK) and Serine/threonine-protein kinase 10 (STK10), including the approved drugs gefitinib, vandetanib, lapatinib, neratinib, afatinib, erlotinib, and other candidate quin(az)olines including pelitinib and dacomitinib (Figure 1).^2, 6–7^ These drugs were originally designed as inhibitors of either epidermal growth factor receptor (EGFR) or Src family kinases, and all show GAK, SLK, and STK10 as common collateral targets (Figure 1).^2, 6–7^ GAK activity in this chemotype has already been observed in the erlotinib quinoline derivative (**1**)^11^ and a selective chemical probe for GAK, SGC-GAK-1 (**2**).^12^ This probe was developed from a series of quinolines exemplified by **3** (Figure 1).^13^ The SLK/STK10 activity is less consistent, but several examples display potent profiles with erlotinib demonstrating single digit nanomolar potency on both SLK and STK10 (Figure 1).^6–7^

**Figure 1.**
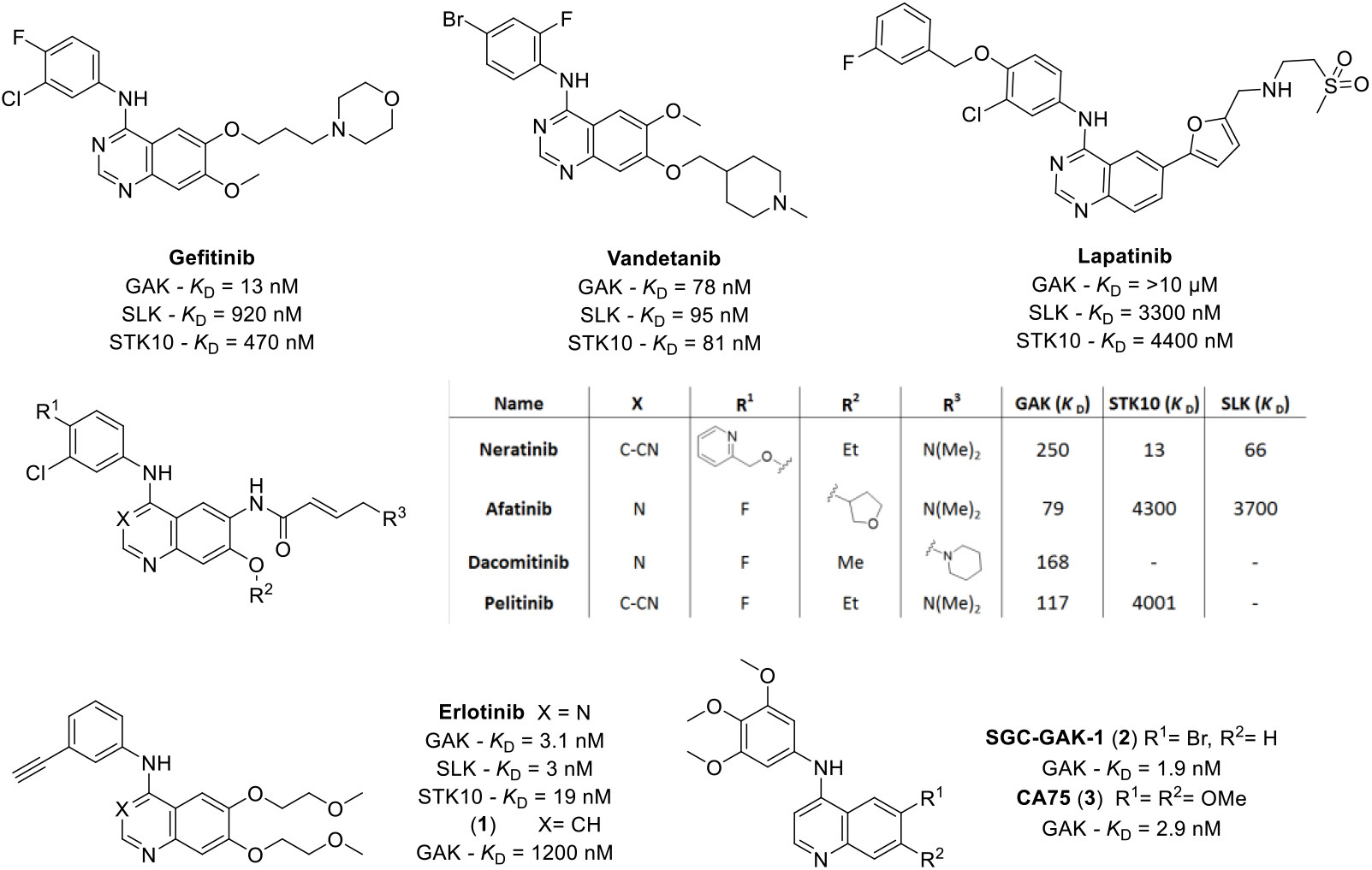
Previously reported EGFR 4-anilinoquin(az)olines with GAK/SLK/STK10 activity

GAK is a 160 kDa serine/threonine kinase originally identified as a direct association partner of cyclin G and a member of the numb-associated kinase (NAK) family.^3, 14–15^ In addition to its kinase domain, the GAK *C*-terminus has high homology with auxilin and tensin, with broad expression and localization to the Golgi complex, cytoplasm, and nucleus.^16–17^ GAK has been implicated in diverse biological processes, and genome-wide association studies have identified single nucleotide polymorphisms in the GAK gene associated with susceptibility to Parkinson’s disease.^18^ GAK is also involved in membrane trafficking and uncoating of clathrin-coated vesicles in the cytoplasm,^19–20^ and is required for maintenance of centrosome maturation and progression through mitosis.^21^ GAK is over-expressed in osteosarcoma, and prostate cancer where GAK is implicated in progression to androgen independence.^12, 22^ GAK inhibition has been associated with clinical toxicity due to pulmonary alveolar dysfunction based upon the phenotype of transgenic mice expressing catalytically inactive GAK, but this hypothesis has not yet been tested with a selective small molecule GAK inhibitor.^23–24^

STK10/SLK are members of the Ste20 family of serine/threonine protein kinase sub-family. SLK has just one amino acid difference in the ATP active site of STK10.^3, 25–26^ The expression and activity SLK is increased during kidney development and recovery from ischemia-reperfusion injury.^27–29^ SLK mediates apoptosis in various cells, and can regulate cell cycle progression and cytoskeletal remodelling.^30^ STK10 is closely connected with several polo-like kinase (PLK) kinases.^3, 26^ The protein can associate with and phosphorylate polo-like kinase 1, and over-expression of a kinase-dead version of the protein interferes with normal cell cycle progression.^26^ The kinase can also negatively regulate interleukin 2 expression in T-cells *via* the mitogen activated protein kinase kinase 1 pathway.^31^

STK10 is also known as lymphocyte-oriented kinase (LOK), is known to prevent ezrin, radixin, and moesin (ERM) phosphorylation during apoptosis that play a crucial role linking the plasma membrane to actin filaments.^32^ In lymphocytes, ERM phosphorylation contributes to cell rigidity and to the maintenance of microvilli.^33^ This causes restricted formation of microvilli at the apical surface and disrupted cell function.^34^ Furthermore, broad siRNA knock down screening of four cell lines representative of Ewing’s sarcoma, an aggressive musculoskeletal tumour have demonstrated that STK10 is a key kinase for tumour progression.^35^ STK10 was not only shown to have a key role in proliferation of the tumours, but siRNA knock down of STK10 showed increased apoptosis of the tumour cells.^36^

The 4-anilino-quin(az)oline shows differing kinome wide inhibition profiles in the literature. The primary EGFR target is often accompanied by a number of secondary kinases targets including GAK, STK10 and SLK.^5–10^ We wanted to explore the possibility of chemical probe development for SLK and STK10 in addition to GAK.^12^ We now describe through a series of QSAR and rational design a series of diverse explorations and kinase profiles on the 4-anilino-quin(az)oline scaffold.

## RESULTS

We first selected a set of diverse literature 4-anilinoquin(az)oline based kinase inhibitors that covered a cross section of chemical space (Figure 1). The compounds were screened for activity on the kinase domains of the NAK sub-family (GAK, AAK1, BMP2K, and STK16) and Ste20 family of serine/threonine protein kinase (STK10 and SLK) using a time-resolved fluorescence energy transfer (TR-FRET) binding displacement assay in a 16-point dose-response format to determine the inhibition constant (K_i_) (Table 1, **Table S1**).^13^ The compounds were then evaluated to assess their in-cell activity on GAK in a nanoBRET cellular target engagement assay in an eight-point dose–response format to determine the cellular binding potency (IC_50_). The nanoBRET evaluated the compounds ability to displace a fluorescent tracer molecule from the ATP binding site of a transiently expressed full length GAK protein *N*-terminally fused with NanoLuc luciferase (Nluc) in HEK293T cells. In the absence of compound, the tracer molecule and fusion protein were in proximity and able to generate an observable bioluminescence resonance energy transfer (BRET) signal.^37^

**Table 1.**
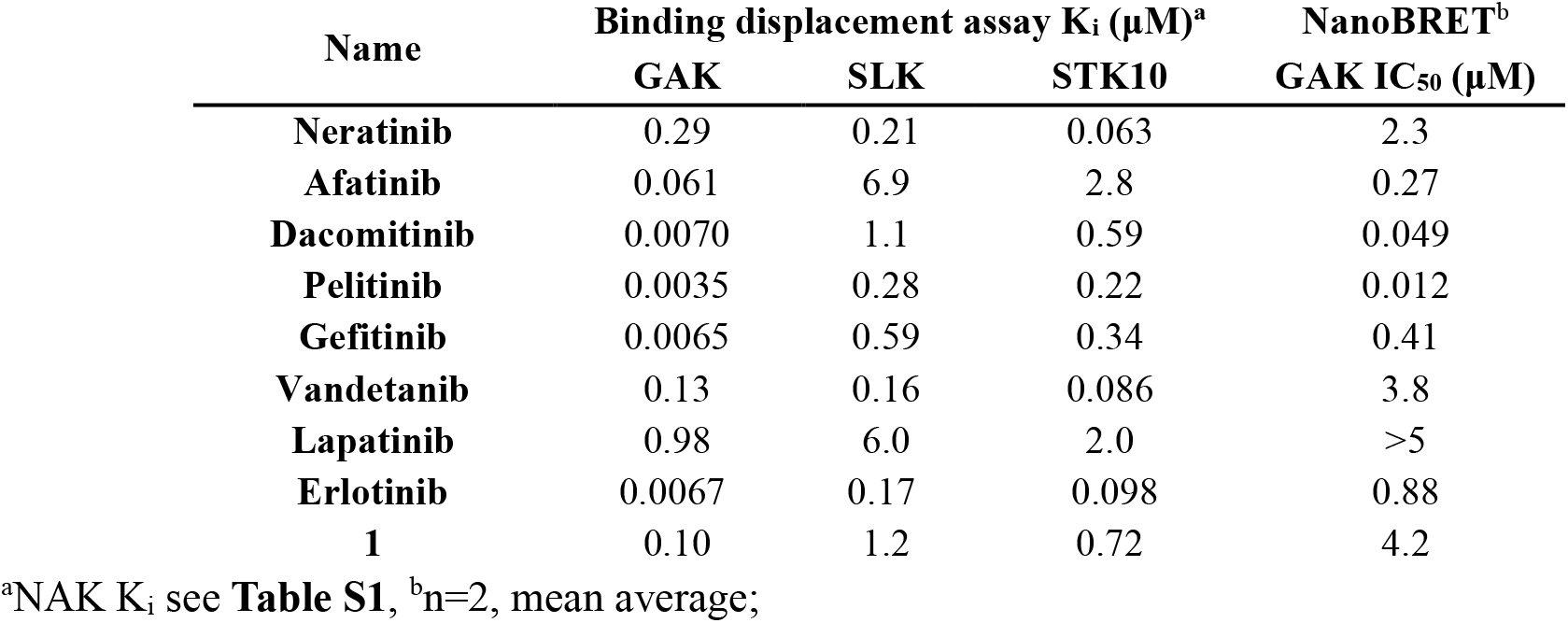
Screening results of literature inhibitors

### Screening of literature inhibitors to assess NAK and SLK/STK10 activity

We found neratinib to relatively potent on our main targets of interest GAK, SLK and STK10 with a 3-fold preference towards STK10. There was micromolar activity on the other NAK family members (**Table S1**), despite the low K_i_ (290 nM) on GAK the nanoBRET data demonstrated good potency at IC_50_ = 2.3 μM less than a 10-fold right shift. Afatinib, another covalent EGFR inhibitor was found to be a potent GAK inhibitor with a 3-fold increase and showed some selectivity over the other 5 kinases screened. Dacomitinib was found to be single digit nanomolar inhibitor of GAK with a potent GAK nanoBRET (IC_50_ = 49 nM). Pelitinib from the same class of covalent EGFR inhibitors with a 3-cyano quinoline hinge binder was the most potent GAK inhibitor screened from the literature compounds with an in-cell target engagement of IC_50_ = 12 nM. Pelitinib was also a good inhibitor of other NAK family members BMPK2 (K_i_ = 200 nM) and AKK1 (K_i_ = 570 nM) (**Table S1**) and SLK/STK10 K_i_ = 280 nM and 220 nM respectfully. Interestingly gefitinib was shown to have near equipotencie in GAK K_i_ compared with pelitinib, but a corresponding 34-fold drop off in the GAK nanoBRET compared to 3-fold with pelitinib. This highlights the important of combining both binding assay data and in-cell data to get a more complete compound profile. Gefitinib did however show the same off targets as with the covalent EGFR inhibitors, with GAK/SLK/STK10 all inhibited below one K_d_ = 100 nM. However, gefitinib showed only limited activity on AAK1/BMPK2 despite having the same substitution pattern on the aniline portion of the compound as pelitinib.

Vandetanib has a similar profile to neratinib, showing a preference for STK10 over GAK with potencies in the triple digit nanomolar range for SLK/STK10/GAK and limited activity on the other targets screened (**Table S1**). Lapatinib showed a drop off in activity on all 6 targets, this is likely due to the *para*-substituted benzyl on the aniline portion of the molecule accessing a deeper pocket within the ATP binding site of EGFR.^38^ Interestingly this is not the case with neratinib that has a similar benzyl substitution but with a 2-pyridyl instead of a *meta*-fluoro and a *meta*-chloro on the parent aniline ring system (Figure 1). Erlotinib was shown to be a potent inhibitor of nearly all 6 targets, only STK16 showing very limited binding (K_i_ = 42 μM). We observed potent inhibition of GAK at K_i_ = 6.7 nM with SLK/STK10/AAK1 and BMPK2 all below K_i_ = 400 nM. Erlotinib and gefitinib both showed a drop off in the GAK nanoBRET, highlighting the need for comprehensive data collection. Surprisingly the erlotinib quinoline (**1**) showed reduced GAK activity over erlotinib, highlighting the importance of the solvent exposed 6,7-substitution pattern within this scaffold.

### Synthesis of analogs to probe NAK and SLK/STK10

We synthesized the compounds **1**-**113** to probe the NAK/SLK/STK10 inhibitor landscape through nucleophilic aromatic displacement of 4-chloroquin(az)olines (**Scheme 1**).^13^ This enabled us to furnish the products in good-excellent overall yields (52-95 %). The powerful electron withdrawing trifluoromethyl group on the quinazoline (**30**, **35**, **61** & **66**) all had lower yields (21-23 %). The 6,7-difluoro (**57**) was also low yielding (21 %), with the 7-carbonitrile (**67**) (21 %) and 6-carbonitrile (**68**) (20 %) close behind. This was rounded off by two other lower yielding analogs **107** (45 %) and **63** (44 %) both having a sub-optimal yield likely due to the electronic of the ring systems.

**Scheme 1.**
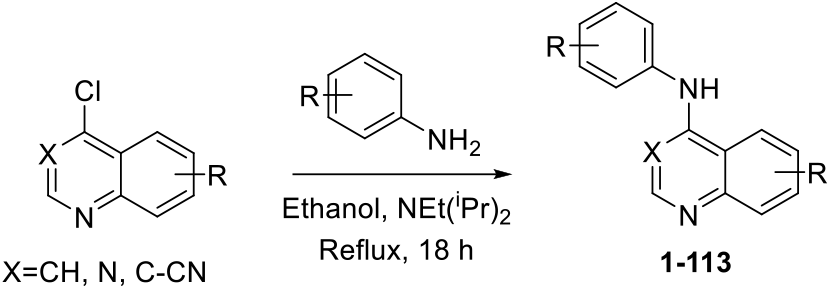
General synthetic procedure

We first screened 3 quinolines (**4**-**6**) with differing *meta*-substituted head groups and a fixed 6,7-dimethoxyquinoline core (Table 2, **Table S2**). We found the GAK data to be consistent with previous reports,^13^ but with the new cross-screening against SLK/STK10 we observed that kinase selectivity was able to be modulated towards SLK/STK10. The *meta*-alkyne (**4**) showed increased selectivity towards SLK/STK10 over the corresponding bromo (**5**) and methoxy (**6**) substitution.

**Table 2.** Initial screening results of *meta*-substituted head groups

| Name | R | Binding displacement assay K<sub>i</sub> (μM) |  |  | Nano-BRET<sup>a</sup> | Ratio<sup>b</sup> |
| --- | --- | --- | --- | --- | --- | --- |
|  |  | GAK | SLK | STK10 | GAK IC<sub>50</sub> (μM) |  |
| **4** | C≡C | 0.0035 | 0.38 | 0.18 | 0.042 | 51 |
| **5** | Br | 0.0025 | 5.8 | 2.3 | 0.51 | 920 |
| **6** | OMe | 0.0094 | 32 | 10 | 0.077 | 1100 |

<sup>a</sup>n=2, mean average; <sup>b</sup>NAK over STK10/SLK (See **Table S2**)

One of the major kinase targets of the 4-anilinoquin(az)oline scaffold is GAK. If we were going to get selectivity over GAK for SLK/STK10, first we the sought to investigate a known GAK inhibitor template. This enabled the development of an inhibition landscape and allowed for design deviations on the 4-anilinoquin(az)oline template (Table 3, **Table S3**). The *N*-(3,4,5-trimethoxyphenyl)quinolin-4-amine on a known potent GAK inhibitor chemotype.^11–13^

**Table 3.** Screening results of GAK inhibitor *N*-(3,4,5-trimethoxyphenyl)quinolin-4-amine template

| Name | R<sup>1</sup> | R<sup>2</sup> | Binding displacement assay K<sub>i</sub> (μM) |  |  | NanoBRET<sup>a,b</sup> | Ratio<sup>c</sup> |
| --- | --- | --- | --- | --- | --- | --- | --- |
|  |  |  | GAK<sup>a</sup> | SLK | STK10 | GAK IC<sub>50</sub> (μM) |  |
| **7** | H | H | 0.040 | >55 | 27.8 | 0.73 | 700 |
| **8** | F | H | 0.0057 | 5.1 | 45.9 | 0.27 | 900 |
| **9** | F | F | 0.0069 | 26 | 2.7 | 0.14 | 390 |
| **10** | Cl | H | 0.00068 | 11 | 1.0 | 0.21 | 1500 |
| **2** | Br | H | 0.0031 | 34 | 4.6 | 0.048 | 1500 |
| **11** | I | H | 0.0038 | 5.8 | 24.3 | 0.11 | 1500 |
| **12** | CF<sub>3</sub> | H | 0.0039 | 33 | 4.6 | 0.17 | 1200 |
| **13** | CN | H | 0.0015 | 19 | 1.7 | 0.36 | 1100 |
| **14** | OMe | H | 0.013 | >55 | 12 | 0.36 | 920 |
| **15** | SO<sub>2</sub>Me | H | 0.018 | 23 | 2.4 | 2.9 | 130 |
| **16** | H | F | 0.0024 | 17 | 1.6 | 0.54 | 670 |
| **17** | H | Cl | 0.0013 | 2.1 | 0.21 | 0.25 | 160 |
| **18** | H | Br | 0.0049 | 2.7 | 0.38 | 0.27 | 78 |
| **19** | H | I | 0.0044 | 1.7 | 0.21 | 0.074 | 48 |
| **20** | H | CF<sub>3</sub> | 0.0062 | 3.2 | 0.37 | 0.52 | 60 |
| **21** | H | CN | 0.0027 | 1.1 | 0.13 | 0.35 | 48 |
| **22** | H | OMe | 0.0026 | 13 | 1.3 | 0.44 | 500 |
| **3** | OMe | OMe | 0.00054 | 7.1 | 1.1 | 0.026 | 2000 |

<sup>a</sup>Consistent with previous report, <sup>b</sup>n=2, mean average; <sup>c</sup>NAK over STK10/SLK (See **Table S3**)

The simple unsubstituted quinoline (**7**) showed only limited STK10 binding (K_i_ = 28 μM) and no SLK binding. The introduction of a fluorine to the 6-position of the quinoline (**8**) provided a boost in binding to SLK, but also led to a corresponding increase in GAK binding so the ratio of (NAK)/(SLK/STK10) increased. The introduction of a second fluoro in the 7-position (**9**) switched the binding preference between SLK and STK10 but did lead to an almost 3-fold reduction in the ratio between NAK and SLK/STK10. However, the progression of increasingly large mono-substituted 6-position halogens: chloro (**10**), bromo (**2**), iodo (**11**) and trifluoromethyl (**12**) yielded a static ratio at around 1500-fold selective towards the NAK family. The switch to the 6-position cyano (**13**) and methoxy (**14**) showed minimal improvements to the selectivity ratio. The larger methylsulfone (**15**) was significantly weaker on GAK compared to the rest of the series, but still maintained a similar SLK/STK10 binding ratio.

The 7-position halogens (**16**-**20**) demonstrated an improved ratio over the 6-position with the halogens larger than the chlorine, showing consistent double-digit ratios. This was also the case with the 7-position cyano (**21**). The 7-position methoxy (**22**) showed a 10-fold drop off for STK10 compared with **21** which pushed the ratio of NAK vs SLK/STK10 above 500. The 6,7-dimethoxy quinoline (**3**), showed the worst ratio in the series (>2000), due in part to the picomolar potency on GAK (K_i_ = 540 pM).

The previous NAK family data on the 4-anilinoquinoline and 4-anilinoquinazoline cores suggested that the quinazoline was a weaker binder of GAK with certain anilines.^13^ We observed that this was not necessarily the case with the literature compounds (Table 1, **Table S1**). We decided to explore this as an option to maintain SLK/STK10 potency while dialing back the GAK binding (Table 4, **Table S4**).

**Table 4.** Screening results of trimethoxyphenylquinazoline template

| Name | R<sup>1</sup> | R<sup>2</sup> | Binding displacement assay K<sub>i</sub> (μM) |  |  | NanoBRET<sup>a</sup> | Ratio<sup>b</sup> |
| --- | --- | --- | --- | --- | --- | --- | --- |
|  |  |  | GAK | SLK | STK10 | GAK IC<sub>50</sub> (μM) |  |
| **23** | H | H | 0.034 | 50 | 6.0 | 4.6 | 180 |
| **24** | Me | H | 0.022 | 25 | 5.1 | 2.1 | 230 |
| **25** | F | H | 0.033 | 9.4 | 11 | 3.6 | 280 |
| **26** | F | F | 0.093 | >55 | 7.1 | >5 | 76 |
| **27** | Cl | H | 0.017 | 47 | 7.0 | 2.4 | 410 |
| **28** | Br | H | 0.010 | 21 | 3.8 | 1.8 | 380 |
| **29** | I | H | 0.011 | 18 | 4.1 | 0.89 | 370 |
| **30** | CF<sub>3</sub> | H | 0.037 | 25 | 4.0 | 4.5 | 110 |
| **31** | H | F | 0.055 | 44 | 6.7 | >5 | 120 |
| **32** | H | Cl | 0.030 | 3.8 | 0.81 | 3.2 | 27 |
| **33** | H | Br | 0.025 | 2.5 | 0.43 | 4.2 | 17 |
| **34** | H | I | 0.021 | 1.8 | 0.21 | 2.4 | 10 |
| **35** | H | CF<sub>3</sub> | 0.079 | 8.2 | 1.2 | >5 | 15 |
| **36** | H | CN | 0.21 | 10 | 1.6 | >5 | 7.6 |
| **37** | CN | H | 0.099 | 44 | 7.2 | >5 | 73 |
| **38** | SO<sub>2</sub>Me | H | 0.60 | 44 | 7.4 | >5 | 12 |
| **39** | OMe | H | 0.022 | 18 | 4.3 | 3.4 | 200 |
| **40** | OMe | OMe | 0.0014 | 2.1 | 0.2 | 0.35 | 140 |
| **41** | H | OMe | 0.012 | 7.6 | 1.5 | 1.6 | 130 |
| **42** | 6,7-(OCH<sub>2</sub>CH<sub>2</sub>OMe)<sub>2</sub> |  | 0.026 | 6.8 | 0.58 | 3 | 22 |

<sup>a</sup>n=2, mean average; <sup>b</sup>NAK over STK10/SLK (See **Table S4**)

While the unsubstituted *N*-(3,4,5-trimethoxyphenyl)quinazolin-4-amine (**23**) and 6-position methyl (**24**) demonstrated almost equipotence GAK binding, but the ratio between NAK vs SLK/STK10 did show some widening. The introduction of a 6-position fluoro (**25**) had only a limited effect on the activity profile, but the introduction of a second fluorine at the 7-position (**26**) did reduce the GAK activity by 3-fold and led to an almost 4-fold increase in the selectivity ratio; and interestingly removed all SLK affinity. The introduction of larger halogens in the 6-position (**27-30**) show potent GAK profiles, with only the trifluoromethyl (**30**) showing a similar profile to 6-fluoro (**25**). In addition, **30**-**32** showed increased binding to STK16 (K_i_ = 2-3 μM) (**Table S4**).

The switch to the 7-position halogen substitution (**31**-**35**) yielded a slightly weaker GAK profile, consistent with the quinolines 7-position halogens (**16**-**20**). The 7-fluoro (**31**) appeared too small to impact the binding, but the 7-chloro (**32**) to 7-iodo (**34**) showed progressively better NAK vs SLK/STK10 ratios down to 10. The 7-cyano substitution (**36**) proved to be a further improvement due in part to the significant reduction in GAK binding to K_i_ = 210 nM and complete obliteration of other NAK family members (**Table S4**). However, **36** did not had the same potency against STK10 observed with the 7-position halogens (**32**-**34**).

The switch to the 6-cyano (**37**) yielded an increase in GAK binding by 2-fold, with the 6-methylsulfone (**38**) showing a similar overall 6 kinase profile but with a 6-fold drop compared with **37**. While the 6-methoxy (**39**) showed a jump of 5-fold GAK binding compared with **37**, the 6,7-dimethoxy (**40**) showed a 15-fold increase in GAK binding and a jump in GAK target engagement. However, with a K_i_ = 200 nM on STK10 the overall ratio showed a small improvement, significantly better than the quinoline (**3**). The 7-methoxy (**41**) reversed the trend observed with the 7-position halogens (**34**-**37**) with an improved GAK binding profile. The trimethoxyaniline erlotinib quinazoline (**42**) showed a 10-fold drop compared with **40** and a 4-fold drop compared with erlotinib itself. However, the ratio between NAK and SLK/STK10 was an almost 100-fold improvement towards STK10/SLK compared with **3**.

The cone angle of the quinoline/aniline amine bond is around 60 degrees while the quinazoline is almost flat.^13^ The 3-cyanoquinoline with the projected cyano in the arched region of the molecule should push the aniline into an almost perpendicular position with respect to the quinoline. This combined with the new electronic conformation provides for a new chemical space to seek the separation of activities between the three main kinases (GAK, SLK & and STK10) of interest.

The unsubstituted trimethoxyphenylquinoline-3-carbonitrile (**43**) showed a halving in activity on GAK compared with quinazoline (**23**) and quinoline (**7**) (Table 5). There was no improvement on SLK/STK binding meaning the ratio between GAK and SLK/STK10 was static. The introduction of a larger halogen in the 6-position (**44**-**46**) boosted GAK binding and GAK in-cell target engagement, SLK/STK10 binding was negligible. The 6-methoxy (**47**) showed a 3-fold boost in the GAK nanoBRET compared to **43**, but no ratio improvement.

**Table 5.** Screening results of trimethoxyphenylquinoline-3-carbonitrile template

| Name | R<sup>1</sup> | R<sup>2</sup> | Binding displacement assay K<sub>i</sub> (μM) |  |  | NanoBRET<sup>a</sup><br>GAK IC<sub>50</sub> (μM) | Ratio<sup>b</sup> |
| --- | --- | --- | --- | --- | --- | --- | --- |
|  |  |  | GAK | SLK | STK10 |  |  |
| **43** | H | H | 0.0088 | 7.7 | 4.7 | 0.33 | 530 |
| **44** | Cl | H | 0.0038 | 53 | 26 | 0.24 | 6700 |
| **45** | Br | H | 0.0023 | 9.4 | 11 | 0.20 | 4100 |
| **46** | I | H | 0.0048 | 9.7 | 4.2 | 0.26 | 880 |
| **47** | OMe | H | 0.0024 | 3.2 | 1.9 | 0.092 | 790 |
| **48** | OMe | OMe | 0.00085 | 0.57 | 0.42 | 0.006 | 140 |
| **49** | SO<sub>2</sub>Me | H | 0.21 | 1.2 | 1.5 | >5 | 6 |
| **50** | H | Cl | 0.0066 | 1.4 | 0.79 | 0.47 | 120 |
| **51** | H | Br | 0.0029 | 0.71 | 0.44 | 0.14 | 150 |
| **52** | H | I | 0.0052 | 0.49 | 0.20 | 0.20 | 38 |
| **53** | H | OMe | 0.0020 | 2.1 | 1.2 | 0.078 | 580 |

<sup>a</sup>n=2, mean average; <sup>b</sup>NAK over STK10/SLK (See **Table S5**)

The 6,7-dimethoxy (**48**) is the most potent GAK nanoBRET result ever observed. Improvement was observed across all kinases screened, with SLK/STK10 and AAK1 all showing binding below K_i_ = 600 nM. The 6-methylsulfone (**49**) showed a drop on GAK binding, but also a reducing of binding across all 6 kinases. **49** was still the lowest ratio observed between NAK vs SLK/STK10. There was no difference when switching to the 7-position halogens (**50**-**52**) on the GAK binding observed. However other NAK family members (AAK1, BMPK2, STK16) were in-active, while STK10/SLK showed activity mostly below 1 μM. The progression of increasing size from fluorine to iodine did however demonstrate an increase in selectivtity towards SLK/STK10 over the NAK family. This highlighted that selectivity maybe achievable on this template. The fragility of this selectivity was demonstrated by the switch between 7-iodo (**52**) to 7-methoxy (**53**) with a reversal of selectivity towards GAK and a reduction in selectivity ratio to above 500.

### Development of *meta*-alkyne substituted aniline analogs

In order to find more SLK/STK10 selective compounds we switch back to the original *meta*-alkyne substituted aniline which demonstrated the original selectivity preference towards SLK/STK10 in **4**. Combining this with the screening knowledge gained with **4**-**53**, we decided to pursue the quinazoline core. This is due to the consistent 5-10-fold drop off in GAK binding/activity observed compared with the quinoline core.

The unsubstituted analog (**54**) showed a reduction in GAK binding that was 3-5-fold greater that the trimethoxy counterparts (**7**, **23**, **43**), the limited activity on SLK/STK10 (Table 6). The addition of a methyl in the 6-position (**55**) provided a 4-fold improvement in GAK binding with a limited improvement in SLK/STK10 binding. A single fluorine in the 6-position (**56**) showed no significant change in profile across the kinases screened, however addition of a second fluorine in the 7-position (**57**) did boost the SLK/STK10 while maintain GAK binding bringing the ratio of GAK vs SLK/STK10 down to 34. Increasing the size of the 6-position halogen (**58**-**60**) above fluorine increased the GAK binding while maintaining the SLK/STK10 in place, almost tripling the ratio of 57. Increasing to the trifluoromethyl (**61**) reduced the GAK binding by more than 5-fold, maintaining the SLK/STK10 activity and pushing the ratio down to just above 20.

**Table 6.** Screening results of *meta*-acetylene quinazoline template

| Name | R<sup>1</sup> | R<sup>2</sup> | Binding displacement assay K<sub>i</sub> (μM) |  |  | NanoBRET<sup>a</sup><br>GAK IC<sub>50</sub> (μM) | Ratio<sup>b</sup> |
| --- | --- | --- | --- | --- | --- | --- | --- |
|  |  |  | GAK | SLK | STK10 |  |  |
| **54** | H | H | 0.17 | 23 | 10 | 4.5 | 61 |
| **55** | Me | H | 0.043 | 4.3 | 2.6 | 3.6 | 60 |
| **56** | F | H | 0.070 | 8.5 | 5.5 | 2.0 | 79 |
| **57** | F | F | 0.085 | 3.8 | 2.9 | >5 | 34 |
| **58** | Cl | H | 0.030 | 3.9 | 2.8 | >5 | 93 |
| **59** | Br | H | 0.028 | 3.4 | 2.5 | >5 | 89 |
| **60** | I | H | 0.044 | 4.3 | 3.7 | >5 | 84 |
| **61** | CF<sub>3</sub> | H | 0.150 | 3.2 | 3.4 | >5 | 21 |
| **62** | H | F | 0.022 | 1.2 | 1.2 | 4.1 | 55 |
| **63** | H | Cl | 0.035 | 0.36 | 0.44 | >5 | 10 |
| **64** | H | Br | 0.031 | 0.26 | 0.29 | >5 | 8.4 |
| **65** | H | I | 0.020 | 0.18 | 0.15 | >5 | 9 |
| **66** | H | CF<sub>3</sub> | 0.088 | 1.2 | 1.0 | >5 | 11 |
| **67** | H | CN | 0.10 | 0.26 | 0.31 | >5 | 2.6 |
| **68** | CN | H | 0.17 | 7.9 | 4.7 | >5 | 28 |
| **69** | SO<sub>2</sub>Me | H | 0.42 | 1.3 | 1.3 | >5 | 3.1 |
| **70** | OMe | H | 0.024 | 2.8 | 2.2 | 2.9 | 92 |
| **71** | OMe | OMe | 0.0010 | 0.14 | 0.12 | 0.079 | 120 |
| **72** | H | OMe | 0.0088 | 0.67 | 0.46 | 1.2 | 52 |
| **73** | OCH<sub>2</sub>O |  | 0.019 | 2.7 | 1.5 | 1.7 | 79 |
| **74** | CH<sub>2</sub>OCH<sub>2</sub> |  | 0.035 | 3.6 | 1.7 | 3.2 | 49 |
| **Erlotinib** | 6,7-(OCH<sub>2</sub>CH<sub>2</sub>OMe)<sub>2</sub> |  | 0.0067 | 0.17 | 0.10 | 0.88 | 15 |

<sup>a</sup>n=2, mean average; <sup>b</sup>NAK over STK10/SLK (See **Table S6**)

Switching to the 7-position halogens (**62**-**66**) yielded an improvement in the ratio of the NAK vs SLK/STK10. The 7-fluoro (**62**) was too small to significantly impact the profile, however increasing the size improved the GAK vs SLK/STK10 profile to single digit ratios. The 7-iodo (**65**) provided the best STK10 binding with a K_i_ = 150 nM with a ratio to GAK to 9. The 6-cyano (**67**) provided potential route to a selective SLK/STK10 inhibitor with a ratio of just 2.6 and GAK binding of K_i_ = 100 nM. Switching to the 7-position cyano (**68**) proved to be good to reducing GAK binding but also knocked SLK/STK10 back to the low micromolar range.

The 6-methylsulfone (**69**) reduced the GAK binding to K_i_ = 420 nM, while simultaneously increasing the affinity to just above 1 μM for both SLK/STK10. This brought the ratio down to the same level as **67** while reducing the GAK binding selectively. The switch to the 6-methoxy (**70**) increased GAK binding by 18-fold while maintaining SLK/STK10 binding at similar levels to **69**; **70** also showed increased AAK1 binding with a K_i_ = 930 nM (**Table S6**). The introduction of a second methoxy in the 7-position (**71**) provided a strong template for GAK consistent with **3**, **40** and **48** with binding at K_i_ = 1 nM. However, the ratio was not as high as previous 6,7-dimethoxy examples and potency of SLK and STK10 was K_i_ = 140 nM and 120 nM respectively with a corresponding increase in AAK1 and BMP2K at K_i_ = 200 nM and 600 nM respectively. The removal of the 6-position methoxy from **71** highlights that the methoxy groups are synergetic and not additive as the 7-methoxy analog (**72**) was 2.5-fold more potent on GAK than **70**, with a similar drop on SLK/STK10 and AAK1 binding. The closing of the 6,7-dimethoxy to form a 5-membered ring (**73**) or a 6-membered ring (**74**) reduced the GAK binding but also reduced binding to other the other kinases screened. Erlotinib was not equi-potent to the 6,7-dimethoxy with an increase in selectively towards SLK/STK10 but still more potent on GAK. The in-cell target engagement on GAK is surprisingly low considering the potent GAK K_i_.

To increase selectivity towards SLK/STK we explored the 3-cyanoquinoline hinge binder in order to manipulate the electronics and sterics of the template (Table 7). The unsubstituted analog (**75**) showed good GAK binding with relatively potent SLK/STK10 potency. The introduction of progressively larger 6-position halogens (**76**-**79**) eroded the SLK/STK10 potency, while GAK binding was maintained. The increase in halogen size also appeared to correlate with a decrease in-cell target engagement despite only a limited change in binding. The introduction of a 6-methylsulfone (**80**) lead to an overall decrease in binding with a drop of 10-fold in GAK binding compared to **75**.

**Table 7.** Screening results of meta-acetylene quinazoline template

| Name | R<sup>1</sup> | R<sup>2</sup> | Binding displacement assay K<sub>i</sub> (μM) |  |  | NanoBRET<sup>a</sup> | Ratio<sup>b</sup> |
| --- | --- | --- | --- | --- | --- | --- | --- |
|  |  |  | GAK | SLK | STK10 | GAK IC<sub>50</sub> (μM) |  |
| 75 | H | H | 0.018 | 0.35 | 0.34 | 1.5 | 19 |
| 76 | F | H | 0.026 | 0.54 | 0.58 | 3.8 | 21 |
| 77 | Cl | H | 0.016 | 0.71 | 0.72 | 3.2 | 44 |
| 78 | Br | H | 0.017 | 1.2 | 1.1 | 4.4 | 65 |
| 79 | I | H | 0.020 | 1.1 | 1.0 | >5 | 50 |
| 80 | SO<sub>2</sub>Me | H | 0.18 | >55 | 42 | >5 | 230 |
| 81 | OMe | H | 0.0063 | 0.27 | 0.31 | 0.59 | 43 |
| 82 | OMe | OMe | 0.00032 | 0.018 | 0.027 | 0.040 | 56 |
| 83 | H | OMe | 0.0035 | 0.062 | 0.073 | 0.29 | 18 |
| 84 | H | Cl | 0.028 | 0.18 | 0.21 | >5 | 6.4 |
| 85 | H | Br | 0.027 | 0.16 | 0.13 | >5 | 4.9 |
| 86 | H | I | 0.019 | 0.075 | 0.080 | >5 | 3.9 |

<sup>a</sup>n=2, mean average; <sup>b</sup>NAK over STK10/SLK (See **Table S7**)

An affinity well was observed around the methoxy substitutions (**81**-**83**). These analogs showed a high potency across most of the kinases except STK16 (**Table S7**). The 6,7-dimethoxy (**82**) was the most potent GAK binder observed to date with picomolar affinity (K_i_ = 320 pM), this did not translate to an equivalent high GAK in-cell target engagement value, despite **82** still being highly active (IC_50_ = 40 nM). The introduction of 7-positionhalogens (**84**-**86**) boosted the SLK/STK activity while keeping the GAK binding static. This culminated in the 7-iodo compound (**86**) having a potency on SLK/STK10 of K_i_ = 75 nM and 80 nM respectively and a NAK vs SLK/STK10 ratio of just under 4.

### Optimization of the aniline head group

Four compounds (**60**, **65**, **67**, **71**) with the some of the best potency and selectivity towards SLK/STK10 we chosen for further optimization within the *meta*-acetylene aniline head group. GAK is known to be sensitive to aniline modifications particularly in the *ortho-* and *para-*position.^38^ We first selected **67**, the 7-cyano quinazoline and installed several small halogens (**87**-**90**) on the aniline portion of the molecule. The three fluorine derivatives (**87**-**89**) showed a selectivity ratio improvement whereas the 4-chloro analog (**91**) did not. The addition of a fluoro in the 6-position of the aniline ring (**87**) lead to a 120-fold reduction in GAK binding with maintenance of activity of both SLK and STK10. Compound **87** had the highest selectivity ratio of any compound synthesized with a ratio below 1, where GAK binding was K_i_ = 1.2 μM and SLK/STK10 binding was K_i_ = 560 and 670 nM respectively. The analogs with the fluoro in the 2-position (**88**) and the 4-position (**89**) showed slightly reduced overall binding to the kinases screened. There was very limited activity in the wider NAK family for any of these analogs (**67**, **87**-**90**) (Table S8)

**Table 8.** Screening results of optimized *meta*-acetylene quinazolines (**60**, **65**, **67**, **71**)

| Name | R<sup>1</sup> | R<sup>2</sup> | R<sup>3</sup> | R<sup>4</sup> | R<sup>5</sup> | Binding displacement assay K<sub>i</sub> (μM) |  |  | NanoBRET<sup>a</sup> | Ratio<sup>b</sup> |
| --- | --- | --- | --- | --- | --- | --- | --- | --- | --- | --- |
|  |  |  |  |  |  | GAK | SLK | STK10 | GAK IC<sub>50</sub> (μM) |  |
| **67** | H | CN | H | H | H | 0.010 | 0.26 | 0.31 | >5 | 2.6 |
| **87** | H | CN | F | H | H | 1.2 | 0.56 | 0.67 | >5 | 0.47 |
| **88** | H | CN | H | F | H | 0.33 | 1.1 | 0.50 | >5 | 1.5 |
| **89** | H | CN | H | H | F | 0.55 | 0.69 | 0.87 | >5 | 1.2 |
| **90** | H | CN | H | H | Cl | 0.32 | 6.3 | 1.6 | >5 | 5 |
| **65** | H | I | H | H | H | 0.020 | 0.18 | 0.15 | >5 | 9 |
| **91** | H | I | F | H | H | 0.11 | 0.39 | 0.25 | >5 | 2.3 |
| **92** | H | I | H | F | H | 0.093 | 0.86 | 0.27 | >5 | 2.9 |
| **93** | H | I | H | H | F | 0.035 | 0.11 | 0.11 | >5 | 3.1 |
| **94** | H | I | H | H | Cl | 0.092 | 0.93 | 0.24 | >5 | 2.6 |
| **60** | I | H | H | H | H | 0.044 | 4.3 | 3.7 | >5 | 84 |
| **95** | I | H | F | H | H | 0.19 | 4.5 | 4.9 | >5 | 24 |
| **96** | I | H | H | F | H | 0.29 | 18 | 6.7 | >5 | 23 |
| **97** | I | H | H | H | F | 0.068 | 1.6 | 0.87 | >5 | 13 |
| **98** | I | H | H | H | Cl | 0.14 | 7.1 | 1.0 | >5 | 7.1 |
| **71** | OMe | OMe | H | H | H | 0.0010 | 0.14 | 0.12 | 0.079 | 120 |
| **99** | OMe | OMe | F | H | H | 0.0058 | 0.28 | 0.22 | 0.39 | 38 |
| **100** | OMe | OMe | H | F | H | 0.0063 | 0.67 | 0.25 | 0.40 | 40 |
| **101** | OMe | OMe | H | H | F | 0.0024 | 0.072 | 0.047 | 0.14 | 20 |
| **102** | OMe | OMe | H | H | Cl | 0.0034 | 0.33 | 0.075 | 0.39 | 22 |

<sup>a</sup>n=2, mean average; <sup>b</sup>NAK over STK10/SLK (See **Table S8**)

The 7-iodo quinazoline (**65**) showed some potency towards SLK/STK10 with a promising selectivity ratio. The small halogen modifications (**91**-**94**) yielded compounds with improved selectivity ratios primarily modulated through the GAK binding affinity. Interestingly the 4-chloro (**94**) was still relatively active against GAK unlike with **90**. There was no improvement in SLK/STK10 potency and no activity in the wider NAK family (**Table S8**). The corresponding 6-iodo derivative (**60**) had a larger selectivity deficit towards SLK/STK10, however this was still able to be reduced by the introduction of halogens into the aniline portion of the molecule (**95**-**98**). The 4-fluoro (**97**) was able to drive down affinity for STK10 to K_i_ = 870 nM with a ratio to NAK of 13. The 4-chloro (**98**) was able to improve that ratio with a slight drop in potency to 1 μM.

The 6,7-dimethoxyquinazoline (**71**) is one of the better ATP competitive kinase inhibitor substitution patterns on this template. Despite the potent GAK profile and triple digit NAK vs SLK/STK10 ratio the halogen substitutions (**99**-**102**) proved to be successful in driving down the selectivity and potency on SLK/STK10. The ratios were improved by increasing potency on SLK/STK10 with the 4-fluoro (**101**) being the most potent binder on SLK/STK10 with a K_i_ = 72 nM and 47 nM respectively with a ratio of 20.

### Halogenated aniline analogs

Finally, to map the chemical space and explore the halogen substitutions further we synthesized a series of 6-bromo quinoline and quinazolines derivatives with differing halogenated substitution patterns (**103**-**113**). While binding was maintained almost consistently on GAK, affinity towards SLK/STK10 was limited (Table 9). The bulkier substitutions, 3,4,5-trifluoro (**103**), 3,4-difluoro (**104**) and 3,5-difluoro (**105**) all showed between K_i_ = 10-25 nM binding to GAK but no activity on the other 5 kinases (**Table S9**). Movement of the second fluorine around the ring (**106**-**107**) maintaining one fluorine in the 6-position meant a small increase in binding across the NAK-family but limited improvement for SLK/STK10. The mono-substituted fluoro compounds (**108**-**111**) had surprisingly similar properties, but the 4-fluoro quinazoline did show some reduction in GAK binding and an increase in AAK1 binding to K_i_= 490 nM (**Table S9**). Switching to a bromo in the 3-position (**112**) did not immediately improve the NAK vs SLK/STK10 selectivity ratio but it did remove all activity across the wider NAK family. A switch to the quinazoline (**113**) increased binding to SLK/STK10 brought the ratio down to double digits. However, the increases in affinity across the NAK family made this ratio not a true reflection of overall selectivity improvement in this case.

**Table 9.** Screening results of halogenated quinolines and quinazolines

| Name | R<sup>1</sup> | R<sup>2</sup> | R<sup>3</sup> | R<sup>4</sup> | X | Binding displacement assay K<sub>i</sub> (μM) |  |  | NanoBRET<br>GAK IC<sub>50</sub> (nM) | Ratio<sup>a</sup> |
| --- | --- | --- | --- | --- | --- | --- | --- | --- | --- | --- |
|  |  |  |  |  |  | GAK | SLK | STK10 |  |  |
| 103 | H | F | F | F | C-H | 0.023 | >55 | >55 | 4.4 | >2400 |
| 104 | H | H | F | F | C-H | 0.011 | >55 | >55 | >5 | >5000 |
| 105 | H | F | H | F | C-H | 0.012 | >55 | >55 | >5 | >4600 |
| 106 | F | H | F | H | C-H | 0.035 | 38 | 11 | 2.7 | 310 |
| 107 | F | H | H | F | C-H | 0.015 | >55 | >55 | 0.87 | 3700 |
| 108 | H | H | F | H | C-H | 0.015 | 36 | 21 | 0.94 | 1400 |
| 109 | H | H | F | H | N | 0.12 | >55 | >55 | >5 | 460 |
| 110 | H | F | H | H | C-H | 0.019 | >55 | >55 | 1.9 | 2900 |
| 111 | F | H | H | H | C-H | 0.049 | >55 | 30 | 2.6 | 610 |
| 112 | H | Br | H | H | C-H | 0.072 | >55 | >55 | >5 | >760 |
| 113 | H | Br | H | H | N | 0.084 | 9.4 | 6.0 | >5 | 71 |

<sup>a</sup>n=2, mean average; <sup>b</sup>NAK over STK10/SLK (See Table S9)

### GAK K_i_ vs GAK nanoBRET

We measured GAK affinity of the 4-anilinoquin(az)oline analogues with a wide range of nanoBRET values and affinity over several orders of magnitude in the TR-FRET binding assay. The two binding assays were highly correlated, as shown by a plot of IC_50_ (nM) versus log K_i_ (nM) which has R^2^=0.73 (Figure 2). The cut off for effective GAK target engagement in the nanoBRET (IC_50_ >5 μM) is roughly GAK K_i_ = 20 nM or below.

**Figure 2.**
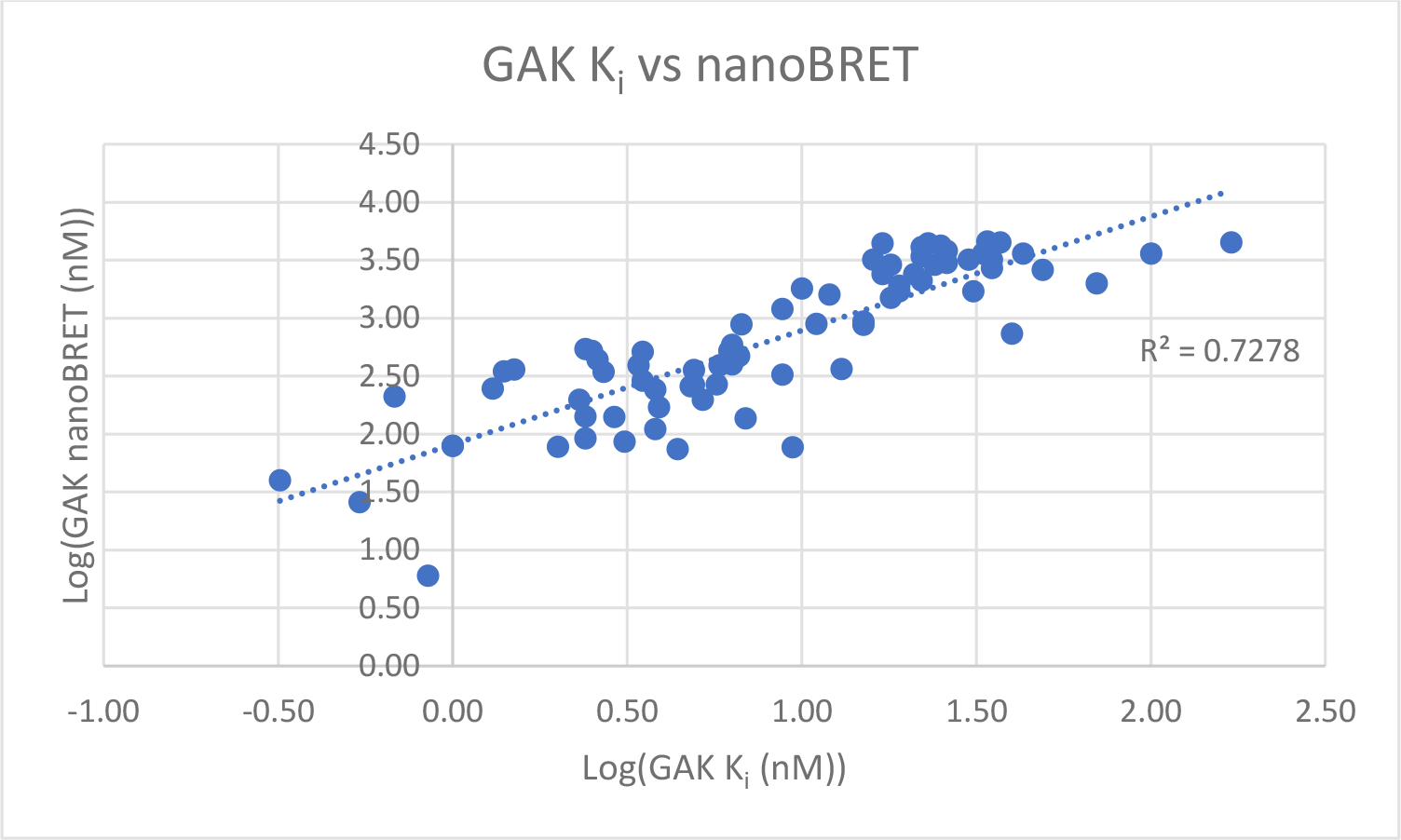
Comparison of GAK K_i_ vs GAK nanoBRET

### Small Molecule Crystal Structures

In order to better understand how the structural characteristics of the compounds might influence the kinase activity profile we solved a series of small molecule crystal structures (**Table S10.** Figure 4). All the compounds apart from **44**, **50**, **53** and **67** crystallized as chloride salts. Each of the compounds have two strong hydrogen bond donors so in the solid-state typically form hydrogen bonded tapes with the chloride ion as the acceptor. The structures which include solvent water (**7**, **11**, **15-17**, **20**, **24**, **27**, **34**, **36**, **39-40**, **43**, **45-46**, **53**) display more complex hydrogen bond networks. We observed that the quinolines (**7**, **11**, **15**-**20**, **22**) had a narrow range of C-N-C-C torsion angles of 46°-71°. This resulted in two narrow distributions of plane angles between 48°-74° and 115°-131° arising from the restricted rotation of the 4-anilinoquinoline which is generally out of plane due to the steric interaction between the C-3 C-H and the aniline ring. The quinolines (**7**, **15**-**17**, and **22**) generally packed in the triclinic P-1 space group, although **11**, **18** and **20** were monoclinic.

**Figure 4.**
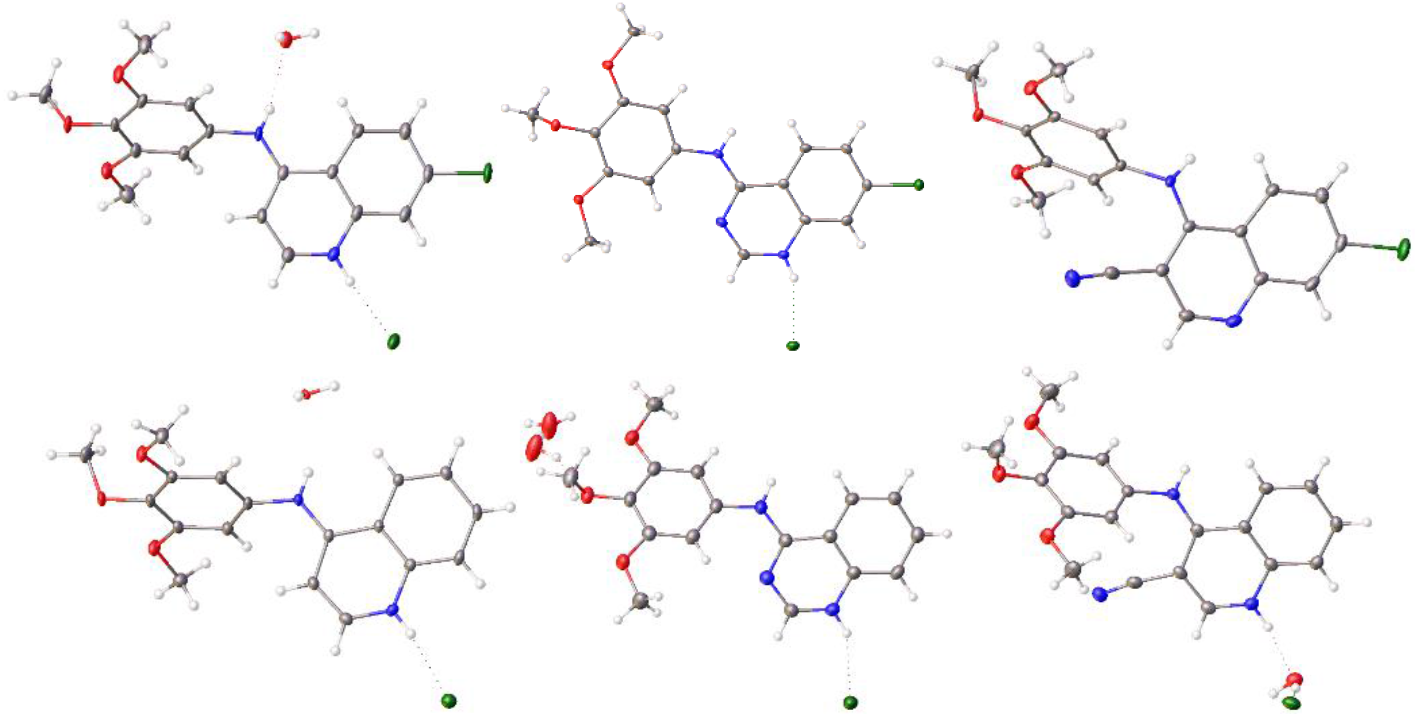
x-ray crystal structures of a set of quin(az)olines (**7**, **17**, **23**, **32**, **43**, **50**) exemplifying the plane angles observed

The quinazolines (**24**, **27**-**29**, **31**-**34**, **37**, **67**) packed in the triclinic crystal system, however **23**, **25**, **36**, **40** and **68** were monoclinic and **39** was orthorhombic. The flat planarity allows for a wider variance in crystal packing. Interestingly **23** had a bigger twist at 38°, but this only resulted in a plane angle of 40°. However, **39** demonstrated a disruption that pushed the aniline ring system out of plane like the quinoline structures (**7**, **11**, **15**-**20**, **22**) with a C-N-C-C torsion angle of 50° degrees and a plane angle of 132°. Additionally the *meta*-acetyl quinazoline, **67**, crystallizes with a C-N-C-C torsion angle of 29° and a plane angle of 29°. The rest of the quinazolines **23**-**25**, **27**-**29**, **31**-**37**, **39**-**40** were near planar (1.4°-7.9°). The 3-cyanoquinoline derivatives **43**, **45**, **46**, **50**, **53** were monoclinic and **44** orthorhombic, with no triclinic packing examples observed. The C-N-C-C torsion angle range between 44°-51° with plane angles ranging between 60°-122°. The projected C-3 cyano group projects out pushing the aniline into an orthogonal orientation.

### GAK QSAR Model

We constructed an explanatory field-based 3D QSAR model, which would agree with SAR of reported structures, as well as with recently released x-ray structures and molecular docking simulations.^39–40^ The 3D-QSAR model was constructed in a way that main fields could be overlaid with co-crystallized ligands and favorable docking poses. WaterMap simulations were computed for active site of GAK (PDB:5Y80) as solvated and with presence of selected active ligands (co-crystal and **82**).^40^ Docking suggests that the overall shape of the current ligand series aligns well with kinase back pocket, and also fully fills typical hydrogen bond interaction with hinge residues (Figure 5).

**Figure 5.**
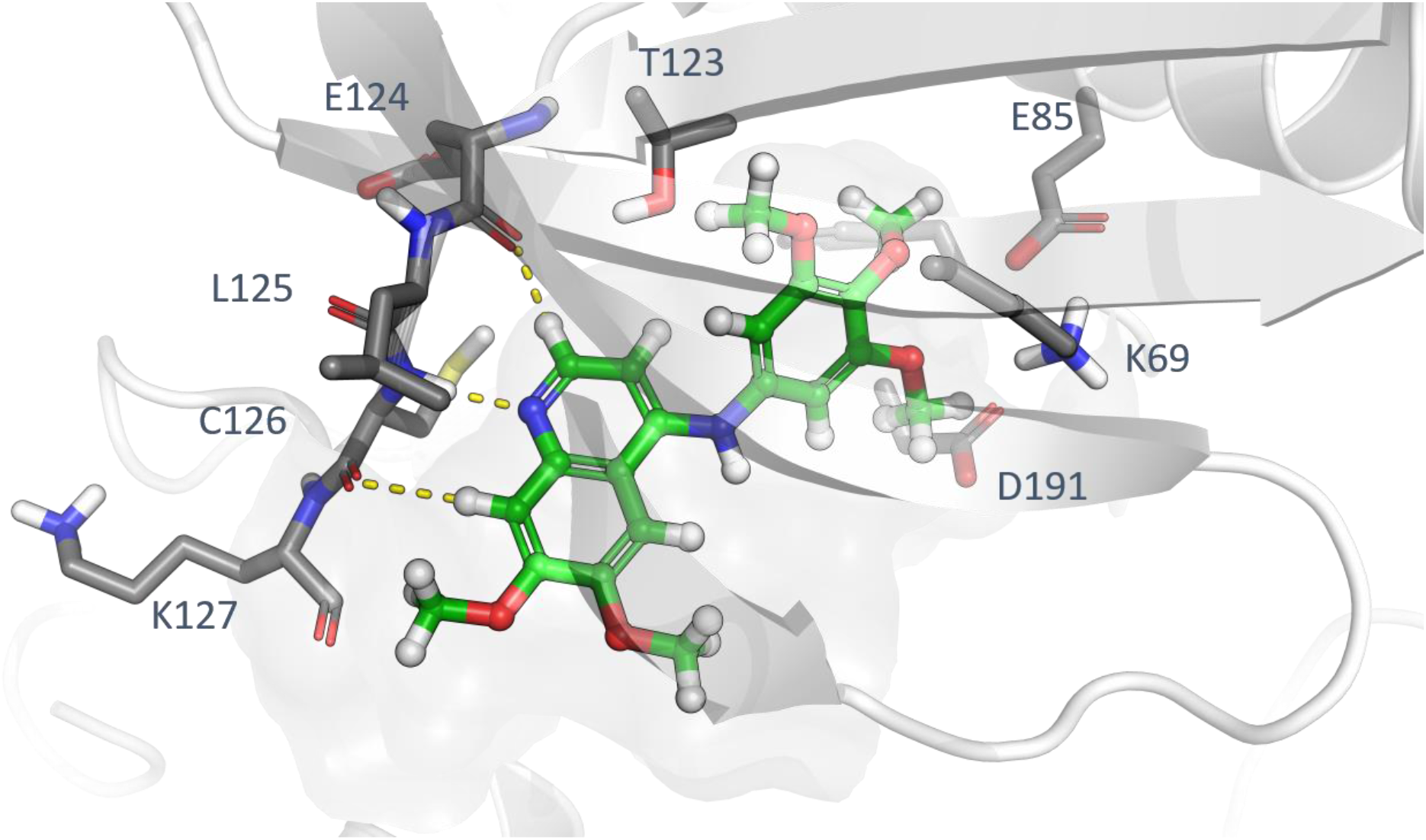
Docking of quinoline scaffold to GAK active site (PDB:5Y80, compound **3**).

The quinoline scaffold is able to form a hydrogen bond interaction to main chain amide of C126 and two additional aromatic hydrogen bond contacts to main chain carbonyl of C126 and L125. The co-crystalline structures of the GAK ATP binding site,^40^ show an additional lipophilic space towards the beta sheet, next to the mainchain of L68, K69 and T123 the where small substituents like methoxy or ethyne are able to orient. This pocket also contains a high energy water that appears to be displaced with certain orientations and substituent patterns.^13, 41^

We constructed 3D-QSAR models for GAK, SLK and STK10 kinases using the field based QSAR functionality of Schrödinger Maestro 2018-4 (Table 10). Flexible ligand alignments were performed using a Bemis-Mucko method,^42^ enabling recognition of the largest common scaffolds using a favorable docking pose of structurally representative derivative as alignment template. Torsion angles of larger substituents, such as methoxy groups, were manually adjusted to fit template structures when needed. The models were constructed by defining 80 % of compounds randomly as training sets and rest of the compounds were considered as test set. Grid spacing of 1.0 Å extended 3.0 beyond training set limits was used to the calculate field values. Subsequently, predicted versus measured affinities were plotted to identify outliers (Figure 6). Compounds **80**, **81** and **83** were omitted from the SLK and STK10 models as potential outliers, which increased Q2-value and decreased standard error of prediction. The statistical metrics of models: R-square values with four PLS components were 0.7402, 0.8064 and 0.7900, for GAK, SLK and STK10 models respectively. All three model show acceptable internal predictivities (Q-square values were 0.5967, 0.7303, 0.7631), determined by leave-one-out (LOO) cross-validation. Stability values above 0.97 with four partial least squares regression (PLS) components let us to suggest that the quality and predictivity of the model inside the studied applicability domain was robust.

**Figure 6.**
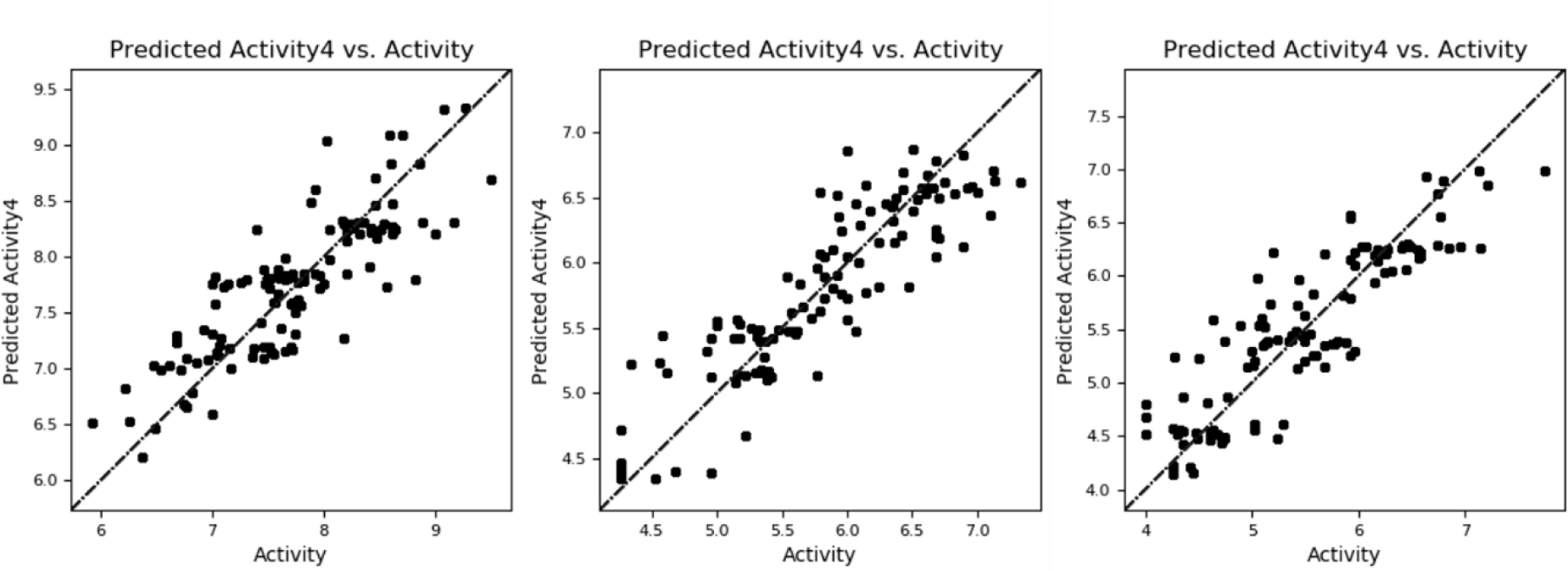
Scatter plots of field based 3D-QSAR models showing predicted versus measured binding affinities (nM) of active compounds printed at logarith-mic scale, from right to left GAK, STK10 and SLK.

**Table 10.** Validation of 3D-QSAR models for GAK

| # Factors | SD | R<sup>2</sup> | R<sup>2</sup> CV | R<sup>2</sup> Scramble | Stability | F | P | RMSE | Q<sup>2</sup> | Pearsson-r |
| --- | --- | --- | --- | --- | --- | --- | --- | --- | --- | --- |
| 1 | 0.5999 | 0.3337 | 0.2685 | 0.0486 | 0.995 | 45.6 | 1.34e-009 | 0.49 | 0.5254 | 0.7440 |
| 2 | 0.5162 | 0.5121 | 0.4160 | 0.0907 | 0.989 | 47.2 | 9.46e-015 | 0.49 | 0.5244 | 0.7301 |
| 3 | 0.4432 | 0.6443 | 0.5188 | 0.1265 | 0.975 | 53.7 | 6.43e-020 | 0.46 | 0.5662 | 0.7537 |
| 4 | 0.3809 | 0.7402 | 0.6201 | 0.1619 | 0.971 | 62.7 | 5.91e-025 | 0.45 | 0.5967 | 0.7973 |
| 5 | 0.3563 | 0.7753 | 0.6624 | 0.1973 | 0.977 | 60.0 | 9.63e-027 | 0.44 | 0.6036 | 0.7971 |
| 6 | 0.3419 | 0.7954 | 0.6613 | 0.2238 | 0.967 | 55.7 | 1.47e-027 | 0.46 | 0.5746 | 0.7963 |

**Table 11.** Validation of 3D-QSAR models for SLK

| # Factors | SD | R<sup>2</sup> | R<sup>2</sup> CV | R<sup>2</sup> Scramble | Stability | F | P | RMSE | Q<sup>2</sup> | Pearsson-r |
| --- | --- | --- | --- | --- | --- | --- | --- | --- | --- | --- |
| 1 | 0.5877 | 0.4180 | 0.3563 | 0.0448 | 0.995 | 63.9 | 4.46e-012 | 0.63 | 0.3679 | 0.6149 |
| 2 | 0.4520 | 0.6595 | 0.6164 | 0.0824 | 0.997 | 85.2 | 2.58e-021 | 0.54 | 0.5467 | 0.7532 |
| 3 | 0.3797 | 0.7625 | 0.7229 | 0.1125 | 0.997 | 93.1 | 4.54e-027 | 0.46 | 0.6588 | 0.8133 |
| 4 | 0.3448 | 0.8064 | 0.7284 | 0.1433 | 0.987 | 89.6 | 7.75e-030 | 0.41 | 0.7303 | 0.8569 |
| 5 | 0.3337 | 0.8208 | 0.7423 | 0.1669 | 0.985 | 77.8 | 3.05e-030 | 0.45 | 0.6843 | 0.8276 |
| 6 | 0.3175 | 0.8397 | 0.7471 | 0.1918 | 0.974 | 73.3 | 2.73e-031 | 0.48 | 0.6427 | 0.8024 |

**Table 12.** Validation of 3D-QSAR models for STK10

| # Factors | SD | R<sup>2</sup> | R<sup>2</sup> CV | R<sup>2</sup> Scramble | Stability | F | P | RMS E | Q<sup>2</sup> | Pearsson-r |
| --- | --- | --- | --- | --- | --- | --- | --- | --- | --- | --- |
| 1 | 0.5556 | 0.5875 | 0.5509 | 0.0627 | 0.998 | 128.2 | 5.36e-019 | 0.45 | 0.7312 | 0.8643 |
| 2 | 0.4327 | 0.7525 | 0.7178 | 0.0861 | 0.998 | 135.3 | 1.03e-027 | 0.42 | 0.7604 | 0.8764 |
| 3 | 0.4092 | 0.7813 | 0.7409 | 0.1226 | 0.997 | 104.8 | 6.06e-029 | 0.44 | 0.7469 | 0.8660 |
| 4 | 0.4032 | 0.7900 | 0.7398 | 0.1637 | 0.995 | 81.8 | 1.16e-028 | 0.42 | 0.7631 | 0.8765 |
| 5 | 0.3798 | 0.8158 | 0.7186 | 0.1850 | 0.978 | 76.2 | 4.23e-030 | 0.39 | 0.7993 | 0.8968 |
| 6 | 0.3769 | 0.8207 | 0.7182 | 0.2127 | 0.976 | 64.8 | 1.25e-029 | 0.37 | 0.8174 | 0.9075 |

Placement of positive steric QSAR fields supports the role of the methoxy groups pointing to the back pocket of the solvent expose region and providing perfect shape complementarity (Figure 7**-**10). Protein structures of the GAK that in WaterMap simulations a high energy water (marked with W1, Figure 11) is found at the top of back pocket/against beta sheet.^40^ The replacement of this water is common among high affinity compounds. This gains additional support from hydrophobic QSAR fields (Figure 9), showing the *meta*-position on the aniline ring should be occupied with this substituent. Interestingly, negative hydrophobic field is in good agreement with oxygen positions of methoxy substituents. Taken together positive steric field (Figure 8) and negative hydrophobic field in (Figure 9) can further explain why triple methoxy compounds are good GAK binders.

**Figure 7.**
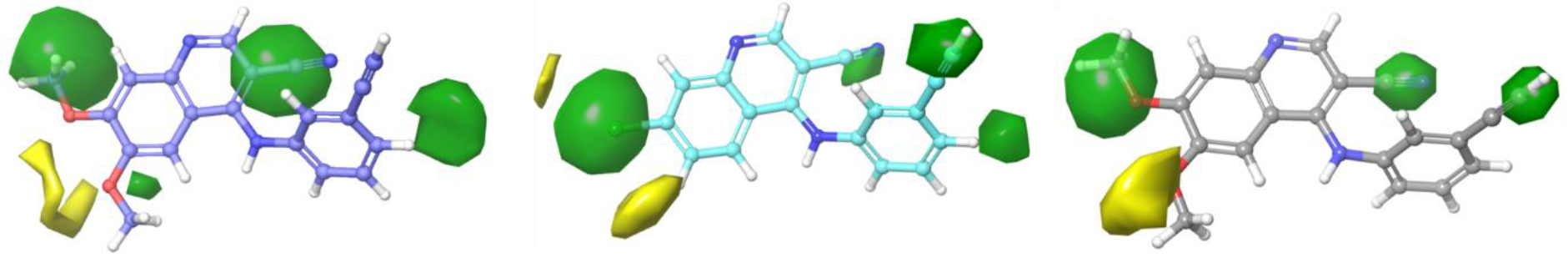
Mains steric fields where green is positive and yellow negative.

**Figure 8.**
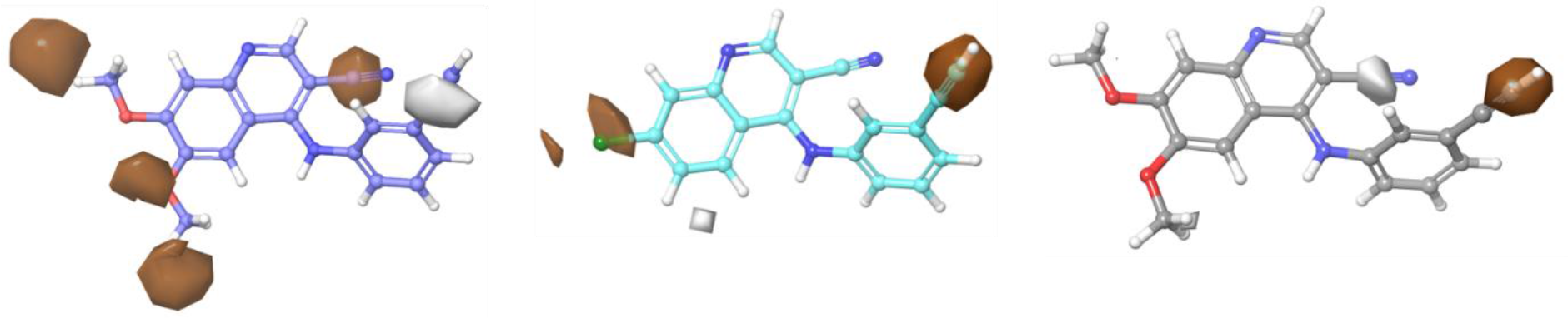
Main Hydrophobic fields where brown is positive and gray negative.

**Figure 9.**
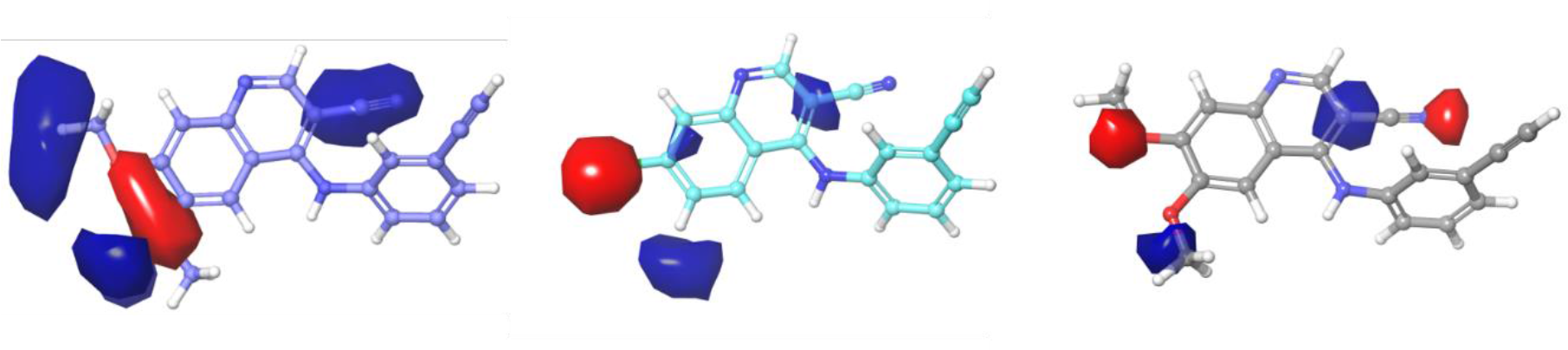
Main electrostatic fields where red stands for negative and blue positive.

**Figure 10.**
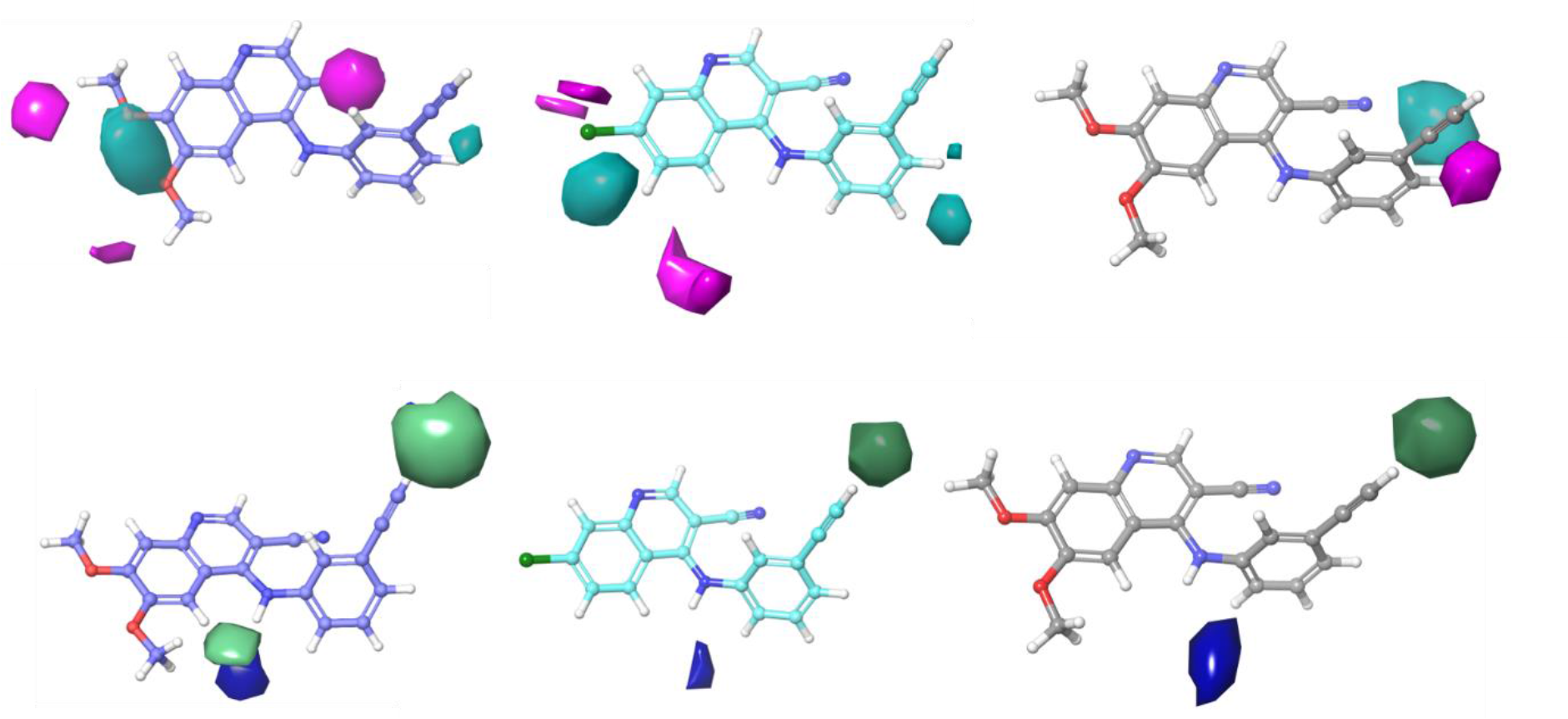
At top row HB acceptor fields where cyan is positive and magenta negative. At the bottom row HB donor fields where blue is positive and green negative.

**Figure 11.**
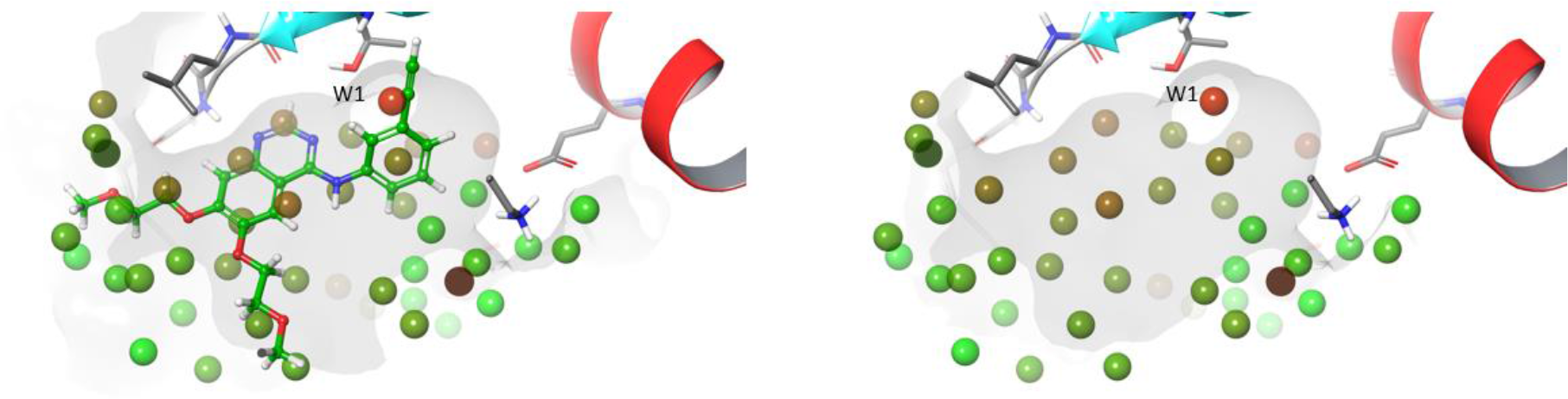
WaterMap analysis of GAK (PDB:5Y80): Showing high energy water W1 at the top of hydrophobic ATP pocket.

Electrostatic fields from QSAR (Figure 10) confirms that the quinoline scaffold is better binder than the quinazoline. The quinazolines provide a negatively charged area, where a positive field is required according to the QSAR model of GAK and leaves also favorable steric field empty (Figure 8). The hydrogen bond acceptor fields shown in figure 10 indicate that the space next to position 7 of the quin(az)oline would also not result in an affinity improvement if filled with hydrogen bond acceptor, however a nitrile group which is significantly bulkier could be tolerated.

WaterMap analysis of the GAK ATP binding site with **82** shows that water occupying site (marked as W3, Figure 12) has less favorable position by means of ΔG. Waters at site W3 serve as hydrogen bond bridge between nitrile group of **82**. The water W3 has HB contact to water bound close to DFG residues, but it is not able to make hydrogen bonds patterns to the bottom back pocket. Thus it appears that water W3 is trapped between ligand and hydrophobic pocket forming an bridged interplay between these residues (Figure 13). The ideal fit of **82** as ligand, results from optimal shape complementarity, high rigidity (favorable entropy upon binding) and replacement of some high energy waters including W1.

**Figure 12.**
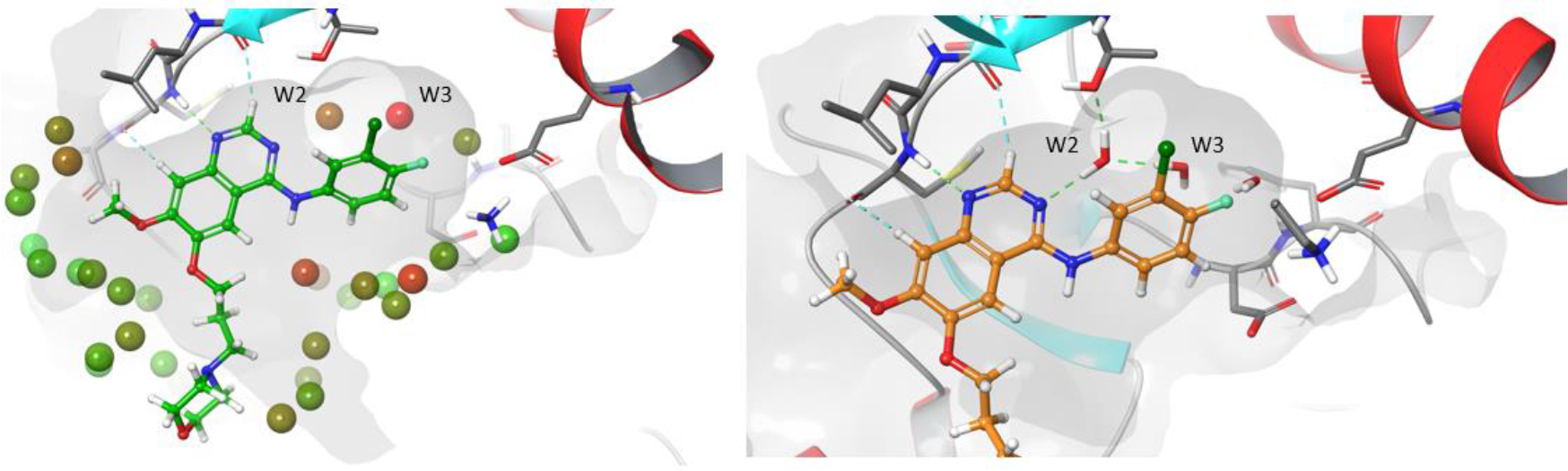
WaterMap analysis of GAK (PDB:5Y80) with preserved co-crytallized ligand shows that bridged water between ligand and protein marked with W2 is stabilizing binging of quinazoline scaffold. On right selected bridging waters taken from hydrogen bond optimized x-ray structure shown. Interestingly, water in site W3 becomes higher in energy compared to empty binding site (Figure 11).

**Figure 13.**
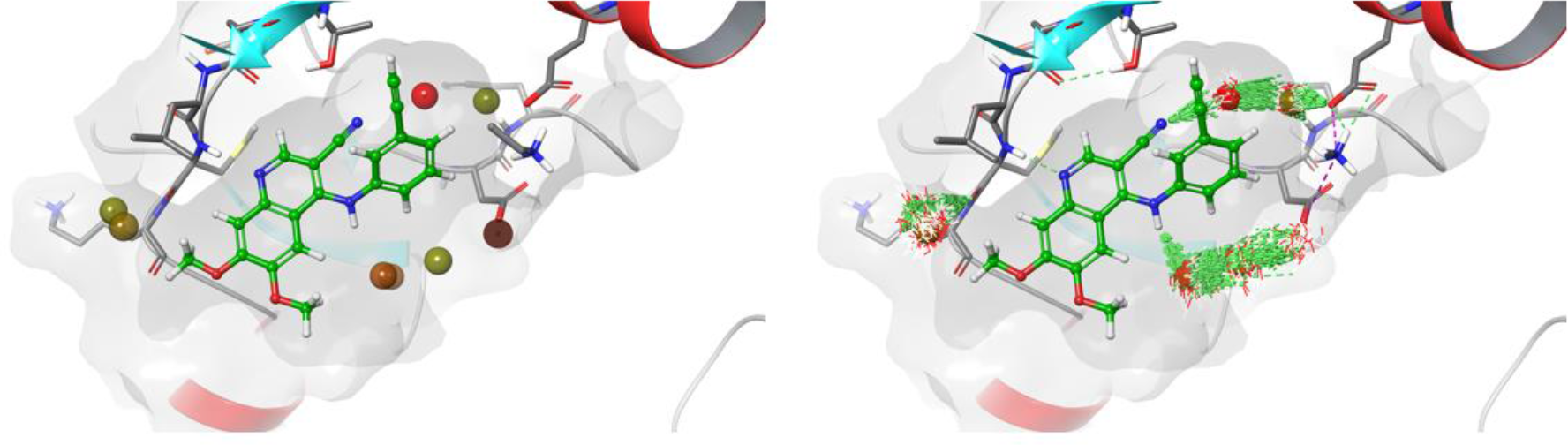
Watermap analysis of active site of GAK (PDB:5Y80) with retained **82** ligand showing that water in site W3 is less favorable for binding. Water W3 serves as hydrogen bond bridged between nitrile group of **82** and water bound close to DFG residues, but it is not able to make hydrogen bonds patterns to the bottom back pocket and is trapped between ligand and hydrophobic pocket.

## DISCUSSION

More than 40 drugs have been approved to target the ATP binding site, mainly in oncology. Although these were developed with specific targets, in practice most of these drugs were multi-kinase inhibitors that leverage ATP pocket across multiple kinases to increase their clinical efficacy. Outside of oncology, selective kinase inhibition profiles are required for treatment of chronic diseases with kinase targets, including inflammation and neurodegeneration. These compounds require enhanced selectivity profiles in order to avoid potential toxicity indications associated with broad kinome inhibition.^43–44^

New chemical approaches and molecular insight into the development of highly selective kinase inhibitors that target the conserved ATP binding site essential if we are to facilitate this unmet clinical need. Binding assays are an accurate and robust method to measure potency and selectivity of ATP-competitive kinase inhibitors.^45^ Binding assays are available at several commercial vendors and are routinely used to profile kinase inhibitors for their selectivity across the human kinome.^7^ Ligand binding displacement assays provide an accepted for direct measurement of kinase inhibition in drug optimization of ATP binding site inhibitors.^46^ This point is particularly relevant for neglected kinases including such as GAK where there are currently no robust and validated enzyme activity assay.

Despite the original target for the 4-anilino-quin(az)oline scaffold being EGFR, SLK/STK10/GAK have been shown to be consistent tractable collateral targets for inhibition.^6–8^ The levels of activity vary for compound to compound and there is not a linear relationship. Erlotinib and vandetanib kinome profiles suggest that there is a potential to find a route to separate these activities. We found the that structural features of the scaffold were key: 1) The cone angle; 2) Displacement of the high energy water and 3) The electronics and sterics of the aniline substitution.

Conformational restriction is an effective tactic in follow-on drug discovery and rational design.^47^ It has been used in several medicinal chemistry programs including the development of a selective CB2 cannabinoid receptor^48^ and on the human histamine H4 receptor (hH4R).^49^ This methodology has shown promise in kinase design by pre-organising the inhibitors in a preferred conformation. This can be used to increase potency on target and enhance selectivity over other kinases/targets. Cardiac troponin I-interacting kinase (TNNI3K)^50–51^ and Spleen tyrosine kinase (Syk)^52^ have both utilised this to increase potency (by 60-160-fold) and selectivity by the addition of a nitrogen (**Figure S1**).

Adding a nitrogen has been proven to have a significant effect on activities across different systems, not only by altered conformation but potentially reduced de-solvation energy potential.^53^ In the 4-anilino-quin(az)oline ring system with GAK we found the opposite observation where going from C-H to nitrogen increases activity, highlighting that importance of both conformation and electronics. (Figure 14). The addition of the nitrogen has allowed for a planar conformation, but the GAK ATP competitive active site prefers a 60-90° 4-anilino-quinoline. The favorable cone angle of the 4-anilino-quinoline laying between to 45-60° is needed for optimal to GAK binding and resulting in increased rigidity along amine linkage with a hierarchy of 3-cyanoquinoline > quinoline > quinazoline.

**Figure 14.**
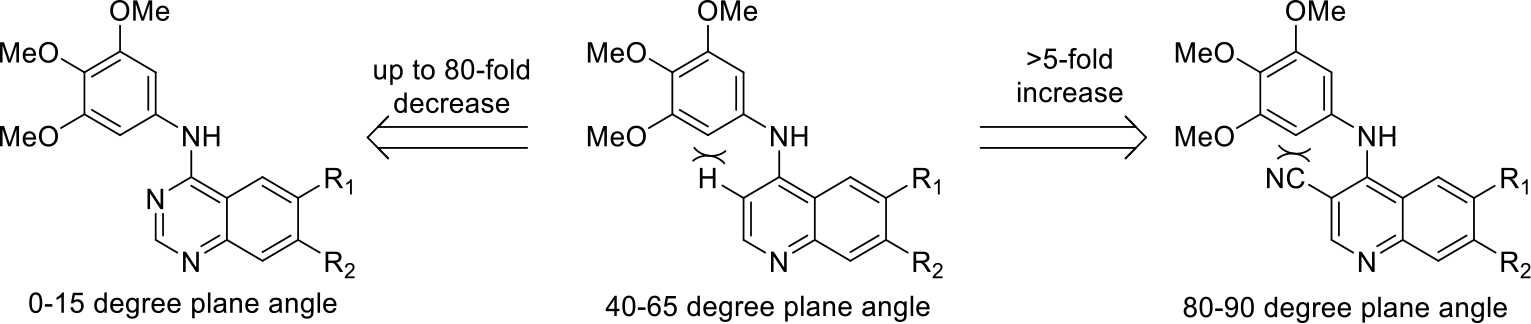
GAK inhibitor conformational sensitive to increased ring rigidity.

There are extensive examples of quinoline and quinazoline in the literature.^6–8^ Our modelling highlights that the quinoline scaffolds seems to have favourable binding to hydrophobic back pocket, whereas quinazoline scaffold seem to require favourable interaction to bridging structural water inside the binding cavity to reach high affinity. The 3-cyanoquinoline, requires more space than quinoline/quinazoline, and according to several experimental co-crystal structures, including bosutinib in Src and Pelitinib in PKMYT1;^54–55^ the bulky cyano group is able form directional hydrogen bond contacts with waters or polar side chains at the hinge binder region. Docking simulations of GAK suggest that favourable effects of 3-cyanoquinoline allow it to completely fill the available space at back pocket; in addition to providing a conformational rigidity to the scaffold.

The small molecule crystal structures (Figure 15) support the theoretical observations with the GAK ATP binding site. The quinazoline is consistently >10-fold less potent for GAK with a maximum of short of 80-fold for **21** vs **36**. The 3-cyanoquinoline is consistently 5-15-fold more potent than the quinoline for GAK. The switch from trimethoxy to 3-actylene show similar trends with regards to GAK activity. This is likely due to the ability of both substitution patterns been able to displace the high energy water W1 in the GAK hydrophobic pocket (Figure 11).^13^ Comparison of WaterMap sites between GAK, STK10 and SLK with representative 4-anilinoquin(az)oline ligands suggests that the high energy water is not able to be accessed easily in the case of SLK and STK10 (Figure 15). Indeed, according to the docking of the 4-anilinoquin(az)olines into SLK the molecules prefer a flipped binding mode preventing effective manipulation of the water network seen in GAK (Figure 15).

**Figure 15.**
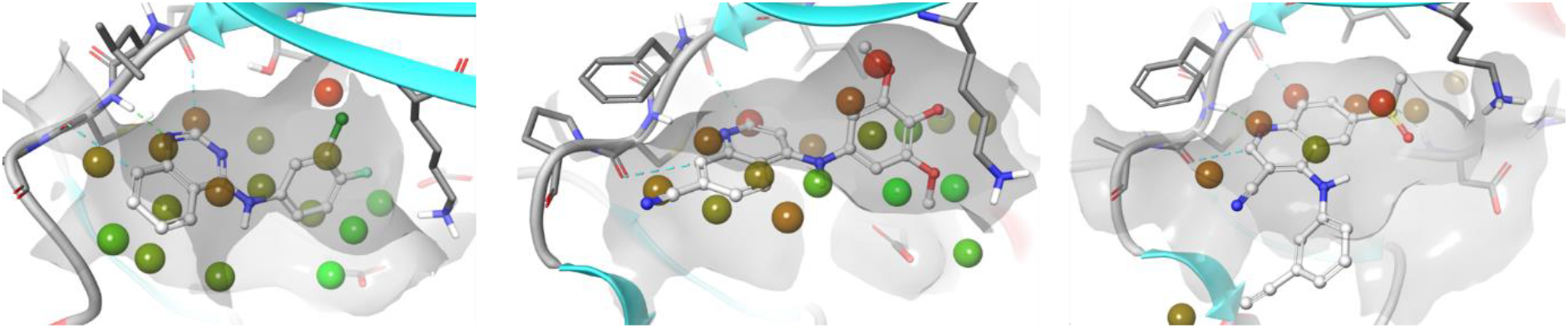
Comparison of WaterMap sites between GAK, STK10 and SLK with selected ligands. According to docking SLK binds quinoline scaffold in flipped geometry.

## Conclusions

While literature inhibitors show good cellular target engagement for GAK and activity and there was a good concordance between nanoBRET assay and GAK K_i_. We were able to design and synthesize a compound that was 0.47-fold SLK/STK10 selective over GAK on a scaffold that has shown consistently high potency towards GAK. The landscape of kinase drug discovery is complex, and the identification of highly selective inhibitors is still a significant challenge.^56^ Our QSAR models for GAK, SLK and STK10 provide new insights into the development of selective chemical probes for SLK and STK10 utilising the 4-anilinoquin(az)oline scaffold.

### Modelling

*Structures of small molecules* were prepared using the LigPrep module of Schrodinger suite (Schrödinger Release 2018-4: Schrödinger, LLC, New York, NY, 2018) employing OPLS3e force field. Prior to field bases QSAR studies the ligand structures were overlaid with co-crystallized ligands or favorable docking poses using Flexible ligand alignment tool employing a Bemis-Mucko method,^42^ Torsion angles of larger substituents, such as methoxy groups, were manually adjusted to fit template structures when needed.

*3D-QSAR models* were constructed using the field based QSAR functionality of Schrödinger Maestro 2018-4.

*Protein preparation.* The experimental structures of studied kinases were obtained from RCSB database (PDB: 5Y80/GAK, 5AJQ/STK10, 6HVD/SLK). Structures were processed with Protein preparation wizard of Schrodinger Maestro suite 2018-4 (Schrödinger Release 2018-4) in order to optimize the hydrogen bonding network followed by constrained minimization (heavy atom rmsd converge of 0.3 Å) to remove crystallographic artifacts.

*Docking Simulations* of three kinases. In the definition process of the docking parameter file, water molecules were removed from prepared protein and the center of grid box was positioned to the center of adenine pocket binding part of co-crystallized ligand.^57^ Finally, h-bond constraints were added to hinge region mainchain amide C126/GAK, C111/SLK, C113/STK10) to improve the convergence.

*Hydration site Analysis* were calculated with WaterMap (Schrödinger Release 2018-4: WaterMap, Schrödinger, LLC, New York, NY, 2018.). The protein structures were prepared with Protein Preparation Wizard (as above). Waters were analyzed within 6 Å from the favorable docking pose of selected high affinity ligands. 2 ns simulations were conputed with OPLS3e force field.

*Graphical illustrations*. PyMol v.2.3.0 and Maestro (Schrödinger release 2018-4) were used for visualization of results.

### Biology

GAK nanoBRET screening was performed as previously described.^13^

Ligand Binding Displacement Assay was performed as previously described,^13^ Constructs used: AAK1 - AAKA-p051; BMPK2K - BMP2KA-p031; GAK - GAKA-p059; STK16 - STK16A-p016; SLK - SLKA-p026; STK10 - STK10A-p047.^13^

### Chemistry Experimental

All reactions were performed using flame-dried round-bottomed flasks or reaction vessels unless otherwise stated. Where appropriate, reactions were carried out under a nitrogen atmosphere with dry solvents, unless otherwise stated. Yields refer to chromatographically and spectroscopically pure isolated yields. Reagents were purchased at the highest commercial quality and used without further purification, unless otherwise stated. Reactions were monitored by thin-layer chromatography carried out on 0.25 mM E. Merck silica gel plates (60_F-254_) using ultraviolet light as visualizing agent. NMR spectra were recorded on a Varian Inova 400 or Inova 500 spectrometer and were calibrated using residual protic solvent as an internal reference (CDCl_3_: ^1^H NMR = 7.26, ^13^C NMR = 77.16). The following abbreviations or combinations thereof were used to explain the multiplicities observed: s = singlet, d = doublet, t = triplet, q = quartet, m = multiplet, br = broad. Liquid chromatography (LC) and high resolution mass spectra (HRMS) were recorded on a ThermoFisher hybrid LTQ FT (ICR 7T). The University of Southampton (Southampton, UK) small molecule x-ray facility collected and analyzed all X-ray diffraction data. Compounds (**1**, **42**, **48**, **62**, **71)**^11^, (**2**, **10**, **11**)^12^, (**3**-**8**, **12**-**16**, **20**-**22**, **30**, **40**)^13^ (**17**-**19**)^58^, (**70**, **72**, **81**-**83**)^38^ were synthesised as previously reported.

**General Procedure for the Synthesis of 4-Anilinoquinolines (**150 mg scale unless otherwise stated). 6-Bromo-4-chloroquinoline (1.0 equiv), aniline (1.1 equiv), and i-Pr_2_NEt (2.5 equiv) were suspended in ethanol (30 mL) and refluxed for 18 h. The crude mixture was purified by flash chromatography using EtOAc/hexane followed by 1−5 % methanol in EtOAc (or by re-crystalisation). After solvent removal under reduced pressure, the product was obtained. All compounds were >98% pure by ^1^H/^13^C NMR and LCMS.

### Compound Characterization

**6,7-difluoro-*N*-(3,4,5-trimethoxyphenyl)quinolin-4-amine** (**7**) was obtained as a green solid (137 mg, 0.396 mmol, 53 %). MP 282-284 °C; ^1^H NMR (400 MHz, DMSO-*d*_6_) δ 11.10 (s, 1H), 9.10 (dd, *J* = 12.2, 8.1 Hz, 1H), 8.51 (d, *J* = 7.0 Hz, 1H), 8.17 (dd, *J* = 10.8, 7.3 Hz, 1H), 6.90 (d, *J* = 7.0 Hz, 1H), 6.81 (s, 2H), 3.80 (s, 6H), 3.72 (s 3H). ^13^C NMR (101 MHz, DMSO-*d*_6_) δ 154.6 (d, *J* = 3.5 Hz), 153.6 (s, 2C), 152.9 (dd, *J* = 256.7, 15.9 Hz), 148.5 (dd, *J* = 249.1, 14.6 Hz), 143.3, 136.6, 136.4 (d, *J* = 10.7 Hz), 132.5, 114.3 (d, *J* = 7.1 Hz), 111.9 (d, *J* = 20.6 Hz), 108.2 (d, *J* = 20.7 Hz), 103.1 (s, 2C), 100.5, 60.2, 56.1 (s, 2C). HRMS *m/z* [M+H]^+^ calcd for C_18_H_17_N_2_O_3_F_2_: 347.1207, found 347.1197, LC *t*_R_ = 3.26 min, >98% Purity.

***N*-(3,4,5-trimethoxyphenyl)quinazolin-4-amine** (**23**) was obtained as a yellow solid (247 mg, 0.793 mmol, 87 %). MP 223-225 °C; ^1^H NMR (400 MHz, DMSO-*d*_6_) δ 11.78 (s, 1H), 9.00 (d, *J* = 8.4 Hz, 1H), 8.94 (s, 1H), 8.11 (ddd, *J* = 8.3, 7.0, 1.2 Hz, 1H), 8.01 (dd, *J* = 8.4, 1.2 Hz, 1H), 7.86 (ddd, *J* = 8.4, 7.0, 1.2 Hz, 1H), 7.16 (s, 2H), 3.80 (s, 6H), 3.71 (s, 3H). ^13^C NMR (101 MHz, DMSO-*d*_6_) δ 159.8, 152.7 (s, 2C), 150.9, 138.4, 136.2, 136.2, 132.4, 128.6, 124.9, 119.6, 113.4, 103.0 (s, 2C), 60.2, 56.1 (s, 2C). HRMS *m/z* [M+H]^+^ calcd for C_17_H_18_N_3_O_3_: 312.1348, found 312.1335, LC *t*_R_ = 2.85 min, >98% Purity.

**6-methyl-*N*-(3,4,5-trimethoxyphenyl)quinazolin-4-amine** (**24**) was obtained as a green solid (147 mg, 0.451 mmol, 54 %). MP 198-200 °C; ^1^H NMR (400 MHz, DMSO-*d*_6_) δ 11.43 (s, 1H), 8.90 (s, 1H), 8.74 (s, 1H), 7.95 (dd, *J* = 8.6, 1.6 Hz, 1H), 7.88 (d, *J* = 8.5 Hz, 1H), 7.17 (s, 2H), 3.80 (s, 6H), 3.71 (s, 3H), 2.57 (s, 3H). ^13^C NMR (101 MHz, DMSO-*d*_6_) δ 159.3, 152.7 (s, 2C), 150.3, 138.8, 137. 6, 136.9, 136.0, 132.6, 123.6, 119.7, 113.4, 102.7 (s, 2C), 60.2, 56.1 (s, 2C), 21.2. HRMS *m/z* [M+H]^+^ calcd for C_18_H_20_N_3_O_3_: 326.1505, found 326.1491, LC *t*_R_ = 3.26 min, >98% Purity.

**6-fluoro-*N*-(3,4,5-trimethoxyphenyl)quinazolin-4-amine** (**25**) was obtained as a yellow solid (195 mg, 0.592 mmol, 72 %). MP 230-232 °C; ^1^H NMR (400 MHz, DMSO-*d*_6_) δ 11.64 (s, 1H), 8.96 (d, *J* = 2.6 Hz, 1H), 8.94 (s, 3H), 8.08 (dd, *J* = 9.2, 5.1 Hz, 1H), 8.03 (ddd, *J* = 9.2, 8.0, 2.6 Hz, 1H), 7.19 (s, 2H), 3.80 (s, 6H), 3.70 (s, 3H).^13^C NMR (101 MHz, DMSO-*d*_6_) δ 160.3 (d, *J* = 247.5 Hz), 159.3 (d, *J* = 3.6 Hz), 152.7 (s, 2C), 150.8, 136.2, 136.1, 132.4, 125.1 (d, *J* = 25.4 Hz), 123.2 (d, *J* = 9.2 Hz), 114.9 (d, *J* = 9.5 Hz), 110.0 (d, *J* = 25.4 Hz), 102.6 (s, 2C), 60.2, 56.1 (s, 2C). HRMS *m/z* [M+H]^+^ calcd for C_17_H_17_FN_3_O_3_: 330.1254, found 330.1240, LC *t*_R_ = 3.03 min, >98% Purity.

**6,7-difluoro-*N*-(3,4,5-trimethoxyphenyl)quinazolin-4-amine** (**26**) was obtained as a green solid (164 mg, 0.471 mmol, 64 %). MP 281-283 °C; ^1^H NMR (400 MHz, DMSO-*d*_6_) δ 11.29 (s, 1H), 9.11 (dd, *J* = 11.6, 8.1 Hz, 1H), 8.88 (s, 1H), 7.98 (dd, *J* = 10.7, 7.4 Hz, 1H), 7.18 (s, 2H), 3.80 (s, 6H), 3.70 (s, 3H). ^13^C NMR (101 MHz, DMSO-*d*_6_) δ 158.5 (d, *J* = 2.3 Hz), 156.2 – 152.6 (m, 1C), 152.7 (s, 2C), 152.4, 149.0 (dd, *J* = 250.0, 14.4 Hz), 139.7, 135.8, 132.7, 112.7 (d, *J* = 20.7 Hz), 111.1 (dd, *J* = 7.4, 1.7 Hz), 109.9 (d, *J* = 20.2 Hz), 102.2 (s, 2C), 60.2, 56.0 (s, 2C). HRMS *m/z* [M+H]^+^ calcd for C_17_H_16_N_3_O_3_F_2_: 348.1160, found 348.1151, LC *t*_R_ = 3.85 min, >98% Purity.

**6-chloro-*N*-(3,4,5-trimethoxyphenyl)quinazolin-4-amine** (**27**) was obtained as a green solid (203 mg, 0.588 mmol, 78 %). MP 221-223 °C; ^1^H NMR (400 MHz, DMSO-*d*_6_) δ 11.69 (s, 1H), 9.17 (d, *J* = 2.2 Hz, 1H), 8.94 (s, 1H), 8.13 (dd, *J* = 9.0, 2.1 Hz, 1H), 8.03 (d, *J* = 8.9 Hz, 1H), 7.18 (s, 2H), 3.80 (s, 6H), 3.70 (s, 3H). ^13^C NMR (101 MHz, DMSO-*d*_6_) δ 158.7, 152.7 (s, 2C), 151.2, 137.9, 136.1, 132.6 (s, 2C), 132.4, 124.2, 122.2, 114.7, 102.6 (s, 2C), 60.2, 56.1 (s, 2C). HRMS *m/z* [M+H]^+^ calcd for C_17_H_17_N_3_O_3_Cl: 346.0958, found 346.0947, LC *t*_R_ = 3.50 min, >98% Purity.

**6-bromo-*N*-(3,4,5-trimethoxyphenyl)quinazolin-4-amine** (**28**) was obtained as a green solid (214 mg, 0.548 mmol, 89 %). MP 237-239 °C; ^1^H NMR (400 MHz, DMSO-*d*_6_) δ 11.66 (s, 1H), 9.27 (d, *J* = 2.0 Hz, 1H), 8.95 (s, 1H), 8.24 (dd, *J* = 8.9, 2.0 Hz, 1H), 7.95 (d, *J* = 8.9 Hz, 1H), 7.17 (s, 2H), 3.80 (s, 6H), 3.70 (s, 3H). ^13^C NMR (101 MHz, DMSO-*d*_6_) δ 158.6, 152.7 (s, 2C), 151.2, 138.7, 138.2, 136.1 (s, 2C), 132.4, 127.2, 122.3, 120.9, 115.1, 102.6, 60.2, 56.1 (s, 2C). HRMS *m/z* [M+H]^+^ calcd for C_17_H_17_BrN_3_O_3_: 390.0453, found 390.0447, LC *t*_R_ = 3.74 min, >98% Purity.

**6-iodo-*N*-(3,4,5-trimethoxyphenyl)quinazolin-4-amine** (**29**) was obtained as a yellow solid (160 mg, 0.366 mmol, 71 %). MP 226-228 °C; ^1^H NMR (400 MHz, DMSO-*d*_6_) δ 11.68 (s, 1H), 9.39 (d, *J* = 1.7 Hz, 1H), 8.94 (s, 1H), 8.36 (dd, *J* = 8.7, 1.7 Hz, 1H), 7.79 (d, *J* = 8.7 Hz, 1H), 7.16 (s, 2H), 3.79 (s, 6H), 3.70 (s, 3H). ^13^C NMR (101 MHz, DMSO-*d*_6_) δ 158.3, 152.6 (s, 2C), 151.0, 144.1, 138.2, 136.1 (s, 2C), 133.0, 132.4, 121.5, 115.1, 102.7, 94.2, 60.2, 56.1 (s, 2C). HRMS *m/z* [M+H]^+^ calcd for C_17_H_17_N_3_O_3_I: 438.0315, found 438.0292, LC *t*_R_ = 3.80 min, >98% Purity.

**7-fluoro-*N*-(3,4,5-trimethoxyphenyl)quinazolin-4-amine** (**31**) was obtained as a yellow solid (216 mg, 0.656 mmol, 80 %). MP 246-248 °C; ^1^H NMR (400 MHz, DMSO-*d*_6_) δ 11.73 (s, 1H), 9.08 (dd, *J* = 9.2, 5.4 Hz, 1H), 8.93 (s, 1H), 7.88 – 7.71 (m, 2H), 7.14 (s, 2H), 3.79 (s, 6H), 3.70 (s, 3H). ^13^C NMR (101 MHz, DMSO-*d*_6_) δ 165.5 (d, *J* = 255.9 Hz), 159.2, 152.7 (s, 2C), 151.9, 141.3 (d, *J* = 13.6 Hz), 136.1 (s, 2C), 132.4, 128.7 (d, *J* = 10.7 Hz), 117.6 (d, *J* = 23.9 Hz), 110.7 (d, *J* = 1.8 Hz), 105.6 (d, *J* = 24.9 Hz), 102.9, 60.2, 56.1 (s, 2C). HRMS *m/z* [M+H]^+^ calcd for C_17_H_17_N_3_O_3_F: 330.1254, found 330.1240, LC *t*_R_ = 3.02 min, >98% Purity.

**7-chloro-*N*-(3,4,5-trimethoxyphenyl)quinazolin-4-amine** (**32**) was obtained as a yellow solid (159 mg, 0.460 mmol, 61 %). MP 230-232 °C; ^1^H NMR (400 MHz, DMSO-*d*_6_) δ 11.54 (s, 1H), 8.90 (d, *J* = 8.3 Hz, 2H), 8.01 (d, *J* = 2.1 Hz, 1H), 7.93 (dd, *J* = 8.9, 2.1 Hz, 1H), 7.14 (s, 2H), 3.80 (s, 6H), 3.70 (s, 3H). ^13^C NMR (101 MHz, DMSO-*d*_6_) δ 159.2, 152.7 (s, 2C), 152.2, 141.0, 140.3, 136.0, 132.5, 128.7, 126.9, 119.9, 112.5, 102.6 (s, 2C), 60.2, 56.1 (s, 2C). HRMS *m/z* [M+H]^+^ calcd for C_17_H_17_N_3_O_3_Cl: 346.0958, found 346.0947, LC *t*_R_ = 3.44 min, >98% Purity.

**7-bromo-*N*-(3,4,5-trimethoxyphenyl)quinazolin-4-amine** (**33**) was obtained as a yellow solid (125 mg, 0.320 mmol, 52 %). MP 233-235 °C; ^1^H NMR (400 MHz, DMSO-*d*_6_) δ 11.57 (s, 1H), 8.91 (s, 1H), 8.82 (d, *J* = 8.9 Hz, 1H), 8.16 (d, *J* = 2.0 Hz, 1H), 8.05 (dd, *J* = 8.9, 2.0 Hz, 1H), 7.14 (s, 2H), 3.80 (s, 6H), 3.70 (s, 3H). ^13^C NMR (101 MHz, DMSO-*d*_6_) δ 159.3, 152.7 (s, 2C), 152.0, 140.8, 136.1, 132.5, 131.4, 129.4, 126.7, 122.9, 112.8, 102.61 (s, 2C), 60.2, 56.1 (s, 2C). HRMS *m/z* [M+H]^+^ calcd for C_17_H_17_N_3_O_3_Br: 390.0453, found 390.0445, LC *t*_R_ = 3.61 min, >98% Purity.

**7-iodo-*N*-(3,4,5-trimethoxyphenyl)quinazolin-4-amine** (**34**) was obtained as a yellow solid (120 mg, 0.274 mmol, 53 %). MP 246-248 °C; ^1^H NMR (400 MHz, DMSO-*d*_6_) δ 11.68 (s, 1H), 8.90 (s, 1H), 8.67 (d, *J* = 8.8 Hz, 1H), 8.37 (d, *J* = 1.6 Hz, 1H), 8.18 (dd, *J* = 8.7, 1.7 Hz, 1H), 7.14 (s, 2H), 3.79 (s, 6H), 3.70 (s, 3H). ^13^C NMR (101 MHz, DMSO-*d*_6_) δ 159.6, 152.7 (s, 2C), 151.4, 139.7, 137.0, 136.1, 132.4, 128.3, 126.0, 112.8, 104.8, 102.7 (s, 2C), 60.2, 56.1 (s, 2C). HRMS *m/z* [M+H]^+^ calcd for C_17_H_17_N_3_O_3_I: 438.0315, found 438.0300, LC *t*_R_ = 3.82 min, >98% Purity.

**7-(trifluoromethyl)-*N*-(3,4,5-trimethoxyphenyl)quinazolin-4-amine** (**35**) was obtained as a green solid (54 mg, 0.142 mmol, 22 %). MP 201-203 °C; ^1^H NMR (400 MHz, DMSO-*d*_6_) δ 11.57 (s, 1H), 9.10 (d, *J* = 8.7 Hz, 1H), 8.97 (s, 1H), 8.32 - 8.24 (m, 1H), 8.16 (dd, *J* = 9.0, 1.8 Hz, 1H), 7.20 (s, 2H), 3.80 (s, 6H), 3.71 (s, 3H). ^13^C NMR (101 MHz, DMSO-*d*_6_) δ 159.0, 153.0, 152.7 (s, 2C), 141.1, 135.9, 134.7 - 133.7 (m, 1C), 132.7, 126.6, 124.4, 123.6 - 123.5 (m, 1C), 121.7, 119.0 - 118.8 (m, 1C), 116.4, 102.4 (s, 2C), 60.2, 56.1 (s, 2C). HRMS *m/z* [M+H]^+^ calcd for C_18_H_17_N_3_O_3_F_3_: 380.1222, found 380.1204, LC *t*_R_ = 4.41 min, >98% Purity.

**4-((3,4,5-trimethoxyphenyl)amino)quinazoline-7-carbonitrile** (**36**) was obtained as a yellow solid (149 mg, 0.443 mmol, 56 %). MP 230-232 °C; ^1^H NMR (400 MHz, DMSO-*d*_6_) δ 11.77 (s, 1H), 9.13 (d, *J* = 8.6 Hz, 1H), 8.95 (s, 1H), 8.42 (d, *J* = 1.6 Hz, 1H), 8.20 (dd, *J* = 8.6, 1.6 Hz, 1H), 7.20 (s, 2H), 3.80 (s, 6H), 3.70 (s, 3H). ^13^C NMR (101 MHz, DMSO-*d*_6_) δ 159.0, 152.8, 152.7 (s, 2C), 140.3, 136.1, 132.5, 129.7, 126.3, 126.0, 117.2, 117.1, 116.5, 102.5 (s, 2C), 60.2, 56.1 (s, 2C). HRMS *m/z* [M+H]^+^ calcd for C_18_H_17_N_4_O_3_: 337.1301, found 337.1288, LC *t*_R_ = 3.67 min, >98% Purity.

**4-((3,4,5-trimethoxyphenyl)amino)quinazoline-6-carbonitrile** (**37**) was obtained as a yellow solid (172 mg, 0.511 mmol, 65 %). MP 239-241 °C; ^1^H NMR (400 MHz, DMSO-*d*_6_) δ 11.58 (s, 1H), 9.49 (d, *J* = 1.7 Hz, 1H), 8.96 (s, 1H), 8.37 (dd, *J* = 8.7, 1.7 Hz, 1H), 8.05 (d, *J* = 8.7 Hz, 1H), 7.20 (s, 2H), 3.81 (s, 6H), 3.71 (s, 3H). ^13^C NMR (101 MHz, DMSO-*d*_6_) δ 158.7, 153.7 (s, 2C), 152.7, 143.2, 136.8, 136.0, 132.5, 130.8, 122.9, 117.8, 114.0, 110.0, 102.2 (s, 2C), 60.2, 56.1 (s, 2C). HRMS *m/z* [M+H]^+^ calcd for C_18_H_17_N_4_O_3_: 337.1301, found 337.1287, LC *t*_R_ = 3.53 min, >98% Purity.

**6-(methylsulfonyl)-*N*-(3,4,5-trimethoxyphenyl)quinazolin-4-amine** (**38**) was obtained as a grey solid (210 mg, 0.539 mmol, 87 %). MP 260-262 °C; ^1^H NMR (400 MHz, DMSO-*d*_6_) δ 11.91 (s, 1H), 9.55 (d, *J* = 1.9 Hz, 1H), 8.98 (s, 1H), 8.49 (dd, *J* = 8.8, 1.9 Hz, 1H), 8.14 (d, *J* = 8.8 Hz, 1H), 7.15 (s, 2H), 3.81 (s, 6H), 3.71 (s, 3H), 3.43 (s, 3H). ^13^C NMR (101 MHz, DMSO-*d*_6_) δ 159.7, 153.3, 152.7 (s, 2C), 142.9, 139.8, 136.1, 132.5, 132.4, 125.3, 122.4, 113.7, 102.6 (s, 2C), 60.2, 56.1 (s, 2C), 43.5. HRMS *m/z* [M+H]^+^ calcd for C_18_H_20_N_3_O_5_S: 390.1124, found 390.1112, LC *t*_R_ = 3.23 min, >98% Purity.

**6-methoxy-*N*-(3,4,5-trimethoxyphenyl)quinazolin-4-amine** (**39**) was obtained as a grey solid (142 mg, 0.416 mmol, 54 %). MP 158-160 °C; ^1^H NMR (400 MHz, DMSO-*d*_6_) δ 11.83 (s, 1H), 8.84 (s, 1H), 8.55 (d, *J* = 2.6 Hz, 1H), 7.95 (d, *J* = 9.1 Hz, 1H), 7.70 (dd, *J* = 9.1, 2.5 Hz, 1H), 7.18 (s, 2H), 4.01 (s, 3H), 3.79 (s, 6H), 3.70 (s, 3H).). ^13^C NMR (101 MHz, DMSO-*d*_6_) δ 159.0, 152.7, 152.6, 148.5, 136.0, 133.1, 132.5, 126.9, 121.7, 121.1, 114.6, 104.9, 104.6, 103.1, 60.2, 56.9, 56.1 (s, 2C). HRMS *m/z* [M+H]^+^ calcd for C_18_H_20_N_3_O_4_: 342.1454, found 342.1440, LC *t*_R_ = 3.17 min, >98% Purity.

**7-methoxy-*N*-(3,4,5-trimethoxyphenyl)quinazolin-4-amine** (**41**) was obtained as a yellow solid (167 mg, 0.489 mmol, 76 %). MP 211-213 °C; ^1^H NMR (400 MHz, DMSO-*d*_6_) δ 11.42 (s, 1H), 8.87 (s, 1H), 8.83 (d, *J* = 9.3 Hz, 1H), 7.48 (dd, *J* = 9.3, 2.5 Hz, 1H), 7.35 (d, *J* = 2.5 Hz, 1H), 7.11 (s, 2H), 3.98 (s, 3H), 3.79 (s, 6H), 3.70 (s, 3H). ^13^C NMR (101 MHz, DMSO-*d*_6_) δ 164.8, 159.0, 152.7 (s, 2C), 151.0, 141.1, 136.0, 132.5, 126.8, 118.9, 107.2, 102.9 (s, 2C), 100.4, 60.2, 56.1, 56.1 (s, 2C). HRMS *m/z* [M+H]^+^ calcd for C_18_H_20_N_3_O_4_: 342.1454, found 342.1440, LC *t*_R_ = 3.08 min, >98% Purity.

**4-((3,4,5-trimethoxyphenyl)amino)quinoline-3-carbonitrile** (**43**) was obtained as a dark orange solid (243 mg, 0.725 mmol, 91%). MP 226-228 °C; ^1^H NMR (400 MHz, DMSO-*d*_6_) δ 11.71 (s, 1H), 9.09 (s, 1H), 8.93 (dd, *J* = 8.7, 1.2 Hz, 1H), 8.17 (dd, *J* = 8.5, 1.3 Hz, 1H), 8.09 (ddd, *J* = 8.4, 7.0, 1.1 Hz, 1H), 7.86 (ddd, *J* = 8.4, 7.0, 1.3 Hz, 1H), 6.88 (s, 2H), 3.78 (s, 6H), 3.71 (s, 3H). ^13^C NMR (101 MHz, DMSO-*d*_6_) δ 154.9, 153.1 (s, 2C), 149.9, 138.5, 137.9, 135.0, 132.5 (s, 2C), 128.2, 124.5, 121.5, 118.2, 113.9, 104.9 (s, 2C), 86.3, 60.3, 56.2 (s, 2C). HRMS *m/z* [M+H]^+^ calcd for C_19_H_18_N_3_O_3_: 336.1348, found 336.1349, LC *t*_R_ = 4.20 min, >98% Purity.

**6-chloro-4-((3,4,5-trimethoxyphenyl)amino)quinoline-3-carbonitrile** (**44**) was obtained as an orange solid (170 mg, 0.460 mmol, 68 %). MP °C; ^1^H NMR (400 MHz, DMSO-*d*_6_) δ 10.91 (s, 1H), 8.92 (s, 1H), 8.86 (t, *J* = 1.3 Hz, 1H), 8.07 – 8.00 (m, 2H), 6.79 (s, 2H), 3.78 (s, 6H), 3.70 (s, 3H). ^13^C NMR (101 MHz, DMSO-*d*_6_) δ 153.6 (s, 2C), 153.2, 152.1, 137.8, 134.4, 134.4, 133.5, 132.6, 127.2, 123.6, 120.1, 115.3, 104.6 (s, 2C), 87.6, 60.75, 56.6 (s, 2C). HRMS *m/z* [M+H]^+^ calcd for C_19_H_17_N_3_O_3_Cl: 370.0958, found 370.0949, LC *t*_R_ = 4.29 min, >98% Purity.

**6-bromo-4-((3,4,5-trimethoxyphenyl)amino)quinoline-3-carbonitrile** (**45**) was obtained as a yellow solid (182 mg, 0.439 mmol, 78 %). MP 204-206 °C; ^1^H NMR (400 MHz, DMSO-*d*_6_) δ 11.54 (s, 1H), 9.19 (d, *J* = 2.1 Hz, 1H), 9.05 (s, 1H), 8.21 (dd, *J* = 8.9, 1.9 Hz, 1H), 8.08 (d, *J* = 9.0 Hz, 1H), 6.84 (s, 2H), 3.78 (s, 6H), 3.70 (s, 3H). ^13^C NMR (101 MHz, DMSO-*d*_6_) δ 153.6, 153.1 (s, 2C), 150.6, 137.7, 137.3 (s, 2C), 1326, 126.8, 124.4, 121.0, 119.8, 114.0, 104.5 (s, 2C), 87.0, 60.3, 56.2 (s, 2C). HRMS *m/z* [M+H]^+^ calcd for C_19_H_17_BrN_3_O_3_: 414.0453, found 414.0444, LC *t*_R_ = 4.49 min, >98% Purity.

**6-iodo-4-((3,4,5-trimethoxyphenyl)amino)quinoline-3-carbonitrile** (**46**) was obtained as a yellow solid (197 mg, 0.427 mmol, 90 %). MP 176-178 °C; ^1^H NMR (400 MHz, DMSO-*d*_6_) δ 11.46 (s, 1H), 9.27 (d, *J* = 1.8 Hz, 1H), 9.04 (s, 1H), 8.33 (dd, *J* = 8.8, 1.7 Hz, 1H), 7.89 (d, *J* = 8.8 Hz, 1H), 6.84 (s, 2H), 3.78 (s, 6H), 3.70 (s, 3H). ^13^C NMR (101 MHz, DMSO-*d*_6_) δ 153.3, 153.1 (s, 2C), 150.5, 142.7, 138.9, 137.7 (s, 2C), 132.6, 120.0, 114.1, 104.5 (s, 2C), 94.2, 87.0, 60.3, 56.2 (s, 2C). HRMS *m/z* [M+H]^+^ calcd for C_19_H_17_N_3_O_3_I: 462.0314, found 462.0303, LC *t*_R_ = 4.62 min, >98% Purity.

**6-methoxy-4-((3,4,5-trimethoxyphenyl)amino)quinoline-3-carbonitrile** (**47**) was obtained as a yellow solid (196 mg, 0.536 mmol, 78 %). MP 220-222 °C; ^1^H NMR (400 MHz, DMSO-*d*_6_) δ 11.36 (s, 1H), 8.98 (s, 1H), 8.26 (d, *J* = 2.6 Hz, 1H), 8.05 (d, *J* = 9.2 Hz, 1H), 7.72 (dd, *J* = 9.2, 2.5 Hz, 1H), 6.86 (s, 2H), 3.99 (s, 3H), 3.79 (s, 6H), 3.71 (s, 3H). ^13^C NMR (101 MHz, DMSO-*d*_6_) δ 158.8, 153.7, 153.2 (s, 2C), 147.7, 137.7, 134.1, 132.7, 126.0, 123.6, 119.5, 114.2, 104.8 (s, 2C), 103.8, 86.2, 60.3, 56.7, 56.2 (s, 2C). HRMS *m/z* [M+H]^+^ calcd for C_20_H_20_N_3_O_4_: 366.1454, found 366.1461, LC *t*_R_ = 4.40 min, >98% Purity.

**6-(methylsulfonyl)-4-((3,4,5-trimethoxyphenyl)amino)quinoline-3-carbonitrile** (**49**) was obtained as a yellow solid (138 mg, 0.334 mmol, 59 %). MP 231-233 °C; ^1^H NMR (400 MHz, DMSO-*d*_6_) δ 11.60 (s, 1H), 9.45 (d, *J* = 1.9 Hz, 1H), 9.04 (s, 1H), 8.42 (dd, *J* = 8.8, 1.8 Hz, 1H), 8.25 (d, *J* = 8.8 Hz, 1H), 6.83 (s, 2H), 3.78 (s, 3H), 3.70 (s, 6H), 3.42 (s, 3H). ^13^C NMR (101 MHz, DMSO-*d*_6_) δ 154.2, 153.3, 153.1 (s, 2C), 139.2, 137.5, 133.0, 130.6, 125.4, 125.0, 118.4, 114.5, 104.2 (s, 2C), 87.9, 60.3, 56.1 (s, 2C), 43.5. HRMS *m/z* [M+H]^+^ calcd for C_20_H_20_N_3_O_5_S: 414.1124, found 414.1109, LC *t*_R_ = 3.77 min, >98% Purity.

**7-chloro-4-((3,4,5-trimethoxyphenyl)amino)quinoline-3-carbonitrile** (**50**) was obtained as an orange solid (174 mg, 0.489 mmol, 73 %). MP 237-239 °C; ^1^H NMR (400 MHz, DMSO-*d*_6_) δ 11.41 (s, 1H), 9.00 (s, 1H), 8.85 (d, *J* = 9.0 Hz, 1H), 8.16 (d, *J* = 2.2 Hz, 1H), 7.89 (dd, *J* = 9.1, 2.2 Hz, 1H), 6.83 (s, 2H), 3.77 (s, 6H), 3.70 (s, 3H). ^13^C NMR (101 MHz, DMSO-*d*_6_) δ 153.9, 153.1 (s, 2C), 151.7, 141.6, 138.9, 137.6 (s, 2C), 132.8, 128.1, 126.4, 122.1, 117.2, 114.4, 104.5 (s, 2C), 87.0, 60.3, 56.1 (s, 2C). HRMS *m/z* [M+H]^+^ calcd for C_16_H_11_N_3_OBr: 340.0085, found 340.0084, LC *t*_R_ = 3.67 min, >98% Purity.

**7-bromo-4-((3,4,5-trimethoxyphenyl)amino)quinoline-3-carbonitrile** (**51**) was obtained as an orange solid (161 mg, 0.389 mmol, 83 %). MP 257-259 °C; ^1^H NMR (400 MHz, DMSO-*d*_6_) δ 11.57 (s, 1H), 9.04 (s, 1H), 8.80 (d, *J* = 9.1 Hz, 1H), 8.34 (d, *J* = 2.0 Hz, 1H), 8.02 (dd, *J* = 9.1, 2.0 Hz, 1H), 6.84 (s, 2H), 3.78 (s, 6H), 3.70 (s, 3H). ^13^C NMR (101 MHz, DMSO-*d*_6_) δ 154.3, 153.1 (s, 2C), 151.3, 140.8, 137.7 (s, 2C), 132.6, 130.9, 128.1, 126.4, 124.7 117.4, 114.1, 104.6 (s, 2C), 86.9, 60.3, 56.2 (s, 2C). HRMS *m/z* [M+H]^+^ calcd for C_19_H_17_N_3_O_3_Br: 414.0453, found 414.0448, LC *t*_R_ = 4.58 min, >98% Purity.

**7-iodo-4-((3,4,5-trimethoxyphenyl)amino)quinoline-3-carbonitrile** (**52**) was obtained as an orange solid (142 mg, 0.308 mmol, 65 %). MP 262-264 °C; ^1^H NMR (400 MHz, DMSO-*d*_6_) δ 11.29 (s, 1H), 8.97 (s, 1H), 8.51 (d, *J* = 8.9 Hz, 1H), 8.45 (d, *J* = 1.7 Hz, 1H), 8.14 (dd, *J* = 8.9, 1.7 Hz, 1H), 6.82 (s, 2H), 3.77 (s, 6H), 3.70 (s, 3H). ^13^C NMR (101 MHz, DMSO-*d*_6_) δ 154.2, 153.1 (s, 2C), 151.3, 141.3, 137.6, 136.2, 132.8, 131.4, 125.4, 117.7, 114.4, 104.5, 102.7 (s, 2C), 86.8, 60.3, 56.1 (s, 2C). HRMS *m/z* [M+H]^+^ calcd for C_19_H_17_N_3_O_3_I: 462.0315, found 462.0299, LC *t*_R_ = 4.77 min, >98% Purity.

**7-methoxy-4-((3,4,5-trimethoxyphenyl)amino)quinoline-3-carbonitrile** (**53**) was obtained as an orange solid (146 mg, 0.400 mmol, 58 %). MP 235-237 °C; ^1^H NMR (400 MHz, DMSO-*d*_6_) δ 11.53 (s, 1H), 9.03 (s, 1H), 8.82 (d, *J* = 9.4 Hz, 1H), 7.55 (d, *J* = 2.6 Hz, 1H), 7.48 (dd, *J* = 9.4, 2.6 Hz, 1H), 6.85 (s, 2H), 3.98 (s, 3H), 3.74 (d, *J* = 31.2 Hz, 9H). ^13^C NMR (101 MHz, DMSO-*d*_6_) δ 163.9, 154.3, 153.1 (s, 2C), 149.8, 140.7, 137.8 (s, 2C), 132.6, 126.5, 118.8, 114.0, 112.1, 104.9 (s, 2C), 101.6, 85.8, 60.3, 56.3, 56.2 (s, 2C). HRMS *m/z* [M+H]^+^ calcd for C_20_H_20_N_3_O_4_: 366.1454, found 366.1446, LC *t*_R_ = 3.24 min, >98% Purity.

***N*-(3-ethynylphenyl)quinazolin-4-amine** (**54**) was obtained as a yellow solid (204 mg, 0.834 mmol, 91 %). MP 250-252 °C; ^1^H NMR (400 MHz, DMSO-*d*_6_) δ 11.21 (s, 1H), 8.98 – 8.85 (m, 1H), 8.53 (d, *J* = 6.9 Hz, 1H), 8.14 (dd, *J* = 8.6, 1.1 Hz, 1H), 8.02 (ddd, *J* = 8.4, 6.9, 1.2 Hz, 1H), 7.79 (ddd, *J* = 8.4, 7.0, 1.2 Hz, 1H), 7.62 (q, *J* = 1.2 Hz, 1H), 7.58 – 7.57 (m, 1H), 7.56 – 7.49 (m, 1H), 6.83 (d, *J* = 6.9 Hz, 1H), 4.35 (s, 1H). ^13^C NMR (101 MHz, DMSO-*d*_6_) δ 154.8, 142.8, 138.3, 137.7, 133.9, 130.5, 130.4, 128.4, 127.0, 126.2, 124.0, 123.3, 120.2, 117.3, 100.0, 82.5, 82.0. HRMS *m/z* [M+H]^+^ calcd for C_17_H_13_N_2_: 245.1079, found 245.1067, LC *t*_R_ = 3.28 min, >98% Purity.

***N*-(3-ethynylphenyl)-6-methylquinazolin-4-amine** (**55**) was obtained as a colourless solid (130 mg, 0.501 mmol, 60 %). MP 220-222 °C; ^1^H NMR (400 MHz, DMSO-*d*_6_) δ 11.71 (s, 1H), 8.94 (s, 1H), 8.86 (d, *J* = 1.7 Hz, 1H), 8.03 – 7.86 (m, 3H), 7.81 (ddd, *J* = 8.1, 2.2, 1.2 Hz, 1H), 7.51 (t, *J* = 7.9 Hz, 1H), 7.43 (dt, *J* = 7.7, 1.3 Hz, 1H), 4.30 (s, 1H), 2.56 (s, 3H). ^13^C NMR (101 MHz, DMSO-*d*_6_) δ 159.5, 150.3, 138.9, 137.7, 137.1, 136.9, 129.6, 129.2, 127.6, 125.3, 123.9, 122.1, 119.5, 113.5, 82.9, 81.4, 21.2. HRMS *m/z* [M+H]^+^ calcd for C_17_H_14_N_3_: 260.1188, found 260.1179, LC *t*_R_ = 3.45 min, >98% Purity.

***N*-(3-ethynylphenyl)-6-fluoroquinazolin-4-amine** (**56**) was obtained as a colourless solid (164 mg, 0.624 mmol, 76 %). MP 241-243 °C; ^1^H NMR (400 MHz, DMSO-*d*_6_) δ 11.93 (s, 1H), 9.07 (dd, *J* = 9.9, 2.7 Hz, 1H), 8.97 (s, 1H), 8.12 (dd, *J* = 9.2, 5.0 Hz, 1H), 8.04 (ddd, *J* = 9.2, 8.1, 2.6 Hz, 1H), 7.94 (t, *J* = 1.8 Hz, 1H), 7.83 (ddd, *J* = 8.1, 2.2, 1.2 Hz, 1H), 7.54 – 7.47 (m, 1H), 7.42 (dt, *J* = 7.7, 1.3 Hz, 1H), 4.29 (s, 1H). ^13^C NMR (101 MHz, DMSO-*d*_6_) δ 160.4 (d, *J* = 247.7 Hz), 159.6 (d, *J* = 3.7 Hz), 150.8 – 150.7 (m, 1C), 137.0, 136.2, 129.7, 129.2, 127.5, 125.4, 125.2 (d, *J* = 4.2 Hz), 123.1 (d, *J* = 9.0 Hz), 122.1, 115.0 (d, *J* = 9.5 Hz), 110.3 (d, *J* = 25.5 Hz), 82.8, 81.5. HRMS *m/z* [M+H]^+^ calcd for C_16_H_11_N_3_F: 264.0937, found 264.0928, LC *t*_R_ = 3.43 min, >98% Purity.

***N*-(3-ethynylphenyl)-6,7-difluoroquinazolin-4-amine** (**57**) was obtained as a yellow solid (37 mg, 0.132 mmol, 21 %). MP 249-251 °C; ^1^H NMR (400 MHz, DMSO-*d*_6_) δ 11.62 (s, 1H), 9.25 (dd, *J* = 11.6, 8.1 Hz, 1H), 8.94 (s, 1H), 8.01 (dd, *J* = 10.6, 7.4 Hz, 1H), 7.95 (t, *J* = 1.8 Hz, 1H), 7.82 (ddd, *J* = 8.1, 2.2, 1.1 Hz, 1H), 7.50 (t, *J* = 7.9 Hz, 1H), 7.40 (dt, *J* = 7.7, 1.3 Hz, 1H), 4.28 (s, 1H). ^13^C NMR (101 MHz, DMSO-*d*_6_) δ 158.8 (d, *J* = 3.5 Hz), 154.2 (dd, *J* = 258.5, 15.3 Hz), 152.3, 149.0 (dd, *J* = 250.2, 14.4 Hz), 139.5 - 139.2 (m, 1C), 137.2, 129.3, 129.2, 127.1, 124.8, 122.1, 113.1 (d, *J* = 20.7 Hz), 111.2 (dd, *J* = 7.6, 2.2 Hz), 109.6 (d, *J* = 20.0 Hz), 82.9, 81.4. HRMS *m/z* [M+H]^+^ calcd for C_16_H_10_N_3_F_2_: 282.0843, found 282.0835, LC *t*_R_ = 4.62 min, >98% Purity.

**6-chloro-*N*-(3-ethynylphenyl)quinazolin-4-amine** (**58**) was obtained as a colourless solid (158 mg, 0.565 mmol, 75 %). MP 230-232 °C; ^1^H NMR (400 MHz, DMSO-*d*_6_) δ 11.99 (s, 1H), 9.28 (d, *J* = 2.1 Hz, 1H), 8.98 (s, 1H), 8.14 (dd, *J* = 8.9, 2.2 Hz, 1H), 8.06 (d, *J* = 9.0 Hz, 1H), 7.93 (t, *J* = 1.8 Hz, 1H), 7.82 (ddd, *J* = 8.0, 2.2, 1.1 Hz, 1H), 7.55 – 7.46 (m, 1H), 7.42 (dt, *J* = 7.7, 1.3 Hz, 1H), 4.29 (s, 1H). ^13^C NMR (101 MHz, DMSO-*d*_6_) δ 159.1, 151.2, 137.9, 136.9, 136.3, 132.8, 129.7, 129.2, 127.5, 125.2, 124.5, 122.1, 122.1, 114.8, 82.8, 81.5. HRMS *m/z* [M+H]^+^ calcd for C_16_H_11_N_3_Cl: 280.0642, found 280.0632, LC *t*_R_ = 4.12 min, >98% Purity.

**6-bromo-*N*-(3-ethynylphenyl)quinazolin-4-amine** (**59**) was obtained as a colourless solid (168 mg, 0.518 mmol, 84 %). MP 212-214 °C; ^1^H NMR (400 MHz, DMSO-*d*_6_) δ 11.84 (s, 1H), 9.34 (d, *J* = 2.0 Hz, 1H), 8.98 (s, 1H), 8.25 (dd, *J* = 8.9, 2.0 Hz, 1H), 7.97 (d, *J* = 8.9 Hz, 1H), 7.93 (t, *J* = 1.8 Hz, 1H), 7.81 (ddd, *J* = 8.1, 2.2, 1.2 Hz, 1H), 7.57 – 7.47 (m, 1H), 7.43 (dt, *J* = 7.7, 1.3 Hz, 1H), 4.30 (s, 1H). ^13^C NMR (101 MHz, DMSO-*d*_6_) δ 158.8, 151.4, 138.8, 138.5, 137.0, 129.7, 129.3, 127.4, 127.3, 125.1, 122.4, 122.1, 121.0, 115.2, 82.8, 81.5. HRMS *m/z* [M+H]^+^ calcd for C_16_H_11_N_3_Br: 324.0136, found 324.0128, LC *t*_R_ = 4.28 min, >98% Purity.

***N*-(3-ethynylphenyl)-6-iodoquinazolin-4-amine** (**60**) was obtained as a colourless solid (159 mg, 0.428 mmol, 83 %). MP 215-217 °C; ^1^H NMR (400 MHz, DMSO-*d*_6_) δ 11.70 (s, 1H), 9.38 (d, *J* = 1.7 Hz, 1H), 8.98 (s, 1H), 8.38 (dd, *J* = 8.7, 1.7 Hz, 1H), 7.91 (t, *J* = 1.8 Hz, 1H), 7.85 – 7.72 (m, 2H), 7.55 – 7.47 (m, 1H), 7.43 (dt, *J* = 7.7, 1.3 Hz, 1H), 4.30 (s, 1H); ^13^C NMR (101 MHz, DMSO-*d*_6_) δ 158.5, 151.3, 144.2, 138.8, 137.0, 133.0, 129.7, 129.3, 127.4, 125.1, 122.1, 122.0, 115.3, 94.2, 82.8, 81.5. HRMS *m/z* [M+H]^+^ calcd for C_16_H_11_N_3_I: 371.9998, found 371.9978, LC *t*_R_ = 4.45 min, >98% Purity.

***N*-(3-ethynylphenyl)-6-(trifluoromethyl)quinazolin-4-amine** (**61**) was obtained as a colourless solid (47 mg, 0.150 mmol, 23 %). MP 232-234 °C; ^1^H NMR (400 MHz, DMSO-*d*_6_) δ 12.10 (s, 1H), 9.58 – 9.43 (m, 1H), 9.03 (s, 1H), 8.38 (dd, *J* = 8.9, 1.8 Hz, 1H), 8.18 (dd, *J* = 8.8, 0.9 Hz, 1H), 7.91 (t, *J* = 1.8 Hz, 1H), 7.81 (ddd, *J* = 8.1, 2.2, 1.2 Hz, 1H), 7.60 – 7.48 (m, 1H), 7.44 (dt, *J* = 7.7, 1.3 Hz, 1H), 4.30 (s, 1H). ^13^C NMR (101 MHz, DMSO-*d*_6_) δ 159.9, 153.0, 142.1, 136.9, 131.56 (q, *J* = 2.9 Hz), 129.8, 129.3, 128.0 (q, *J* = 33.0 Hz), 127.6, 125.2, 124.9, 123.6 (q, *J* = 4.1 Hz), 122.2 (d, *J* = 3.0 Hz), 122.1, 113.7, 82.8, 81.5. HRMS *m/z* [M+H]^+^ calcd for C_17_H_11_N_3_F_3_: 314.0905, found 314.0890, LC *t*_R_ = 5.14 min, >98% Purity.

***N*-(3-ethynylphenyl)-7-fluoroquinazolin-4-amine** (**62**) was obtained as a colourless solid (181 mg, 0.688 mmol, 84 %). MP 209-211 °C; ^1^H NMR (400 MHz, DMSO-*d*_6_) δ 11.67 (s, 1H), 9.04 (dd, *J* = 9.3, 5.5 Hz, 1H), 8.95 (s, 1H), 7.91 (t, *J* = 1.8 Hz, 1H), 7.85 – 7.66 (m, 3H), 7.55 – 7.47 (m, 1H), 7.42 (dt, *J* = 7.7, 1.3 Hz, 1H), 4.29 (s, 1H). ^13^C NMR (101 MHz, DMSO-*d*_6_) δ 165.6 (d, *J* = 255.7 Hz), 159.3, 152.4, 142.3, 137.1, 129.5, 129.3, 128.6 (d, *J* = 10.9 Hz), 127.5, 125.1, 122.1, 117.6 (d, *J* = 24.0 Hz), 111.0 (d, *J* = 1.5 Hz), 106.1 (d, *J* = 24.6 Hz), 82.9, 81.4. HRMS *m/z* [M+H]^+^ calcd for C_16_H_11_N_3_F: 264.0937, found 264.0928, LC *t*_R_ = 3.38 min, >98% Purity.

**7-chloro-*N*-(3-ethynylphenyl)quinazolin-4-amine** (**63**) was obtained as a yellow solid (92 mg, 0.329 mmol, 44 %). MP 218-220 °C; ^1^H NMR (400 MHz, DMSO-*d*_6_) δ 11.95 (s, 1H), 9.06 (d, *J* = 9.0 Hz, 1H), 8.97 (s, 1H), 8.07 (d, *J* = 2.1 Hz, 1H), 8.01 – 7.86 (m, 2H), 7.80 (ddd, *J* = 8.1, 2.2, 1.2 Hz, 1H), 7.53 – 7.47 (m, 1H), 7.43 (dt, *J* = 7.7, 1.3 Hz, 1H), 4.29 (s, 1H). ^13^C NMR (101 MHz, DMSO-*d*_6_) δ 159.5, 152.0, 140.5, 140.4, 137.0, 129.7, 129.2, 128.8, 127.6, 127.3, 125.3, 122.1, 119.4, 112.6, 82.8, 81.5. HRMS *m/z* [M+H]^+^ calcd for C_16_H_11_N_3_Cl: 280.0642, found 280.0633, LC *t*_R_ = 4.00 min, >98% Purity.

**7-bromo-*N*-(3-ethynylphenyl)quinazolin-4-amine** (**64**) was obtained as a colourless solid (131 mg, 0.404 mmol, 66 %). MP 225-227 °C; ^1^H NMR (400 MHz, DMSO-*d*_6_) δ 11.62 (s, 1H), 8.94 (s, 1H), 8.84 (d, *J* = 8.9 Hz, 1H), 8.17 (d, *J* = 1.9 Hz, 1H), 8.06 (dd, *J* = 8.9, 2.0 Hz, 1H), 7.93 (t, *J* = 1.8 Hz, 1H), 7.80 (ddd, *J* = 8.1, 2.2, 1.2 Hz, 1H), 7.51 (t, *J* = 7.9 Hz, 1H), 7.42 (dt, *J* = 7.7, 1.3 Hz, 1H), 4.29 (s, 1H). ^13^C NMR (101 MHz, DMSO-*d*_6_) δ 159.5, 152.3, 141.4, 137.2, 131.4, 129.5, 129.4, 129.3, 127.3, 126.8, 125.0, 123.3, 122.1, 112.9, 82.9, 81.4. HRMS *m/z* [M+H]^+^ calcd for C_16_H_11_N_3_Br: 324.0136, found 324.0128, LC *t*_R_ = 4.19 min, >98% Purity.

***N*-(3-ethynylphenyl)-7-iodoquinazolin-4-amine** (**65**) was obtained as a colourless solid (134 mg, 0.361 mmol, 70 %). MP 230-232 °C; ^1^H NMR (400 MHz, DMSO-*d*_6_) δ 11.86 (s, 1H), 8.94 (s, 1H), 8.74 (d, *J* = 8.8 Hz, 1H), 8.39 (d, *J* = 1.6 Hz, 1H), 8.19 (dd, *J* = 8.8, 1.7 Hz, 1H), 7.91 (t, *J* = 1.8 Hz, 1H), 7.79 (ddd, *J* = 8.0, 2.2, 1.1 Hz, 1H), 7.56 – 7.46 (m, 1H), 7.42 (dt, *J* = 7.7, 1.3 Hz, 1H), 4.29 (s, 1H). ^13^C NMR (101 MHz, DMSO-*d*_6_) δ 159.8, 151.5, 139.9, 137.1, 137.0, 129.7, 129.2, 128.4, 127.5, 126.2, 125.2, 122.1, 113.0, 105.0, 82.8, 81.5. HRMS *m/z* [M+H]^+^ calcd for C_16_H_11_N_3_I: 371.9998, found 371.9983, LC *t*_R_ = 4.43 min, >98% Purity.

***N*-(3-ethynylphenyl)-7-(trifluoromethyl)quinazolin-4-amine** (**66**) was obtained as a colourless solid (36 mg, 0.115 mmol, 21 %). MP 173-175 °C; ^1^H NMR (400 MHz, DMSO-*d*_6_) δ 10.11 (s, 1H), 9.00 – 8.61 (m, 2H), 8.23 – 7.76 (m, 4H), 7.43 (t, *J* = 7.9 Hz, 1H), 7.27 (dt, *J* = 7.6, 1.3 Hz, 1H), 4.22 (s, 1H). ^13^C NMR (101 MHz, DMSO-*d*_6_) δ 157.6, 155.8, 149.3, 139.0, 132.9 (q, *J* = 32.2 Hz), 129.0, 127.3, 125.2 (d, *J* = 4.0 Hz), 125.2 (d, *J* = 4.3 Hz), 125.0 (d, *J* = 12.7 Hz), 123.0, 122.2, 121.8, 121.8 (d, *J* = 3.4 Hz), 117.3, 83.4, 80.8. HRMS *m/z* [M+H]^+^ calcd for C_17_H_11_N_3_F_3_: 314.0905, found 314.0891, LC *t*_R_ = 5.26 min, >98% Purity.

**4-((3-ethynylphenyl)amino)quinazoline-7-carbonitrile** (**67**) was obtained as a colourless solid (45 mg, 0.167 mmol, 21 %). MP 208-210 °C; ^1^H NMR (400 MHz, DMSO-*d*_6_) δ 10.12 (s, 1H), 8.91 – 8.55 (m, 2H), 8.33 (d, *J* = 1.6 Hz, 1H), 8.05 (t, *J* = 1.8 Hz, 1H), 8.02 (dd, *J* = 8.6, 1.7 Hz, 1H), 7.95 – 7.82 (m, 1H), 7.43 (t, *J* = 7.9 Hz, 1H), 7.27 (dt, *J* = 7.7, 1.3 Hz, 1H), 4.22 (s, 1H). ^13^C NMR (101 MHz, DMSO-*d*_6_) δ 157.6, 155.8, 148.9, 138.9, 133.0, 129.0, 127.7, 127.4, 125.3, 124.9, 123.1, 121.9, 117.9, 117.6, 115.5, 83.3, 80.8. HRMS *m/z* [M+H]^+^ calcd for C_17_H_11_N_4_: 271.0978, found 271.0978, LC *t*_R_ = 5.07 min, >98% Purity.

**4-((3-ethynylphenyl)amino)quinazoline-6-carbonitrile** (**68**) was obtained as a colourless solid (43 mg, 0.158 mmol, 20 %). MP 230-232 °C; ^1^H NMR (400 MHz, DMSO-*d*_6_) δ 11.88 (s, 1H), 9.60 (d, *J* = 1.6 Hz, 1H), 9.00 (s, 1H), 8.39 (dd, *J* = 8.7, 1.6 Hz, 1H), 8.08 (d, *J* = 8.7 Hz, 1H), 7.96 (t, *J* = 1.8 Hz, 1H), 7.84 (ddd, *J* = 8.1, 2.2, 1.1 Hz, 1H), 7.56 – 7.47 (m, 1H), 7.43 (dt, *J* = 7.7, 1.3 Hz, 1H), 4.30 (s, 1H). ^13^C NMR (101 MHz, DMSO-*d*_6_) δ 159.0, 153.7, 143.2, 137.0, 136.9, 131.2, 129.6, 129.3, 127.1, 124.8, 122.7, 122.1, 117.7, 114.1, 110.1, 82.8, 81.5. HRMS *m/z* [M+H]^+^ calcd for C_17_H_11_N_4_: 271.0984, found 271.0975, LC *t*_R_ = 4.27 min, >98% Purity.

***N*-(3-ethynylphenyl)-6-(methylsulfonyl)quinazolin-4-amine** (**69**) was obtained as a colourless solid (169 mg, 0.523 mmol, 85 %). MP 282-284 °C; ^1^H NMR (400 MHz, DMSO-*d*_6_) δ 12.31 (s, 1H), 9.71 (d, *J* = 1.8 Hz, 1H), 9.03 (s, 1H), 8.51 (dd, *J* = 8.8, 1.9 Hz, 1H), 8.20 (d, *J* = 8.8 Hz, 1H), 7.93 (t, *J* = 1.8 Hz, 1H), 7.83 (ddd, *J* = 8.1, 2.2, 1.2 Hz, 1H), 7.57 – 7.48 (m, 1H), 7.44 (dt, *J* = 7.7, 1.3 Hz, 1H), 4.30 (s, 1H), 3.47 (s, 3H). ^13^C NMR (101 MHz, DMSO-*d*_6_) δ 160.5, 153.6, 142.8, 140.4, 137.3, 133.1, 130.3, 129.7, 128.0, 126.1, 125.6, 122.6, 122.3, 114.2, 83.2, 82.0. HRMS *m/z* [M+H]^+^ calcd for C_17_H_14_N_3_O_2_S: 324.0807, found 324.0797, LC *t*_R_ = 3.85 min, >98% Purity.

***N*-(3-ethynylphenyl)-**[**1,3**]**dioxolo**[**4,5-*g***]**quinazolin-8-amine** (**73**) (25 mg scale) was obtained as a colourless solid (24 mg, 0.083 mmol, 70 %). MP >300 °C; ^1^H NMR (400 MHz, DMSO-*d*_6_) δ 11.27 (s, 1H), 8.84 (s, 1H), 8.46 (s, 1H), 7.89 (t, *J* = 1.8 Hz, 1H), 7.77 (ddd, *J* = 8.1, 2.2, 1.1 Hz, 1H), 7.51 – 7.45 (m, 1H), 7.44 (s, 1H), 7.39 (dt, *J* = 7.7, 1.3 Hz, 1H), 6.38 (s, 2H), 4.27 (s, 1H). ^13^C NMR (101 MHz, DMSO-*d*_6_) δ 158.5, 154.6, 149.0, 148.9, 137.7, 137.4, 129.2, 129.1, 127.4, 125.1, 122.0, 108.9, 103.9, 100.9, 97.7, 82.9, 81.3. HRMS *m/z* [M+H]^+^ calcd for C_17_H_12_N_3_O_2_: 290.0930, found 290.0920, LC *t*_R_ = 3.36 min, >98% Purity.

***N*-(3-ethynylphenyl)-7,8-dihydro-**[**1,4**]**dioxino**[**2,3-*g***]**quinazolin-4-amine** (**74**) (25 mg scale) was obtained as a yellow solid (24 mg, 0.079 mmol, 70 %). MP >300 °C; ^1^H NMR (400 MHz, DMSO-*d*_6_) δ 11.42 (s, 1H), 8.84 (s, 1H), 8.55 (s, 1H), 7.90 (t, *J* = 1.8 Hz, 1H), 7.79 (ddd, *J* = 8.1, 2.2, 1.1 Hz, 1H), 7.52 – 7.45 (m, 1H), 7.44 (s, 1H), 7.40 (dt, *J* = 7.7, 1.3 Hz, 1H), 4.48 (ddd, *J* = 23.4, 6.0, 2.7 Hz, 4H), 4.28 (s, 1H). ^13^C NMR (101 MHz, DMSO-*d*_6_) δ 158.4, 151.4, 149.3, 145.0, 137.2, 134.6, 129.4, 129.2, 127.5, 125.2, 122.0, 111.0, 108.1, 105.3, 82.9, 81.4, 65.1, 64.1. HRMS *m/z* [M+H]^+^ calcd for C_18_H_14_N_3_O_2_: 304.1086, found 304.1076, LC *t*_R_ = 3.50 min, >98% Purity.

**4-((3-ethynylphenyl)amino)quinoline-3-carbonitrile** (**75**) was obtained as a yellow solid (197 mg, 0.732 mmol, 92 %). MP 236-238 °C; ^1^H NMR (400 MHz, DMSO-*d*_6_) δ 11.84 (s, 1H), 9.13 (s, 1H), 8.98 (dd, *J* = 8.6, 1.2 Hz, 1H), 8.17 (dd, *J* = 8.5, 1.2 Hz, 1H), 8.09 (ddd, *J* = 8.3, 7.0, 1.1 Hz, 1H), 7.86 (ddd, *J* = 8.4, 7.0, 1.3 Hz, 1H), 7.78 – 7.25 (m, 4H), 4.32 (s, 1H). ^13^C NMR (101 MHz, DMSO-*d*_6_) δ 154.8, 149.9, 138.8, 137.5, 135.1, 131.5, 129.7, 129.4, 128.3, 127.2, 124.7, 122.6, 121.6, 118.5, 113.9, 86.6, 82.7, 81.9. HRMS *m/z* [M+H]^+^ calcd for C_18_H_12_N_3_: 270.1031, found 270.1018, LC *t*_R_ = 3.45 min, >98% Purity.

**4-((3-ethynylphenyl)amino)-6-fluoroquinoline-3-carbonitrile** (**76**) was obtained as a yellow solid (121 mg, 0.421 mmol, 58 %). MP 251-253 °C; ^1^H NMR (400 MHz, DMSO-*d*_6_) δ 11.35 (s, 1H), 9.04 (s, 1H), 8.77 (dd, *J* = 10.6, 2.7 Hz, 1H), 8.19 (dd, *J* = 9.2, 5.2 Hz, 1H), 7.99 (ddd, *J* = 9.3, 7.9, 2.7 Hz, 1H), 7.75 – 7.26 (m, 4H), 4.30 (s, 1H). ^13^C NMR (101 MHz, DMSO-*d*_6_) δ 160.4 (d, *J* = 247.1 Hz), 153.5, 150.2, 138.0, 137.8, 131.0, 129.7, 128.8, 126.6, 126.4 – 126.2 (m, 1C), 123.7 (d, *J* = 25.5 Hz), 122.6, 120.1 (d, *J* = 9.9 Hz), 114.4, 109.3 (d, *J* = 25.9 Hz), 87.1, 82.7, 81.8. HRMS *m/z* [M+H]^+^ calcd for C_18_H_11_N_3_F: 288.0937, found 288.0923, LC *t*_R_ = 4.55 min, >98% Purity.

**6-chloro-4-((3-ethynylphenyl)amino)quinoline-3-carbonitrile** (**77**) was obtained as a yellow solid (119 mg, 0.392 mmol, 70 %). MP 307-309 °C; ^1^H NMR (400 MHz, DMSO-*d*_6_) δ 11.55 (s, 1H), 9.06 (d, *J* = 2.1 Hz, 1H), 9.05 (s, 1H), 8.15 (d, *J* = 9.0 Hz, 1H), 8.08 (dd, *J* = 9.0, 2.1 Hz, 1H), 7.56 (q, *J* = 1.2 Hz, 1H), 7.53 – 7.46 (m, 3H), 4.30 (s, 1H). ^13^C NMR (101 MHz, DMSO-*d*_6_) δ 153.3, 150.8, 139.4, 137.7, 134.6, 132.6, 131.1, 129.7, 128.8, 126.6, 125.2, 123.8, 122.6, 119.9, 114.2, 87.5, 82.7, 81.8. HRMS *m/z* [M+H]^+^ calcd for C_18_H_11_N_3_Cl: 304.0642, found 304.0633, LC *t*_R_ = 5.17 min, >98% Purity.

**6-bromo-4-((3-ethynylphenyl)amino)quinoline-3-carbonitrile** (**78**) was obtained as a yellow solid (126 mg, 0.362 mmol, 65 %). MP 240-242 °C; ^1^H NMR (400 MHz, DMSO-*d*_6_) δ 8.94 (d, *J* = 2.1 Hz, 1H), 8.86 (s, 1H), 8.09 (dd, *J* = 8.9, 1.9 Hz, 1H), 7.94 (d, *J* = 8.8 Hz, 1H), 7.60 – 7.24 (m, 4H), 4.25 (s, 1H). ^13^C NMR (100 MHz, DMSO-*d*_6_) δ 152.1, 151.6, 138.7, 136.2, 130.1, 129.6, 128.3, 127.8, 126.1, 125.6, 122.5, 121.9, 120.6, 120.4, 115.3, 88.3, 82. 9, 81.5. HRMS *m/z* [M+H]^+^ calcd for C_18_H_11_N_3_Br: 348.0136, found 348.0134, LC *t*_R_ = 5.32 min, >98% Purity.

**4-((3-ethynylphenyl)amino)-6-iodoquinoline-3-carbonitrile** (**79**) was obtained as a yellow solid (113 mg, 0.286 mmol, 60 %). MP 249-251 °C; ^1^H NMR (400 MHz, DMSO-*d*_6_) δ 11.30 (s, 1H), 9.22 (d, *J* = 1.8 Hz, 1H), 9.03 (s, 1H), 8.31 (dd, *J* = 8.8, 1.7 Hz, 1H), 7.85 (d, *J* = 8.8 Hz, 1H), 7.70 - 7.30 (m, 4H), 4.30 (s, 1H). ^13^C NMR (101 MHz, DMSO-*d*_6_) δ 152.7, 150.8, 142.5, 140.2, 137.8, 132.5, 131.0, 129.7, 128.7, 126.5, 125.0, 122.6, 120.4, 114.4, 94.2, 87.5, 82.7, 81.8. HRMS *m/z* [M+H]^+^ calcd for C_18_H_11_N_3_I: 395.9998, found 395.9982, LC *t*_R_ = 5.54 min, >98% Purity.

**4-((3-ethynylphenyl)amino)-6-(methylsulfonyl)quinoline-3-carbonitrile** (**80**) was obtained as an orange solid (130 mg, 0.374 mmol, 67 %). MP 247-249 °C; ^1^H NMR (400 MHz, DMSO-*d*_6_) δ 11.83 (s, 1H), 9.49 (d, *J* = 1.9 Hz, 1H), 9.10 (s, 1H), 8.43 (dd, *J* = 8.8, 1.8 Hz, 1H), 8.28 (d, *J* = 8.8 Hz, 1H), 7.58 (q, *J* = 1.3 Hz, 1H), 7.51 (t, *J* = 1.5 Hz, 3H), 4.30 (s, 1H), 3.43 (s, 3H). ^13^C NMR (101 MHz, DMSO-*d*_6_) δ 154.2, 153.1, 143.8, 139.4, 137.9, 131.0, 130.7, 129.7 (s, 2C), 128.6, 126.4, 125.2, 125.1, 122.6, 118.8, 114.3, 88.2, 82.7, 81.8, 43.5. HRMS *m/z* [M+H]^+^ calcd for C_19_H_14_N_3_O_2_S: 348.0807, found 348.0795, LC *t*_R_ = 3.18 min, >98% Purity.

**7-chloro-4-((3-ethynylphenyl)amino)quinoline-3-carbonitrile** (**84**) was obtained as a yellow solid (94 mg, 0.310 mmol, 46 %). MP 254-256 °C; ^1^H NMR (400 MHz, DMSO-*d*_6_) δ 11.61 (s, 1H), 9.06 (s, 1H), 8.90 (d, *J* = 9.1 Hz, 1H), 8.18 (d, *J* = 2.2 Hz, 1H), 7.90 (dd, *J* = 9.1, 2.2 Hz, 1H), 7.57 (dt, *J* = 2.4, 1.2 Hz, 1H), 7.50 (d, *J* = 1.4 Hz, 3H), 4.30 (s, 1H). ^13^C NMR (101 MHz, DMSO-*d*_6_) δ 154.3, 152.0, 141.9, 139.5, 138.2, 131.6, 130.1 (s, 2C), 129.3, 128.7, 127.15, 127.13, 123.0, 122.4, 118.11, 114.6, 87.9, 83.1, 82.2. HRMS *m/z* [M+H]^+^ calcd for C_18_H_10_ClN_3_: 303.0563, found 303.0567, LC *t*_R_ = 4.36 min, >98% Purity.

**7-bromo-4-((3-ethynylphenyl)amino)quinoline-3-carbonitrile** (**85**) was obtained as a yellow solid (128 mg, 0.368 mmol, 66 %). MP 253-255 °C; ^1^H NMR (400 MHz, DMSO-*d*_6_) δ 11.34 (s, 1H), 9.01 (s, 1H), 8.73 (d, *J* = 9.1 Hz, 1H), 8.29 (d, *J* = 2.0 Hz, 1H), 8.00 (dd, *J* = 9.1, 2.1 Hz, 1H), 7.55 (q, *J* = 1.3 Hz, 1H), 7.49 (q, *J* = 1.9 Hz, 3H), 4.30 (s, 1H). ^13^C NMR (101 MHz, DMSO-*d*_6_) δ 153.6, 151.8, 142.5, 138.0, 130.9, 130.8, 129.7, 128.6, 127.8, 126.43 (s, 2C), 126.3, 125.9, 122.6, 118.0, 114.6, 87.6, 82.7, 81.7. HRMS *m/z* [M+H]^+^ calcd for C_18_H_11_N_3_Br: 348.0136, found 348.0130, LC *t*_R_ = 5.48 min, >98% Purity.

**4-((3-ethynylphenyl)amino)-7-iodoquinoline-3-carbonitrile** (**86**) was obtained as a yellow solid (138 mg, 0.349 mmol, 73 %). MP 247-249 °C; ^1^H NMR (400 MHz, DMSO-*d*_6_) δ 11.43 (s, 1H), 9.00 (s, 1H), 8.57 (d, *J* = 8.9 Hz, 1H), 8.50 (d, *J* = 1.6 Hz, 1H), 8.13 (dd, *J* = 8.9, 1.7 Hz, 1H), 7.55 (q, *J* = 1.3 Hz, 1H), 7.50 – 7.44 (m, 3H), 4.30 (s, 1H). ^13^C NMR (101 MHz, DMSO-*d*_6_) δ 154.0, 151.1, 141.6, 137.9, 136.3, 131.6, 131.0, 129.6, 128.7, 126.5, 125.7, 122.6, 118.1, 114.4, 102.9, 87.4, 82.7, 81.8. HRMS *m/z* [M+H]^+^ calcd for C_18_H_11_N_3_I: 395.9998, found 395.9984, LC *t*_R_ = 5.60 min, >98% Purity.

**4-((5-ethynyl-2-fluorophenyl)amino)quinazoline-7-carbonitrile** (**87**) was obtained as a colourless solid (201 mg, 0.697 mmol, 88 %). MP 224-226 °C; ^1^H NMR (400 MHz, DMSO-*d*_6_) δ 12.01 (s, 1H), 9.09 (d, *J* = 8.6 Hz, 1H), 8.93 (s, 1H), 8.46 (d, *J* = 1.5 Hz, 1H), 8.22 (dd, *J* = 8.6, 1.6 Hz, 1H), 7.70 (dd, *J* = 7.2, 2.2 Hz, 1H), 7.57 (ddd, *J* = 8.6, 4.7, 2.2 Hz, 1H), 7.45 (dd, *J* = 10.1, 8.6 Hz, 1H), 4.29 (s, 1H). ^13^C NMR (101 MHz, DMSO-*d*_6_) δ 160.0, 158.0, 155.6, 153.3, 141.1, 132.7 (d, *J* = 8.1 Hz, 1C), 131.7, 129.8, 126.8, 126.5, 124.9 (d, *J* = 13.2 Hz, 1C), 118.4 (d, *J* = 3.8 Hz), 117.3 (t, *J* = 6.9 Hz, 1C), 117.0, 116.2, 81.6, 81.4. HRMS *m/z* [M+H]^+^ calcd for C_17_H_10_N_4_F: 289.0889, found 289.0880, LC *t*_R_ = 4.62 min, >98% Purity.

**4-((3-ethynyl-2-fluorophenyl)amino)quinazoline-7-carbonitrile** (**88**) was obtained as a yellow solid (144 mg, 0.500 mmol, 63 %). MP 234-236 °C; ^1^H NMR (400 MHz, DMSO-*d*_6_) δ 11.35 (s, 1H), 8.88 (d, *J* = 8.6 Hz, 1H), 8.79 (s, 1H), 8.39 (d, *J* = 1.6 Hz, 1H), 8.13 (dd, *J* = 8.6, 1.6 Hz, 1H), 7.63 – 7.53 (m, 2H), 7.32 (td, *J* = 7.9, 0.8 Hz, 1H), 4.59 (s, 1H). ^13^C NMR (101 MHz, DMSO-*d*_6_) δ 159.2, 157.3 (d, *J* = 254.1 Hz), 154.3, 147.3, 144.2, 132.1, 131.9, 129.3, 129.0, 127.3, 125.9, 124.77 (d, *J* = 4.5 Hz), 117.5, 116.7, 111.0 (d, *J* = 14.5 Hz), 86.9 (d, *J* = 3.5 Hz), 76.4. HRMS *m/z* [M+H]^+^ calcd for C_17_H_10_N_4_F: 289.0889, found 289.0879, LC *t*_R_ = 4.46 min, >98% Purity.

**4-((3-ethynyl-4-fluorophenyl)amino)quinazoline-7-carbonitrile** (**89**) was obtained as a yellow solid (190 mg, 0.659 mmol, 83 %). MP 240-242 °C; ^1^H NMR (400 MHz, DMSO-*d*_6_) δ 11.90 (s, 1H), 9.19 – 9.11 (m, 1H), 8.96 (s, 1H), 8.49 – 8.39 (m, 1H), 8.20 (dd, *J* = 8.6, 1.6 Hz, 1H), 8.01 (dd, *J* = 6.4, 2.7 Hz, 1H), 7.86 (ddd, *J* = 9.0, 4.7, 2.7 Hz, 1H), 7.44 (t, *J* = 9.1 Hz, 1H), 4.60 (d, *J* = 0.6 Hz, 1H). ^13^C NMR (101 MHz, DMSO-*d*_6_) δ 160.20 (d, *J* = 249.9 Hz), 159.2, 153.2, 141.0, 133.3 (d, *J* = 3.3 Hz, 1C), 129.6, 129.3, 127.2 (d, *J* = 8.3 Hz, 1C), 126.6, 126.4, 117.2 (d, *J* = 15.0 Hz, 1C), 116.6, 116.1 (d, *J* = 22.2 Hz, 1C), 110.1, 110.0, 86.8 (d, *J* = 3.3 Hz), 76.3. HRMS *m/z* [M+H]^+^ calcd for C_17_H_10_N_4_F: 289.0889, found 289.0880, LC *t*_R_ = 4.58 min, >98% Purity.

**4-((4-chloro-3-ethynylphenyl)amino)quinazoline-7-carbonitrile** (**90**) was obtained as a yellow solid (241 mg, 0.791 mmol, 75 %). MP 240-242 °C; ^1^H NMR (400 MHz, DMSO-*d*_6_) δ 11.66 (s, 1H), 9.10 (d, *J* = 8.6 Hz, 1H), 8.96 (s, 1H), 8.41 (d, *J* = 1.6 Hz, 1H), 8.18 (dd, *J* = 8.7, 1.6 Hz, 1H), 8.13 (d, *J* = 2.6 Hz, 1H), 7.90 (dd, *J* = 8.8, 2.6 Hz, 1H), 7.65 (d, *J* = 8.8 Hz, 1H), 4.66 (s, 1H). ^13^C NMR (101 MHz, DMSO-*d*_6_) δ 158.9, 153.5, 142.2, 136.3, 131.6, 129.6, 129.4, 128.6, 127.6, 126.2, 125.9, 121.5, 117.3, 117.0, 116.8, 86.6, 79.7. HRMS *m/z* [M+H]^+^ calcd for C_17_H_10_N_4_Cl: 305.0594, found 305.0584, LC *t*_R_ = 5.31 min, >98% Purity.

***N*-(5-ethynyl-2-fluorophenyl)-7-iodoquinazolin-4-amine** (**91**) was obtained as a colourless solid (169 mg, 0.434 mmol, 84 %). MP 295-297 °C; ^1^H NMR (400 MHz, DMSO-*d*_6_) δ 11.87 (s, 1H), 8.88 (s, 1H), 8.58 (d, *J* = 8.8 Hz, 1H), 8.38 (d, *J* = 1.6 Hz, 1H), 8.22 (dd, *J* = 8.7, 1.6 Hz, 1H), 7.70 (dd, *J* = 7.2, 2.1 Hz, 1H), 7.57 (ddd, *J* = 8.6, 4.7, 2.2 Hz, 1H), 7.45 (dd, *J* = 10.0, 8.6 Hz, 1H), 4.29 (s, 1H). ^13^C NMR (101 MHz, DMSO-*d*_6_) δ 160.1, 156.8 (d, *J* = 252.5 Hz), 152.7, 143.0, 136.7, 132.3 (d, *J* = 8.2 Hz), 131.7, 130.8, 125.7, 125.3 (d, *J* = 13.3 Hz), 118.3 (d, *J* = 3.8 Hz), 117.0 (d, *J* = 21.1 Hz), 113.0, 104.1, 81.8, 81.2. HRMS *m/z* [M+H]^+^ calcd for C_16_H_10_N_3_FI: 389.9904, found 389.9892, LC *t*_R_ = 3.72 min, >98% Purity.

***N*-(3-ethynyl-2-fluorophenyl)-7-iodoquinazolin-4-amine** (**92**) was obtained as a grey solid (121 mg, 0.311 mmol, 60 %). MP 232-234 °C; ^1^H NMR (400 MHz, DMSO-*d*_6_) δ 12.02 (s, 1H), 8.90 (s, 1H), 8.63 (d, *J* = 8.8 Hz, 1H), 8.40 (d, *J* = 1.6 Hz, 1H), 8.22 (dd, *J* = 8.7, 1.7 Hz, 1H), 7.69 – 7.53 (m, 2H), 7.35 (td, *J* = 7.9, 0.8 Hz, 1H), 4.61 (s, 1H). ^13^C NMR (101 MHz, DMSO-*d*_6_) δ 160.7, 157.3 (d, *J* = 254.7 Hz), 151.8, 140.4, 137.3, 132.8, 129.5, 128.9, 126.1, 125.0 – 124.6 (m, 2C), 112.6, 111.1 (d, *J* = 14.5 Hz, 1C), 105.2, 87.2 (d, *J* = 3.5 Hz, 1C), 76.2. HRMS *m/z* [M+H]^+^ calcd for C_16_H_10_N_3_FI: 389.9904, found 389.9889, LC *t*_R_ = 4.62 min, >98% Purity.

***N*-(3-ethynyl-4-fluorophenyl)-7-iodoquinazolin-4-amine** (**93**) was obtained as a yellow solid (195 mg, 0.501 mmol, 97 %). MP 277-279 °C; ^1^H NMR (400 MHz, DMSO-*d*_6_) δ 11.77 (s, 1H), 8.92 (s, 1H), 8.68 (d, *J* = 8.8 Hz, 1H), 8.36 (d, *J* = 1.6 Hz, 1H), 8.19 (dd, *J* = 8.7, 1.7 Hz, 1H), 7.97 (dd, *J* = 6.4, 2.7 Hz, 1H), 7.82 (ddd, *J* = 8.9, 4.7, 2.7 Hz, 1H), 7.44 (t, *J* = 9.1 Hz, 1H), 4.60 (s, 1H). ^13^C NMR (101 MHz, DMSO-*d*_6_) δ 160.3 (d, *J* = 249.9 Hz), 159.7, 151.8, 140.5, 137.0, 133.2 (d, *J* = 3.3 Hz), 129.5, 128.9, 127.4 (d, *J* = 8.3 Hz), 126.0, 116.1 (d, *J* = 22.3 Hz), 113.0, 110.1 (d, *J* = 16.9 Hz), 104.7, 86.9 (d, *J* = 3.2 Hz), 76.3. HRMS *m/z* [M+H]^+^ calcd for C_16_H_10_N_3_FI: 389.9904, found 389.9892, LC *t*_R_ = 4.59 min, >98% Purity.

***N*-(4-chloro-3-ethynylphenyl)-7-iodoquinazolin-4-amine** (**94**) was obtained as a yellow solid (142 mg, 0.350 mmol, 68 %). MP 271-273 °C; ^1^H NMR (400 MHz, DMSO-*d*_6_) δ 11.79 (s, 1H), 8.96 (s, 1H), 8.70 (d, *J* = 8.8 Hz, 1H), 8.38 (d, *J* = 1.6 Hz, 1H), 8.20 (dd, *J* = 8.8, 1.7 Hz, 1H), 8.08 (d, *J* = 2.6 Hz, 1H), 7.85 (dd, *J* = 8.8, 2.6 Hz, 1H), 7.66 (d, *J* = 8.8 Hz, 1H), 4.67 (s, 1H). ^13^C NMR (101 MHz, DMSO-*d*_6_) δ 159.7, 151.7, 140.5, 137.0, 135.9, 132.1, 129.7, 129.1, 128.9, 126.3, 126.1, 121.5, 113.1, 104.9, 86.7, 79.6. HRMS *m/z* [M+H]^+^ calcd for C_16_H_10_N_3_ClI: 405.9608, found 405.9897, LC *t*_R_ = 5.31 min, >98% Purity.

***N*-(5-ethynyl-2-fluorophenyl)-6-iodoquinazolin-4-amine** (**95**) was obtained as a colourless solid (112 mg, 0.288 mmol, 56 %). MP 231-233 °C; ^1^H NMR (400 MHz, DMSO-*d*_6_) δ 11.99 (s, 1H), 9.35 (d, *J* = 1.8 Hz, 1H), 8.94 (s, 1H), 8.40 (dd, *J* = 8.8, 1.7 Hz, 1H), 7.80 (d, *J* = 8.8 Hz, 1H), 7.70 (dd, *J* = 7.1, 2.1 Hz, 1H), 7.58 (ddd, *J* = 8.6, 4.8, 2.2 Hz, 1H), 7.46 (dd, *J* = 10.0, 8.6 Hz, 1H), 4.30 (s, 1H). ^13^C NMR (101 MHz, DMSO-*d*_6_) δ 159.9, 157.3 (d, *J* = 252.9 Hz, 1C), 151.9, 144.9, 139.4, 133.6, 133.28 (d, *J* = 8.2 Hz), 132.26, 125.1 (d, *J* = 13.6 Hz), 122.6, 118.4 (d, *J* = 3.8 Hz), 117.6 (d, *J* = 21.0 Hz)., 115.4, 94.8, 82.2, 81.8. HRMS *m/z* [M+H]^+^ calcd for C_16_H_10_N_3_FI: 389.9904, found 389.9892, LC *t*_R_ = 4.76 min, >98% Purity.

***N*-(3-ethynyl-2-fluorophenyl)-6-iodoquinazolin-4-amine** (**96**) was obtained as a colourless solid (145 mg, 0.373 mmol, 72 %). MP 245-247 °C; ^1^H NMR (400 MHz, DMSO-*d*_6_) δ 11.94 (s, 1H), 9.36 – 9.29 (m, 1H), 8.93 (t, *J* = 1.0 Hz, 1H), 8.40 (dd, *J* = 8.8, 1.7 Hz, 1H), 7.79 (dt, *J* = 8.8, 1.9 Hz, 1H), 7.61 (dd, *J* = 7.9, 6.8 Hz, 2H), 7.39 – 7.32 (m, 1H), 4.62 (s, 1H). ^13^C NMR (101 MHz, DMSO-*d*_6_) δ 159.3, 157.26 (d, *J* = 254.7 Hz), 151.6, 144.4, 139.1, 133.1 (d, *J* = 2.8 Hz), 132.7, 129.4, 124.9 (d, *J* = 4.6 Hz), 124.7, 122.3, 114.9, 111.1 (d, *J* = 14.5 Hz), 94.4 (d, *J* = 3.2 Hz), 87.2 (d, *J* = 3.4 Hz), 76.2. HRMS *m/z* [M+H]^+^ calcd for C_16_H_10_N_3_FI: 389.9904, found 389.9889, LC *t*_R_ = 3.89 min, >98% Purity.

***N*-(3-ethynyl-4-fluorophenyl)-6-iodoquinazolin-4-amine** (**97**) was obtained as a tan solid (116 mg, 0.298 mmol, 58 %). MP 282-284 °C; ^1^H NMR (400 MHz, DMSO-*d*_6_) δ 11.70 (s, 1H), 9.35 (d, *J* = 1.8 Hz, 1H), 8.97 (s, 1H), 8.37 (dd, *J* = 8.7, 1.7 Hz, 1H), 7.97 (dd, *J* = 6.4, 2.7 Hz, 1H), 7.82 (ddd, *J* = 9.0, 4.7, 2.7 Hz, 1H), 7.76 (d, *J* = 8.8 Hz, 1H), 7.45 (t, *J* = 9.1 Hz, 1H), 4.61 (s, 1H). ^13^C NMR (101 MHz, DMSO-*d*_6_) δ 160.33 (d, *J* = 250.2 Hz), 158.5, 151.3, 144.2, 138.6, 133.1 - 133.0 (m, *J* = 3.1 Hz, 2C), 129.5 (d, *J* = 1.5 Hz), 127.4 (d, *J* = 8.4 Hz), 121.9, 116.2 (d, *J* = 22.2 Hz), 115.2, 110.1 (d, *J* = 17.1 Hz), 94.2, 86.9 (d, *J* = 3.3 Hz), 76.2. HRMS *m/z* [M+H]^+^ calcd for C_16_H_10_N_3_FI: 389.9904, found 389.9892, LC *t*_R_ = 4.66 min, >98% Purity.

***N*-(4-chloro-3-ethynylphenyl)-6-iodoquinazolin-4-amine** (**98**) was obtained as a tan solid (194 mg, 0.478 mmol, 93 %). MP 266-268 °C; ^1^H NMR (400 MHz, DMSO-*d*_6_) δ 11.70 (s, 1H), 9.37 (d, *J* = 1.8 Hz, 1H), 9.00 (s, 1H), 8.37 (dd, *J* = 8.7, 1.7 Hz, 1H), 8.08 (d, *J* = 2.5 Hz, 1H), 7.86 (dd, *J* = 8.8, 2.6 Hz, 1H), 7.77 (d, *J* = 8.7 Hz, 1H), 7.68 (d, *J* = 8.8 Hz, 1H), 4.68 (s, 1H). ^13^C NMR (101 MHz, DMSO-*d*_6_) δ 158.4, 151.4, 144.2, 139.1, 135.9, 133.0, 132.1, 129.7, 129.0, 126.2, 122.3, 121.5, 115.4, 94.3, 86.7, 79.6. HRMS *m/z* [M+H]^+^ calcd for C_16_H_10_N_3_ClI: 405.9608, found 405.9597, LC *t*_R_ = 5.37 min, >98% Purity.

***N*-(5-ethynyl-2-fluorophenyl)-6,7-dimethoxyquinazolin-4-amine** (**99**) was obtained as a yellow solid (200 mg, 0.619 mmol, 93 %). MP 260-262 °C; ^1^H NMR (400 MHz, DMSO-*d*_6_) δ 11.87 (s, 1H), 8.83 (s, 1H), 8.43 (s, 1H), 7.69 (dd, *J* = 7.2, 2.1 Hz, 1H), 7.56 (ddd, *J* = 8.5, 4.7, 2.2 Hz, 1H), 7.47 – 7.40 (m, 2H), 4.28 (s, 1H), 4.02 (s, 3H), 4.00 (s, 3H). ^13^C NMR (101 MHz, DMSO-*d*_6_) δ 159.0, 157.1 (d, *J* = 252.5 Hz), 156.6, 150.4, 148.7, 135.7, 132.5 (d, *J* = 8.4 Hz), 132.0 (d, *J* = 1.8 Hz), 125.0 (d, *J* = 13.5 Hz), 118.4 (d, *J* = 3.8 Hz), 117.0 (d, *J* = 21.1 Hz), 107.0, 104.2, 99.6, 81.8, 81.3, 57.0, 56.5. HRMS *m/z* [M+H]^+^ calcd for C_18_H_15_N_3_O_2_F: 324.1148, found 324.1138, LC *t*_R_ = 3.56 min, >98% Purity.

***N*-(3-ethynyl-2-fluorophenyl)-6,7-dimethoxyquinazolin-4-amine** (**100**) was obtained as a colourless solid (178 mg, 0.551 mmol, 82 %). MP 244-246 °C; ^1^H NMR (400 MHz, DMSO-*d*_6_) δ 11.87 (s, 1H), 8.83 (s, 1H), 8.42 (s, 1H), 7.68 – 7.50 (m, 2H), 7.42 (s, 1H), 7.34 (td, *J* = 7.9, 0.8 Hz, 1H), 4.61 (s, 1H), 4.02 (s, 3H), 4.00 (s, 3H). ^13^C NMR (101 MHz, DMSO-*d*_6_) δ 159.0, 157.49 (d, *J* = 254.4 Hz), 156.6, 150.4, 148.8, 135.7, 132.5, 129.7, 125.0 (d, *J* = 12.0 Hz), 124.8 (d, *J* = 4.6 Hz), 111.0 (d, *J* = 14.6 Hz), 107.0, 104.2, 99.7, 87.0 (d, *J* = 3.7 Hz), 76.3, 57.0, 56.5. HRMS *m/z* [M+H]^+^ calcd for C_18_H_15_N_3_O_2_F: 324.1148, found 324.1137, LC *t*_R_ = 3.59 min, >98% Purity.

***N*-(3-ethynyl-4-fluorophenyl)-6,7-dimethoxyquinazolin-4-amine** (**101**) was obtained as a yellow solid (197 mg, 0.609 mmol, 91 %). MP 260-262 °C; ^1^H NMR (400 MHz, DMSO-*d*_6_) δ 11.32 (s, 1H), 8.78 (s, 1H), 8.33 (s, 1H), 7.96 (dd, *J* = 6.4, 2.7 Hz, 1H), 7.84 (ddd, *J* = 9.0, 4.7, 2.7 Hz, 1H), 7.40 (t, *J* = 9.1 Hz, 1H), 7.33 (s, 1H), 4.58 (s, 1H), 4.01 (s, 3H), 3.97 (s, 3H). ^13^C NMR (101 MHz, DMSO-*d*_6_) δ 159.8 (d, *J* = 249.1 Hz), 157.8, 156.0, 150.0, 149.3, 137.4, 133.8 (d, *J* = 3.3 Hz), 129.2, 127.2 (d, *J* = 8.2 Hz), 115.9 (d, *J* = 22.1 Hz), 109.9 (d, *J* = 16.8 Hz), 107.5, 103.8, 100.9, 86.6 (d, *J* = 3.4 Hz), 76.4, 57.0, 56.4. HRMS *m/z* [M+H]^+^ calcd for C_18_H_15_N_3_O_2_F: 324.1148, found 324.1138, LC *t*_R_ = 3.66 min, >98% Purity.

***N*-(4-chloro-3-ethynylphenyl)-6,7-dimethoxyquinazolin-4-amine** (**102**) was obtained as a yellow solid (216 mg, 0.636 mmol, 95 %). MP 261-263 °C; ^1^H NMR (400 MHz, DMSO-*d*_6_) δ 11.28 (s, 1H), 9.19 (d, *J* = 2.0 Hz, 1H), 8.63 (d, *J* = 6.9 Hz, 1H), 8.19 (dd, *J* = 9.0, 2.0 Hz, 1H), 8.12 (d, *J* = 9.0 Hz, 1H), 7.66 – 7.50 (m, 2H), 7.06 (d, *J* = 6.9 Hz, 1H). ^13^C NMR (101 MHz, DMSO-*d*_6_) δ 159.8 (d, *J* = 249.1 Hz), 157.8, 156.0, 150.0, 149.3, 137.4, 133.8 (d, *J* = 3.3 Hz), 129.2, 127.2 (d, *J* = 8.2 Hz), 115.9 (d, *J* = 22.1 Hz), 109.9 (d, *J* = 16.8 Hz), 107.5, 103.8, 100.9, 86.2 (d, *J* = 3.4 Hz), 76.4, 57.0, 56.4. HRMS *m/z* [M+H]^+^ calcd for C_18_H_15_N_3_O_2_Cl: 340.0853, found 340.0842, LC *t*_R_ = 4.08 min, >98% Purity.

**6-bromo-*N*-(3,4,5-trifluorophenyl)quinolin-4-amine** (**103**) was obtained as a colourless solid (185 mg, 0.524 mmol, 85 %). MP >300 °C; ^1^H NMR (400 MHz, DMSO-*d*_6_) δ 11.28 (s, 1H), 9.19 (d, *J* = 2.0 Hz, 1H), 8.63 (d, *J* = 6.9 Hz, 1H), 8.19 (dd, *J* = 9.0, 2.0 Hz, 1H), 8.12 (d, *J* = 9.0 Hz, 1H), 7.66 – 7.50 (m, 2H), 7.06 (d, *J* = 6.9 Hz, 1H). ^13^C NMR (101 MHz, DMSO-*d*_6_) δ 153.8, 151.9 (dd, *J* = 10.2, 5.2 Hz), 149.4 (dd, *J* = 10.3, 5.1 Hz), 143.4, 139.0 (t, *J* = 15.5 Hz), 137.3, 136.7, 136.5 (t, *J* = 15.6 Hz), 133.6 (td, *J* = 11.4, 4.4 Hz), 126.3, 122.6, 120.2, 118.8, 110.8 −110.6 (m, 1C), 101.4. HRMS *m/z* [M+H]^+^ calcd for C_15_H_9_BrF_3_N_2_: 352.9901, found 352.9892, LC *t*_R_ = 3.89 min, >98% Purity.

**6-bromo-*N*-(3,4-difluorophenyl)quinolin-4-amine** (**104**) was obtained as a yellow solid (185 mg, mmol, 89 %). MP >300 °C; ^1^H NMR (400 MHz, DMSO-*d*_6_) δ 11.15 (s, 1H), 9.15 (d, *J* = 2.0 Hz, 1H), 8.57 (d, *J* = 6.9 Hz, 1H), 8.19 (dd, *J* = 9.0, 2.0 Hz, 1H), 8.09 (d, *J* = 9.0 Hz, 1H), 7.75 – 7.56 (m, 2H), 7.37 (dddd, *J* = 8.8, 4.1, 2.6, 1.5 Hz, 1H), 6.91 (d, *J* = 6.9 Hz, 1H). ^13^C NMR (101 MHz, DMSO-*d*_6_) δ 154.1, 151.2 – 149.3 (m, 1C), 148.7 – 146.8 (m, 1C), 148.5 (d, *J* = 13.6 Hz, 1C), 147.1 (d, *J* = 12.6 Hz, 1C), 0.043.3, 137.3, 136.7, 134.0 (dd, *J* = 8.4, 3.4 Hz), 126.2, 122.7 (dd, *J* = 6.7, 3.3 Hz, 1C), 122. 6, 120.0, 118.7 (d, *J* = 18.1 Hz, 1C), 118.6, 115.2 (d, *J* = 19.0 Hz, 1C), 100.8. HRMS *m/z* [M+H]^+^ calcd for C_15_H_10_N_2_F_2_Br: 334.9995, found 334.9987, LC *t*_R_ = 3.68 min, >98% Purity.

**6-bromo-*N*-(3,5-difluorophenyl)quinolin-4-amine** (**105**) was obtained as a colourless solid (164 mg, 0.489 mmol, 79 %). MP >300 °C; ^1^H NMR (400 MHz, DMSO-*d*_6_) δ 11.28 (s, 1H), 9.20 (d, *J* = 2.0 Hz, 1H), 8.62 (d, *J* = 6.9 Hz, 1H), 8.19 (dd, *J* = 9.0, 2.0 Hz, 1H), 8.12 (d, *J* = 9.0 Hz, 1H), 7.31 (ddt, *J* = 11.2, 7.1, 2.0 Hz, 3H), 7.13 (d, *J* = 6.9 Hz, 1H). ^13^C NMR (101 MHz, DMSO-*d*_6_) δ 164.1 (d, *J* = 15.0 Hz), 161.6 (d, *J* = 15.0 Hz), 153.4, 143.5, 140.1(t, *J* = 13.0 Hz), 139.9, 137.4, 136.7, 126.3, 122.6, 120.2, 119.0, 108.5 - 108.3 (m, 1C), 102.6 (t, *J* = 26.1 Hz), 101.7. HRMS *m/z* [M+H]^+^ calcd for C_15_H_10_N_2_F_2_Br: 334.9995, found 334.9987, LC *t*_R_ = 3.72 min, >98% Purity.

**6-bromo-*N*-(2,5-difluorophenyl)quinolin-4-amine** (**106**) was obtained as a colourless solid (125 mg, 0.373 mmol, 60 %). MP 187-189 °C; ^1^H NMR (400 MHz, DMSO-*d*_6_) δ 11.11 (s, 1H), 9.21 (d, *J* = 2.0 Hz, 1H), 8.59 (d, *J* = 6.9 Hz, 1H), 8.20 (dd, *J* = 9.1, 1.9 Hz, 1H), 8.12 (d, *J* = 9.0 Hz, 1H), 7.78 – 7.46 (m, 2H), 7.38 – 7.25 (m, 1H), 6.56 (dd, *J* = 6.9, 2.3 Hz, 1H). ^13^C NMR (101 MHz, DMSO-*d*_6_) δ 161.5 (dd, *J* = 247.9, 11.7 Hz), 157.1 (dd, *J* = 251.6, 13.2 Hz), 154.5, 143.4, 137.3, 136.73, 130.24 (dd, *J* = 10.2, 2.0 Hz), 126.21, 122.66, 121.00 (dd, *J* = 12.6, 3.9 Hz), 120.18, 118.35, 113.0 (dd, *J* = 22.6, 3.7 Hz), 105.8 (dd, *J* = 27.1, 24.0 Hz), 100.9 (d, *J* = 1.8 Hz). HRMS *m/z* [M+H]^+^ calcd for C_15_H_10_N_2_F_2_Br: 334.9995, found 334.9985, LC *t*_R_ = 3.65 min, >98% Purity.

**6-bromo-*N*-(2,4-difluorophenyl)quinolin-4-amine** (**107**) was obtained as a colourless solid (94 mg, 0.281 mmol, 45 %). MP 301-303 °C; ^1^H NMR (400 MHz, DMSO-*d*_6_) δ 11.18 (s, 1H), 9.20 (d, *J* = 2.0 Hz, 1H), 8.62 (d, *J* = 6.8 Hz, 1H), 8.20 (dd, *J* = 9.0, 2.0 Hz, 1H), 8.12 (d, *J* = 9.0 Hz, 1H), 7.65 – 7.48 (m, 2H), 7.40 (ddt, *J* = 9.2, 8.0, 3.5 Hz, 1H), 6.68 (dd, *J* = 6.8, 2.7 Hz, 1H). ^13^C NMR (101 MHz, DMSO-*d*_6_) δ 158.2 (dd, *J* = 242.3, 2.3 Hz), 153.2 (dd, *J* = 245.2, 2.9 Hz), 153.9, 143.6, 137.4, 136.7, 126.2, 125.6 (dd, *J* = 14.7, 11.0 Hz), 122.8, 120.3, 118.5, 118.4 (dd, *J* = 22.5, 9.7 Hz), 116.3 (dd, *J* = 23.9, 8.2 Hz), 115.5, 115.2, 101.5 (d, *J* = 2.3 Hz). HRMS *m/z* [M+H]^+^ calcd for C_15_H_10_N_2_F_2_Br: 334.9995, found 334.9985, LC *t*_R_ = 3.63 min, >98% Purity.

**6-bromo-*N*-(4-fluorophenyl)quinolin-4-amine** (**108**) was obtained as a yellow solid (181 mg, 0.571 mmol, 92 %). MP 175-177 °C; ^1^H NMR (400 MHz, DMSO-*d*_6_) δ 11.12 (s, 1H), 9.17 (d, *J* = 2.1 Hz, 1H), 8.52 (d, *J* = 7.0 Hz, 1H), 8.17 (dd, *J* = 9.0, 2.0 Hz, 1H), 8.08 (d, *J* = 9.0 Hz, 1H), 7.62 – 7.48 (m, 2H), 7.48 – 7.35 (m, 2H), 6.75 (d, *J* = 6.9 Hz, 1H). ^13^C NMR (101 MHz, DMSO-*d*_6_) δ 160.8 (d, *J* = 244.6 Hz), 154.3, 143.0, 137.2, 136.6, 133.3 (d, *J* = 2.9 Hz), 127.8 (d, *J* = 8.7 Hz, 2C), 126.2, 122.4, 119.9, 118.5, 116.8 (d, *J* = 22.9 Hz, 2C), 100.3. HRMS *m/z* [M+H]^+^ calcd for C_15_H_11_N_2_FBr: 317.0090, found 317.0080, LC *t*_R_ = 3.57 min, >98% Purity.

**6-bromo-*N*-(4-fluorophenyl)quinazolin-4-amine** (**109**) was obtained as a yellow solid (126 mg, 0.396 mmol, 64 %). MP 257-259 °C; ^1^H NMR (400 MHz, DMSO-*d*_6_) δ 10.99 (s, 1H), 9.10 (d, *J* = 2.1 Hz, 1H), 8.77 (s, 1H), 8.11 (dd, *J* = 8.9, 2.1 Hz, 1H), 8.03 – 7.46 (m, 3H), 7.49 – 7.00 (m, 2H). ^13^C NMR (101 MHz, DMSO-*d*_6_) δ 160.7, 158.2, 157.9, 152.9, 142.9, 137.5, 133.9, 133.9, 126.5, 125.8, 125.7 (d, *J* = 8.2 Hz), 119.9, 115.63 115.3 (d, *J* = 22.5 Hz). HRMS *m/z* [M+H]^+^ calcd for C_14_H_10_N_3_FBr: 318.0042, found 318.0032, LC *t*_R_ = 3.71 min, >98% Purity.

**6-bromo-*N*-(3-fluorophenyl)quinolin-4-amine** (**110**) was obtained as a yellow solid (182 mg, 0.574 mmol, 93 %).) MP 187-189 °C; ^1^H NMR (400 MHz, DMSO-*d*_6_) δ 11.19 (s, 1H), 9.19 (d, *J* = 2.0 Hz, 1H), 8.57 (d, *J* = 6.9 Hz, 1H), 8.18 (dd, *J* = 9.0, 2.0 Hz, 1H), 8.10 (d, *J* = 9.0 Hz, 1H), 7.61 (td, *J* = 8.2, 6.5 Hz, 1H), 7.42 (dt, *J* = 10.1, 2.2 Hz, 1H), 7.36 (ddd, *J* = 8.0, 2.0, 0.9 Hz, 1H), 7.27 (tdd, *J* = 8.6, 2.6, 0.9 Hz, 1H), 6.97 (d, *J* = 6.9 Hz, 1H). ^13^C NMR (101 MHz, DMSO-*d*_6_) δ 162.5 (d, *J* = 245.0 Hz), 153.7, 143.3, 138.9 (d, *J* = 10.3 Hz), 137.4, 136.7, 131.6 (d, *J* = 9.3 Hz), 126.3, 122.5, 121.2 (d, *J* = 3.0 Hz), 120.0, 118.8, 114.2 (d, *J* = 21.0 Hz), 112.3 (d, *J* = 23.9 Hz), 100.9. HRMS *m/z* [M+H]^+^ calcd for C_15_H_11_N_2_FBr: 317.0090, found 317.0081, LC *t*_R_ = 3.60 min, >98% Purity.

**6-bromo-*N*-(2-fluorophenyl)quinolin-4-amine** (**111**) was obtained as a yellow solid (155 mg, 0.489 mmol, 79 %). MP 183-185 °C; ^1^H NMR (400 MHz, DMSO-*d*_6_) δ 11.16 (s, 1H), 9.23 (d, *J* = 2.0 Hz, 1H), 8.58 (d, *J* = 6.9 Hz, 1H), 8.20 (dd, *J* = 9.0, 2.0 Hz, 1H), 8.12 (d, *J* = 9.0 Hz, 1H), 7.59 (td, *J* = 7.9, 1.6 Hz, 1H), 7.56 – 7.47 (m, 2H), 7.45 – 7.38 (m, 1H), 6.54 (dd, *J* = 6.9, 2.6 Hz, 1H). ^13^C NMR (101 MHz, DMSO-*d*_6_) δ 156.8 (d, *J* = 249.1 Hz), 154.3, 143.3, 137.2, 136.7, 130.1 (d, *J* = 8.0 Hz), 128.8, 126.2, 125.8 (d, *J* = 3.7 Hz), 124.3 (d, *J* = 12.2 Hz), 122.6, 120.2, 118.4, 117.1 (d, *J* = 19.3 Hz), 100.9 (d, *J* = 2.1 Hz). HRMS *m/z* [M+H]^+^ calcd for C_15_H_11_N_2_BrF: 317.0090, found 317.0084, LC *t*_R_ = 3.44 min, >98% Purity.

**6-bromo-*N*-(3-bromophenyl)quinolin-4-amine** (**112**) was obtained as a yellow solid (204 mg, 0.540 mmol, 87 %). MP 262-264 °C; ^1^H NMR (400 MHz, DMSO-*d*_6_) δ 11.26 (s, 1H), 9.22 (d, *J* = 2.0 Hz, 1H), 8.56 (d, *J* = 6.9 Hz, 1H), 8.17 (dd, *J* = 9.0, 1.9 Hz, 1H), 8.11 (d, *J* = 9.0 Hz, 1H), 7.77 – 7.70 (m, 1H), 7.61 (ddd, *J* = 6.2, 2.7, 1.9 Hz, 1H), 7.57 – 7.49 (m, 2H), 6.92 (d, *J* = 6.9 Hz, 1H). ^13^C NMR (101 MHz, DMSO-*d*_6_) δ 154.2, 143.7, 139.0, 137.8, 137.1, 132.2, 130.6, 128.3, 126.8, 124.6, 122.9, 122.7, 120.5, 119.2, 101.3. HRMS *m/z* [M+H]^+^ calcd for C_15_H_11_N_2_Br_2_: 376.9289, found 376.9287, LC *t*_R_ = 3.96 min, >98% Purity.

**6-bromo-*N*-(3-bromophenyl)quinazolin-4-amine** (**113**) was obtained as a colourless solid (161 mg, 0.421 mmol, 69 %). MP 221-223 °C; ^1^H NMR (400 MHz, DMSO-*d*_6_) δ 11.71 (s, 1H), 9.31 – 9.23 (m, 1H), 8.99 (t, *J* = 0.8 Hz, 1H), 8.24 (dd, *J* = 8.9, 2.0 Hz, 1H), 8.08 (t, *J* = 1.9 Hz, 1H), 7.97 – 7.90 (m, 1H), 7.81 (ddd, *J* = 7.9, 2.1, 1.1 Hz, 1H), 7.52 (ddd, *J* = 8.0, 1.9, 1.1 Hz, 1H), 7.46 (t, *J* = 8.0 Hz, 1H). ^13^C NMR (101 MHz, DMSO-*d*_6_) δ 158.7, 151.6, 139.1, 138.8, 138.4, 137.3, 130.7, 129.0, 128.0, 127.2, 126.8, 123.2, 122.8, 121.2, 120.9, 115.3. HRMS *m/z* [M+H]^+^ calcd for C_15_H_9_Br_2_N_3_: 379.9398, found 379.9209, LC *t*_R_ 4.96 min, >98% Purity.

## Acknowledgments

The SGC is a registered charity (number 1097737) that receives funds from AbbVie, Bayer Pharma AG, Boehringer Ingelheim, Canada Foundation for Innovation, Eshelman Institute for Innovation, Genome Canada, Innovative Medi-cines Initiative (EU/EFPIA) [ULTRA-DD grant no. 115766], Janssen, Merck KGaA Darmstadt Germany, MSD, Novartis Pharma AG, Ontario Ministry of Economic Development and Innovation, Pfizer, São Paulo Research Foundation-FAPESP, Takeda, and Wellcome [106169/ZZ14/Z]. We thank Biocenter Finland/DDCB for financial support and CSC - IT Center for Science Ltd. Finland for the use of their facilities, software licenses and computational resources. We are grateful Dr. Brandie Ehrmann for LC-MS/HRMS support provided by the Mass Spectrometry Core Laboratory at the University of North Carolina at Chapel Hill. We also thank the EPSRC UK National Crystallography Service for funding and collection of the crystallographic data for (**7**, **11**, **15**-**20**, **22**-**25**, **27**-**29**, **31**-**34**, **36**-**37**, **39**-**40**, **43**-**46**, **50**, **53, 67**).

## AUTHOR INFORMATION

### Author Contributions

The manuscript was written through contributions of all authors. All authors approved of the final version of the manuscript.

### Notes

The authors declare no competing financial interest.

